# Amino acid auxotrophy is a feature of a distinct bacterial lifestyle across environments

**DOI:** 10.64898/2026.06.03.729626

**Authors:** Alyssa Henderson, Marco Gabrielli, Rachel Szabo, A. Murat Eren, Martin Ackermann, Olga T. Schubert

## Abstract

Amino acid auxotrophy, the loss of biosynthetic potential for an amino acid, is a highly prevalent feature of microbial life, yet the evolutionary processes driving its distribution in nature remain unclear. Here, we sought to determine whether auxotrophy emerges as an independent response to nutrient availability or as part of a broader transition in genomic lifestyle. By first predicting auxotrophies through genome-scale metabolic models and flux-balance analysis, and subsequently applying phylogenetically informed statistical frameworks, we tested the predictions of the Black Queen Hypothesis, which posits that auxotrophy evolves as a dynamic response to metabolite availability, and disentangled the signal of convergent environmental selection from the background of shared evolutionary history. This approach reveals that phylogeny is the dominant predictor of auxotrophy, vastly outweighing environmental context. Beyond correcting for ancestry, our analysis uncovered a previously obscured signal of consistent, co-occurring genomic changes: auxotrophies do not occur as independent losses, but rather as part of a broader reorganization of the genome. We show that auxotrophies co-occur significantly more often than predicted by chance and that the probability of loss scales with the number of other auxotrophies. These findings reveal auxotrophy as a feature of a conserved, phylogenetically entrenched lifestyle rather than a transient response to nutrient availability — inconsistent with the Black Queen Hypothesis and instead aligning with genome streamlining as the overarching driver of metabolic dependency in natural microbial systems.

## Introduction

Auxotrophy, the inability of an organism to synthesize one or more of the amino acids, vitamins, and nucleotides required for its own growth, is a widespread feature of bacterial life, documented across phylogenetically diverse lineages recovered from environments ranging from open ocean surface waters to animal-associated communities (D’Souza et al., 2014; Ramoneda et al. 2023). Its opposite, prototrophy, describes organisms that retain full biosynthetic self-sufficiency. Under laboratory conditions, amino acid auxotrophies evolve readily — experimental populations of *Escherichia coli* supplied with amino acids at high concentrations can evolve auxotrophy within a few thousand generations, with auxotrophic strains outcompeting their prototrophic ancestors (D’Souza and Kost 2016). However, how auxotrophy evolves under the far more dilute and temporally variable amino acid concentrations that characterize most natural environments remains an open question (Jørgensen 1987; Amaral et al. 2024; Niu et al. 2023; Fasching et al. 2020; Ramesh et al. 2026).

Three related but distinct conceptual frameworks have been invoked to explain why microorganisms lose biosynthetic capabilities. The simplest is a cost-benefit argument: amino acid biosynthesis carries measurable metabolic and energetic costs (Kaleta et al. 2013; Akashi and Gojobori 2002), and when the amino acid is reliably available from the surrounding environment, selection may favor strains that shed the biosynthetic burden and reallocate those cellular resources toward growth (D’Souza et al. 2014). If auxotrophy is driven solely by independent cost-benefit trade-offs, the loss of biosynthetic capabilities would be dictated only by the specific metabolites available in a given niche.

As an extension of cost-benefit analysis, the Black Queen Hypothesis (BQH) invokes selection in the context of communities that lack an external supply of amino acids (Morris et al. 2012). In this framework, metabolites are leaked from the cells that synthesise them, such that non-producers can benefit without paying biosynthetic costs. The BQH predicts polymorphic behavior within communities, where a biosynthetic pathway is maintained at intermediate frequencies as some members grow on molecules leaked by others, rather than universal loss across entire clades of bacteria (Morris 2015). Furthermore, because the BQH depends on metabolite leakage, it is most relevant to metabolites that are released readily from cells — a condition that may not be generally satisfied by amino acids, whose biosynthesis and leakage in natural environments is highly variable and context-dependent (McKinlay 2023; Ramesh et al. 2026; Sulheim et al. 2025).

A third framework, genome streamlining theory, invokes directional selection for genome economy in obligate host-associated microbes or in populations characterized by large effective population sizes and stable nutrient regimes (Giovannoni et al. 2014). Streamlined organisms shed biosynthetic pathways and non-essential functions, including regulatory networks, signal transduction, and secondary metabolism, yielding reduced genome size, elevated coding density, and relative enrichment of core translational machinery (Giovannoni et al. 2014; Lynch 2006). Under streamlining, amino acid auxotrophies are not independent events tracking individual metabolite availability, but consequences of a broader lifestyle transition. This framework predicts auxotrophies as phylogenetically entrenched, co-occurring with one another, and embedded in a recognizable reorganization of genomic functional content towards translational and core cellular functions and away from regulatory complexity, metabolic versatility, and biosynthetic breadth.

These three frameworks make fundamentally different predictions about the evolutionary drivers and the extant distribution of auxotrophies. Cost-benefit logic predicts auxotrophy wherever a metabolite is available, largely independent of phylogeny or other genomic changes. BQH predicts community-dependent, polymorphic losses rather than universal loss across clades. Genome streamlining predicts systematic, phylogenetically entrenched losses co-occurring with a broader shift in genomic content. Distinguishing among them therefore requires asking whether auxotrophies co-occur beyond chance, whether they are accompanied by a consistent functional reorganisation of the genome, and whether their distribution is structured primarily by evolutionary history or by local environmental conditions.

Previous large-scale comparative analyses have characterized the taxonomic and habitat distribution of amino acid auxotrophies, finding higher prevalence in host-associated environments and in taxa bearing streamlined genomes (Ramoneda et al. 2023; D’Souza et al. 2014). These analyses have largely been conducted without taking into account phylogenetic relationships. However, because species are not independent samples due to their shared evolutionary history (Felsenstein 1985), associations between environment and auxotrophy may not only reflect convergent adaptations to environmental characteristics but also the shared ancestry of clades that happen to occupy similar habitats. Phylogenetic comparative methods are well established for resolving this ambiguity (Martiny et al. 2013; Amend et al. 2016; Berlemont and Martiny 2013; Peay et al. 2011; Ives and Garland 2014; Felsenstein 1985; Li et al. 2020), but are not yet consistently applied to bacterial gene-content evolution at scale.

A further challenge for comparative analyses is that genomic databases have historically been dominated by genomes recovered from cultivated organisms, and cultivation imposes a powerful selective filter. Auxotrophic bacteria that require specific exogenous metabolites for growth are systematically underrepresented in culture collections, producing an isolation bias that distorts estimates of auxotrophy prevalence (Ramoneda et al. 2023; Pacheco-Valenciana et al. 2025). Accordingly, comparative analyses relying heavily on isolate genomes likely underestimate the prevalence of auxotrophy. Metagenome-assembled genomes (MAGs) circumvent this filter by recovering genomic diversity from environmental samples directly (Pacheco-Valenciana et al. 2025; Ramoneda et al. 2023).

Here, we use genome-scale metabolic models and flux-balance analysis to predict amino acid auxotrophies across thousands of high-quality MAGs. We apply phylogenetically corrected statistical models to disentangle the contributions of environment, gene content, and shared evolutionary history in driving observed auxotrophy patterns. We ask whether environmental associations are robust to phylogenetic correction, whether auxotrophic genomes carry a functional signature extending beyond the lost pathways themselves, and whether individual amino acid auxotrophies accumulate independently or co-occur as part of a distinct lifestyle.

Our findings reveal that amino acid auxotrophy exhibits a strong and pervasive phylogenetic signal, with evolutionary history accounting for substantial variation in auxotrophic status across the bacterial tree. Broad environmental context, while significant, explains considerably less variation, and its apparent signal is largely absorbed by genomic functional composition once both are considered together. Auxotrophic genomes carry a consistent and predictable functional signature, and individual amino acid auxotrophies co-occur far more frequently than expected by chance, scaling hierarchically with the total number of auxotrophies in a genome. Together, these results suggest that auxotrophy represents a phylogenetically structured genomic lifestyle — with coordinated pathway losses embedded within a broader functional reorganisation of the genome — rather than a purely environment-driven response to nutrient availability.

## Results and Discussion

### 1. Benchmarking of auxotrophy prediction from genome-scale models

To examine the drivers of auxotrophy across the bacterial domain, we constructed genome-scale metabolic models (GEMs) of representative genomes and predicted amino acid auxotrophies using flux balance analysis (Figure 1a). Given previous reports regarding the limitations of utilizing GEMs to predict metabolic features (Price 2023), we benchmarked the accuracy of our approach on a selection of 223 genomes collected from bacteria with published phenotyping data that were compiled by previous studies (Price et al. 2020; Ramoneda et al. 2023; Starke et al. 2023). We derived a sensitivity of 78%, which measures the model’s ability to predict true auxotrophs, and a specificity of 98%, which measures the model’s ability to correctly identify non-auxotrophs. The overall proportion of correct identifications, or accuracy, is 97%, though this metric is skewed by the prevalence of prototrophic observations in the benchmarking dataset (see Methods).

**Figure 1:**
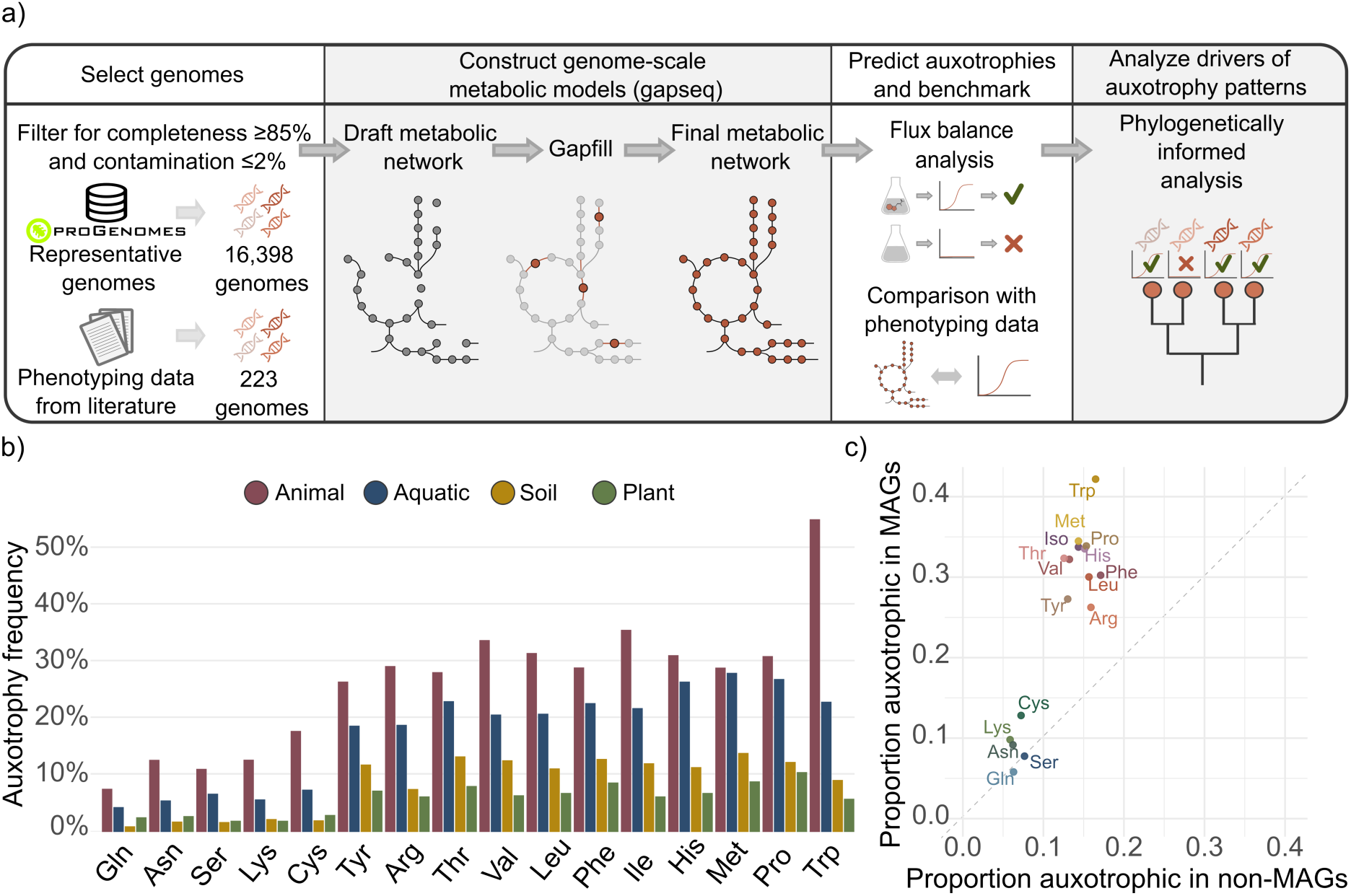
Methodological workflow and resulting distribution of microbial auxotrophies. a) Workflow for generating a database of auxotrophy predictions. b) Frequency of auxotrophy for each amino acid across genomes and environments. c) Comparison of auxotrophy frequency in MAGs and non-MAGs, which were mostly derived from isolates.

While the auxotrophy predictions align well with phenotyping data, GEMs rely on gene presence-absence data and gapfilling algorithms, which can miss auxotrophies caused by inactive gene expression (Kriel et al. 2013), inactivating point mutations rather than full gene deletions, or small losses in biosynthetic pathways that gapfilling algorithms can erroneously restore. While auxotrophies due to point mutations and single-gene losses are theoretically possible, they are likely often transient because selection on the remaining genes on the biosynthetic pathway is relaxed, rendering them susceptible to mutation accumulation and gene decay (Lerat and Ochman 2004; Dagan et al. 2006; Kinjo et al. 2021). Over evolutionary timescales, these non-functional pathways are purged from the genome. Therefore, the presence of a near-complete biosynthetic gene set is likely often an indicator of a true prototrophy rather than a transient state. Based on our models’ sensitivity of 78%, as well as the above-listed sources of erroneous prototrophy predictions, we consider GEM inferences as conservative estimates of auxotrophy.

### 2. Prediction of auxotrophy in a large set of species-representative genomes

Having benchmarked the accuracy of the auxotrophy predictions using phenotyped strains, we then employed this method to infer auxotrophy in genomes across the bacterial tree in the proGenomes3 database (Fullam et al. 2023). This database includes genomic sequences that are assigned to species-specific identifier clusters, each of which includes a representative genome (Fullam et al. 2023). We selected representatives that included environmental metadata and constructed GEMs for each of them, yielding functional models. To predict auxotrophy, we performed flux balance analysis on each model in simulated growth media with and without each amino acid (Starke et al. 2023; Zimmermann et al. 2021). We maintained 16,398 high-quality genomes with functional GEMs that yielded auxotrophy predictions. The examined genomes included both MAGs (n=8249) and genomes not described as MAGs in the proGenomes3 metadata (n=8149).

We found that auxotrophy rates vary substantially across amino acids and environments (Figure 1b). Predictions from the GEMs included auxotrophies for 17 of the 20 proteinogenic amino acids. No auxotrophy was predicted for alanine, aspartate, or glutamate, consistent with the benchmarking dataset (Supplementary Figure 3); these amino acids are both among the cheapest to synthesize and biosynthetically entangled with core nitrogen metabolism, making their loss selectively costly regardless of exogenous availability (Supplementary Discussion 1; Price et al. 2020; Akashi and Gojobori 2002). Glycine auxotrophy was predicted in only 1.7% of genomes and was excluded from further analysis.

Amino acids cluster into two groups, those rarely lost and those frequently lost, and this division maps onto the cost and dispensability of their biosynthetic pathways. Those with low auxotrophy rates — lysine, asparagine, glutamine, serine, and cysteine — share either low biosynthetic costs or deep entanglement with essential cellular processes: for instance, the lysine biosynthesis pathway doubles as the route to peptidoglycan synthesis, glutamine synthetase is a central regulatory node in nitrogen metabolism, and cysteine biosynthesis is coupled to sulfur assimilation (see Supplementary Discussion 2 for discussion of each amino acid) (Akashi and Gojobori 2002; Kaleta et al. 2013; Reitzer 2003, 2004; Yuan et al. 2009; Fernandez et al. 2025). In contrast, the remaining 11 amino acids with higher auxotrophy rates are synthesised via longer, more costly routes with fewer overlaps with essential pathways, making them more dispensable when the amino acid can be acquired from the environment. This holds even for amino acids with critical downstream roles: methionine is the precursor to S-adenosylmethionine, the universal methyl donor, yet shows high predicted auxotrophy rates, potentially because a cell can synthesise SAM from exogenously acquired methionine, meaning auxotrophy carries no inherent penalty provided methionine is available (Fontecave et al. 2004).

Notably, we determined that, for all 16 amino acids examined in this study, MAGs exhibit auxotrophy rates approximately two-fold higher than the remaining genomes, which are mostly derived from isolates (Figure 1c). To ensure this gap is biological rather than technical, we confirmed that higher auxotrophy rates in MAGs persist in near-complete genomes (98%–100% completeness) and that auxotrophy counts do not increase with genome fragmentation. We further validated gapseq on simulated MAGs, achieving sensitivity 0.895 and specificity 0.928 against experimental phenotypes in high-quality bins (≥85% completeness; Supplementary Figures 5, 6; Supplementary Table S1b; Supplementary Discussion 3). This discrepancy between MAGs and non-MAGs rather hints at potential isolation bias (Supplementary Figure 4) (Dewi Puspita et al. 2012; Hugenholtz 2002; Steen et al. 2019; Stewart 2012; Lewis et al. 2010). While cultivation media are typically supplemented with amino acids, an inability to isolate auxotrophs suggests that amino acid auxotrophy may co-occur with other properties hindering cultivation, such as slow growth or a need for complex cofactors, cross-fed intermediates, close associations with other organisms, or specific environmental conditions (Vartoukian 2016; Kapinusova et al. 2023; Kim et al. 2021). To capture this ecologically relevant diversity and avoid the bias inherent in isolate datasets, the remainder of our analysis was conducted exclusively using MAGs.

### 3. Phylogenetic history plays a larger role than the environment in shaping auxotrophy rates

To investigate whether auxotrophy emerges as an independent response to nutrient availability or as part of a broader transition in genomic lifestyle, we next examined how auxotrophies are distributed across environments.

Initial results indicate that, across amino acids, auxotrophy frequencies in animal-associated bacteria are greater than in other habitats, a result often explained as a consequence of the animal environment containing free amino acids (Figure 1b) (Yu et al. 2009; Ramoneda et al. 2023). However, these estimates based purely on comparing auxotrophy frequencies may be misleading when used to infer ecological or evolutionary drivers. A significant correlation between a niche and a trait may simply reflect phylogenetic inertia if the majority of species belong to a single, closely related clade. To identify a functional relationship, one must demonstrate that these traits co-occur across multiple, independent branches of the phylogenetic tree, suggesting a recurring response to selection.

We assessed the degree of evolutionary conservatism of amino acid auxotrophies by calculating the phylogenetic half-life for each amino acid — a metric representing the evolutionary distance at which a trait’s phylogenetic correlation decays by half (calculated on the latent scale; see Supplementary Methods S2). Nearly every amino acid biosynthetic pathway exhibited strong conservatism, with half-lives exceeding 5% of total tree depth (Figure 2c). Although the phylogeny is not time-calibrated, the bacterial radiation sampled spans approximately 2–2.5 billion years (Coleman et al. 2021; Davín et al. 2025), suggesting these half-lives correspond to hundreds of millions of years — timescales over which the phylogenetic correlation in auxotrophic state is maintained across lineages that diverged before the Cambrian. Serine was the sole exception, with a very low half-life, suggesting either rapid evolutionary turnover or technical artefacts in pathway annotation. Under a null model of random trait permutation, phylogenetic signal was effectively eliminated in all cases, confirming that the observed half-lives reflect genuine evolutionary structure rather than artefacts of tree topology or taxon sampling (Supplementary Figure 20). Consequently, these results demonstrate that microbial lineages cannot be treated as independent observations, since more related species are significantly more likely to share auxotrophy traits due to their shared evolutionary history.

**Figure 2:**
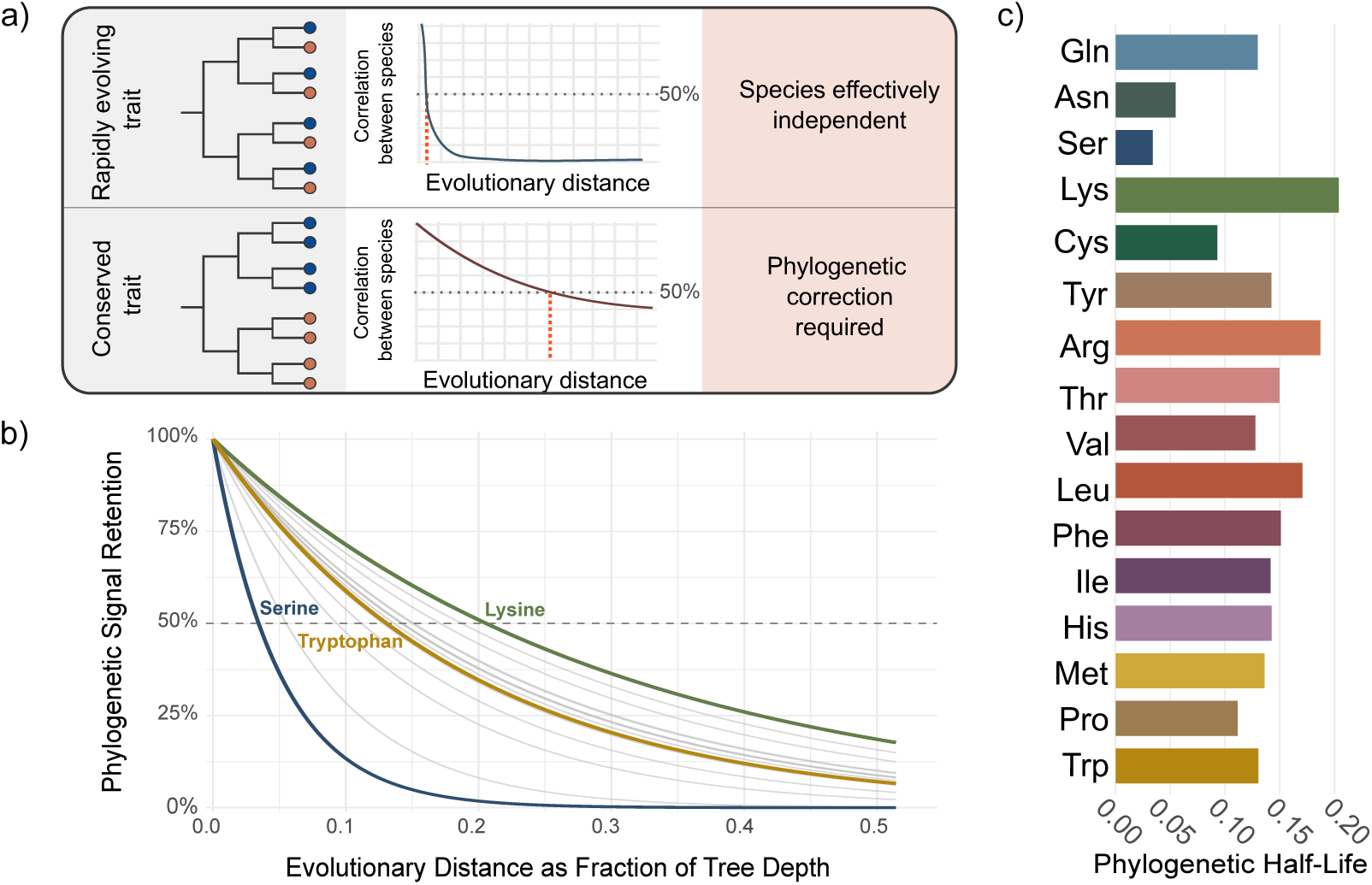
Auxotrophies exhibit varying but significant phylogenetic signal. a) Schematic illustrating the relationship between trait evolution rate and phylogenetic correlation decay. Rapidly evolving traits lose phylogenetic correlation quickly, while conserved traits maintain correlation across deep divergences. The phylogenetic half-life (orange line) marks the evolutionary distance at which correlation decays to 0.5. b) Phylogenetic signal decay curves for amino acid auxotrophy across the bacterial tree. Each curve shows the expected correlation in auxotrophic state between two lineages as a function of their phylogenetic distance, derived from phylogenetic logistic regression models. The phylogenetic half-life, which is the branch length at which correlation decays to 0.5, is indicated for each amino acid in grey, with three example curves highlighted in color. c) Phylogenetic half-lives shown here correspond to the points where the decay curves in b) cross the 0.5 correlation threshold. Bars are ordered by auxotrophy frequency, with rarest auxotrophies at the top. Higher values indicate greater evolutionary conservatism, reflecting auxotrophic states that are maintained across deeper divergence scales. Serine exhibits the lowest phylogenetic signal, which may be due to inaccurate assignments of serine auxotrophies due to gapfilling.

To disentangle the roles of phylogenetic history and environment in shaping auxotrophy, we compared models for standard logistic regression (GLM) and for phylogenetic logistic regression (PhyloGLM) of the environment predicting auxotrophy in the subset of MAGs assigned to soil, aquatic, animal-associated, or plant-associated categories (n=4,507). The GLM estimates the log-odds of auxotrophy based on observed phylogenetic tree tip frequencies, assuming that each bacterial genome represents an independent and identically distributed observation. In contrast, the PhyloGLM estimates the log-odds of auxotrophy by modeling the trait as a stochastic process evolving along the phylogeny. By incorporating a variance-covariance matrix derived from shared branch lengths, the model calculates the expected odds of auxotrophy for each condition while accounting for the non-independence of lineages.

For all but five of the 64 amino acid-environment pairs, phylogenetic correction resulted in a higher probability of auxotrophy across the entire dataset, with PhyloGLMs revealing a global upward shift in predicted odds compared to GLMs (Figure 3a). This indicates that once shared history is accounted for, the global propensity toward auxotrophy is higher than standard GLMs suggest. This global underestimation of auxotrophy by standard GLMs likely stems from a high degree of phylogenetic clustering among prototrophs, which masks the broader prevalence of metabolic dependency. Supporting this, prototrophs exhibited lower Mean Phylogenetic Distance than auxotrophs across nearly all amino acids (Supplementary Figure 21), and we observed a positive correlation (r = 0.665, p = 0.0102) between an amino acid’s phylogenetic half-life and the magnitude of the probability increase following correction (Supplementary Figures 7 and 11). This indicates that the GLMs’ underestimation is most severe for the most evolutionarily conserved traits, where phylogenetically clustered prototrophs are counted as independent observations despite their shared ancestry, disproportionately inflating the apparent prevalence of prototrophy. This clustering is likely reinforced by systematic biases: marker-gene-based completeness estimates can penalize streamlined, often auxotrophic genomes (Parks et al. 2015). Additionally, lower relative abundances of specialists can further hinder the assembly of auxotrophic MAGs (Ryback et al. 2022). By accounting for this redundancy in conserved lineages, the PhyloGLMs reveal a more accurate global baseline where auxotrophy is significantly more prevalent than previously recognized.

**Figure 3:**
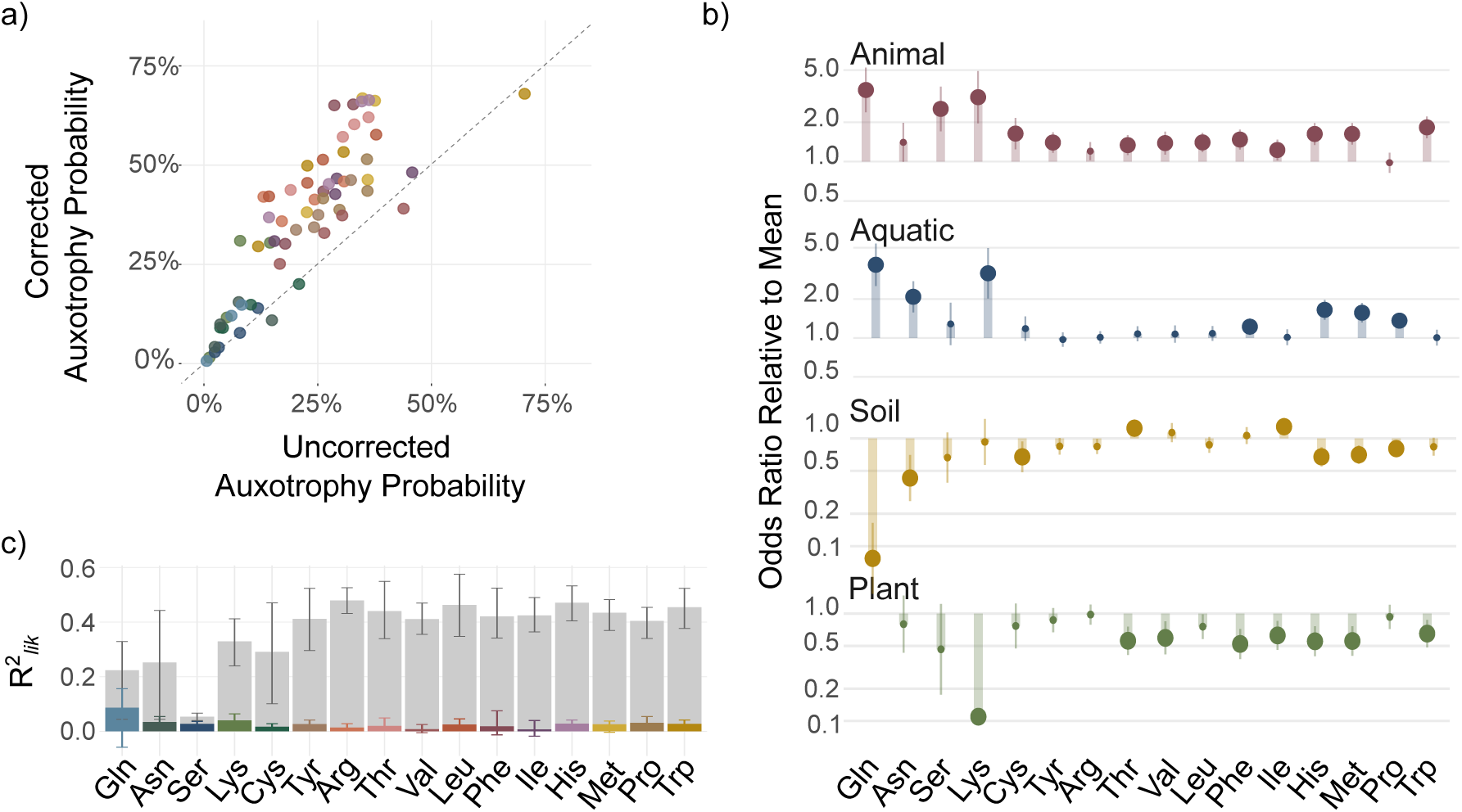
Effects of environment and phylogeny on the odds, probability, and predictability of individual auxotrophies. a) The effect of phylogenetic correction shown by comparing predicted absolute probability of auxotrophy between phylogenetically corrected (PhyloGLM) and uncorrected (GLM) models for each environment–amino acid pair. Points (colored by amino acid) above the diagonal reflect scenarios where phylogenetic correction reveals higher auxotrophy prevalence than the GLM, while points below the diagonal reflect cases where correction reduces the predicted probability of auxotrophy. Because no glutamine auxotrophy was predicted for plant-associated genomes, models for glutamine compared only animal-associated, soil, and aquatic environments. b) The effect of environment on the odds of auxotrophy relative to the grand mean across all four environments. Values exceeding one indicate an increased likelihood of auxotrophy compared to the grand mean, while values less than one indicate decreased likelihood. Large points represent a condition that differs significantly from the grand mean (BH-adjusted p < 0.05) Error bars represent the 95% confidence interval for the estimate. c) Variance partitioning comparing the contributions of phylogeny and environment to auxotrophy status. The relative importance of phylogenetic history versus environmental niche was quantified using the likelihood-based pseudo-R². The coloured portion of each bar indicates the contribution of environment and the grey portion the contribution of the phylogenetic tree. Error bars show +/-1 standard error derived from a leave-one-class-out jackknife analysis, designed to determine model sensitivity to clade composition (n = 13 classes; see SI Methods S14). Error bars are wider for rare auxotrophies, likely because auxotrophs for these amino acids are phylogenetically concentrated in a small number of lineages, and removing those lineages substantially destabilises the model estimate (Supplementary Figure 16).

We then examined the role of the environment in shaping auxotrophy patterns. We compared the corrected distribution of auxotrophy odds across environments normalized to the grand mean across all four environments (Figure 3b). In animal-associated and aquatic MAGs, auxotrophy rates are higher than the grand mean (with phylogenetic correction, significantly higher for 13 and 7 amino acids, respectively). Conversely, plant-associated and soil MAGs exhibit generally lower auxotrophy rates (with phylogenetic correction, significantly lower for 8 and 6 amino acids, respectively) (Figure 3b). Notably, plant-associated bacteria possess markedly less auxotrophy than animal-associated bacteria, suggesting that host identity may shape the selection of auxotrophy and that plant hosts may not readily release exogenous amino acids. Phylogenetic correction significantly altered 16 of 64 amino acid–environment coefficients (Clogg test, p ≤ 0.05), both unmasking previously obscured signals and attenuating apparent inflation due to phylogenetic clustering (Supplementary Figure 2; Supplementary Discussion 4). Overall, our results establish that phylogenetic correction is not optional but necessary for valid inference about environmental and genomic predictors of auxotrophy. Leave-one-class-out cross-validation confirms that the environment signal generalises across held-out phylogenetic lineages (Supplementary Figure 13).

We next investigated the relative importance of the environment and phylogenetic history in explaining auxotrophy distributions. To quantify the contributions of these features, we performed a variance partitioning analysis using likelihood-based pseudo-R² (Ives 2019). Phylogenetic structure is an important factor for predicting auxotrophy across all amino acids (mean R²_phylo_ = 0.36, range 0.03–0.47; Figure 3c) while the unique contribution of the environment is much smaller in absolute terms across amino acids (mean R²_env_ = 0.025; R²_env_ < 0.04 all amino acids except glutamine (R²_env_ = 0.087). We note that these two components are not directly comparable because phylogeny is represented as a continuous tree with full branch-length resolution while the environment is encoded as four coarse categorical labels. R² values can thus not be interpreted as a formal ranking. Nevertheless, the consistent importance of phylogenetic structure, combined with the observation that increasing environmental resolution from 4 to 12 classes yields nearly identical results (mean change Akaike Information Criterion (ΔAIC) 92.4 vs 95.9; Supplementary Figure 12), suggests that the limited explanatory power of the environment is not simply an artifact of label coarseness, and may rather reflect both the imprecision of categorical habitat labels for describing an environment’s chemical metabolite composition and a decoupling of present-day sampling location from ancestral selective pressures (Supplementary Discussion 6).

The existence of a deep phylogenetic signal has direct implications for the mechanistic frameworks proposed to explain gene loss. The BQH predicts that auxotrophy should be dynamic, tracking the availability of specific metabolites provided by community members, and auxotrophic states would thus be expected to vary within phylogenetic clades as organisms encounter different community contexts. However, the observed conservation of auxotrophic states across millions of years of divergence are inconsistent with this picture. Similarly, a simple cost-benefit model of independent gene loss would not predict auxotrophy tracking with phylogenetic history across deep evolutionary scales. Instead, the pattern we observe is most consistent with genome streamlining, in which auxotrophy reflects a transition to a metabolically dependent lifestyle rather than an environment-driven response. These frameworks need not be mutually exclusive: biosynthetic cost and leakage from coexisting microbes may influence which pathways are prioritised for loss, but genome streamlining is the process by which these losses become fixed and persist.

### 4. Genome functional composition predicts auxotrophy and changes with auxotrophy count

The limited effect of the environment on auxotrophy patterns suggests that other factors — namely, an organism’s lifestyle and functional capabilities — drive the evolution of auxotrophy. To characterize these signatures, we quantified genome functional composition as the fraction of protein-coding genes assigned to each of 18 Clusters of Orthologous Groups (COG) functional categories (see Methods), and used PhyloGLMs to examine how the prevalence of each COG category predicts amino acid auxotrophy (Figure 4a) (Cantalapiedra et al. 2021; Tatusov et al. 1997). We observed that COGs tend to consistently associate with auxotrophy across multiple amino acids. For example, across amino acids, auxotrophy is associated with an increase in relative gene content classified as translation machinery, while there is a consistent reduction in gene content dedicated to signal transduction. However, in some cases, COGs showed heterogeneous effects on auxotrophy odds. For example, across most amino acids, auxotrophy is significantly negatively associated with the fraction of genes involved in motility. In contrast, for methionine and isoleucine, this association is significantly positive, highlighting that gene content may influence auxotrophy in a pathway-specific manner.

**Figure 4:**
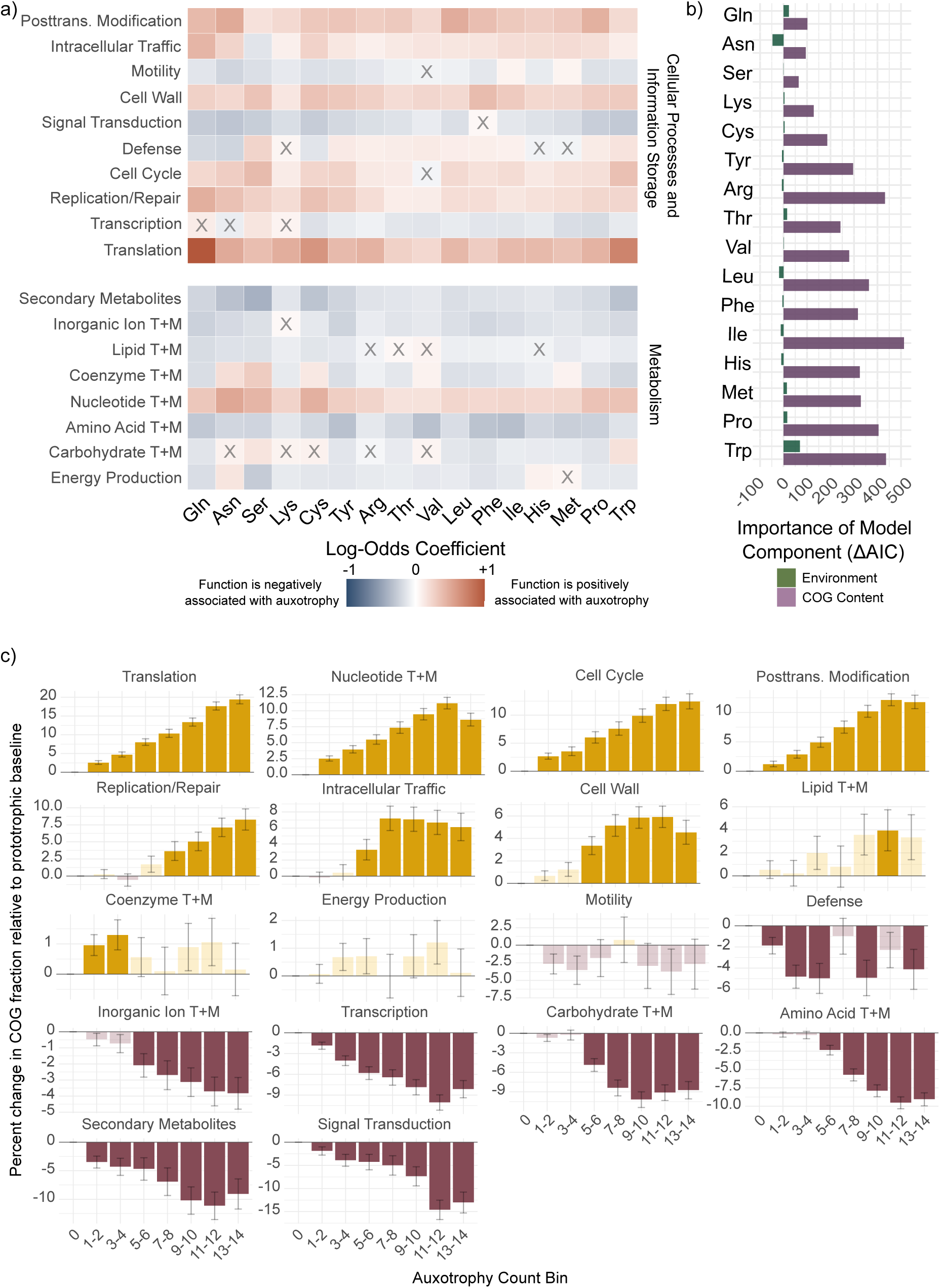
Auxotrophy status is related to the functional content of genomes. a) Associations between genome functional composition and amino acid auxotrophy. Each cell shows for each amino acid the estimated effect of the relative gene content dedicated to a Clusters of Orthologous Groups (COG) on odds of auxotrophy from phylogenetic logistic regression models (specifically, the log-odds coefficient represents the effect of a 1-standard deviation increase in the CLR-transformed fractional representation of a COG functional category on the log-odds of auxotrophy for a given amino acid). Amino acids are ordered by global auxotrophy prevalence (lowest to highest). Color indicates effect direction and magnitude; cells marked with a cross are non-significant after Benjamini–Hochberg correction (adjusted p < 0.05). “T+M” indicates transport and metabolism, and “Posttrans. Modification” is posttranslational modification. Models for translation reached the btol boundary for several amino acids; coefficients are conservative lower-bound estimates (see Supplementary Methods). b) Relative contributions of broad environmental class and genomic functional composition (pooled COGs) to explaining amino acid auxotrophy, quantified as change in Akaike Information Criterion (ΔAIC) from predictor-drop-out model comparisons. Bars represent the increase in AIC when each predictor set is removed from the full phylogenetic logistic regression model; larger values indicate a greater contribution to model fit, while negative values indicate that models including that predictor perform worse. Translation (COG J) and amino acid transport and metabolism (COG E) were excluded from this analysis; see Methods for details. c) Changes in the proportion of enzymes assigned to each COG functional category based on the number of amino acid auxotrophies per genome. Bars show the model-estimated percentage change in COG fraction relative to fully prototrophic genomes (bin 0), from phylogenetic linear models fitted per COG category. Faded bars are non-significant after Benjamini–Hochberg correction (adjusted p < 0.05).

To assess the relative importance of the environment versus functional capabilities on auxotrophy patterns, we performed a leave-one-out model comparison using the Akaike Information Criterion (AIC) (Figure 4b). Removing COG predictors substantially worsened model fit across all amino acids, indicating that they are essential for explaining auxotrophy patterns. In contrast, removing environmental class did not substantially degrade model performance. This suggests that the explanatory power initially attributed to the environment is in fact largely redundant with the genomic functional profile, and gene functional categories are more direct predictors of auxotrophy status (Supplementary Figure 8).

The consistency of COG–auxotrophy associations across amino acids indicates that these relationships are not amino acid specific but reflect a broader genomic pattern. Indeed, when considering the total number of auxotrophies per genome, these associations often become progressively stronger as auxotrophy count increases (Figure 4c). This indicates further that auxotrophy represents a cumulative genomic state rather than a set of independent traits. The strongest signal is a monotonic increase in the *fraction* of genes assigned to translation machinery as auxotrophy count rises. In genomes carrying 13–14 auxotrophies, 10.8% of protein-coding genes are assigned to translation (COG J), compared to 9.1% in fully prototrophic genomes — a difference of 1.73 percentage points, representing a 19% relative enrichment over the prototrophic baseline (p < 10⁻⁵⁶). Importantly, this does not represent an increase in the absolute number of translation genes, but rather a consequence of genome reduction: as biosynthetic, regulatory, and metabolic genes are progressively lost, the translation core is retained, and therefore constitutes a larger share of an overall smaller gene complement. Several other functional categories increased in parallel, while amino acid biosynthesis, carbohydrate metabolism, transcription, signal transduction, and secondary metabolism all declined sharply — consistent with differential gene loss rather than targeted enrichment of any single function (Figure 4c). Leave-one-clade-out cross-validation confirms that these patterns generalise across phylogenetic lineages (Supplementary Figures 14, 15 and 19; Supplementary Methods S13). Together, this is the genomic signature expected under streamlining: a shrinking genome in which the translational core is retained.

The coordinated nature of these genomic shifts, involving multiple functional categories simultaneously and scaling monotonically with number of auxotrophies, suggests that auxotrophy is not a collection of independent adaptive responses to individual metabolite availability, as cost-benefit or Black Queen frameworks would predict in isolation, but rather reflects a coherent and progressive reorganisation of genome content consistent with genome streamlining.

### 5. Auxotrophies accumulate non-independently and depend on other auxotrophies

To further investigate this coordinated shift toward a streamlined lifestyle, we tested whether multiple auxotrophies arise independently or whether their co-occurrence is fundamentally linked to this streamlined state. We first estimated how often double and triple auxotrophic mutants should appear if each auxotrophy were independent of the others, and compared this to the observed co-occurrence frequencies. We found that amino acid auxotrophies co-occurred far more frequently than expected under independence across all environment groups and phyla, for both pairs and triples (all p < 10⁻⁶; Figures 5a and b, Supplementary Figures 9 and 10).

**Figure 5:**
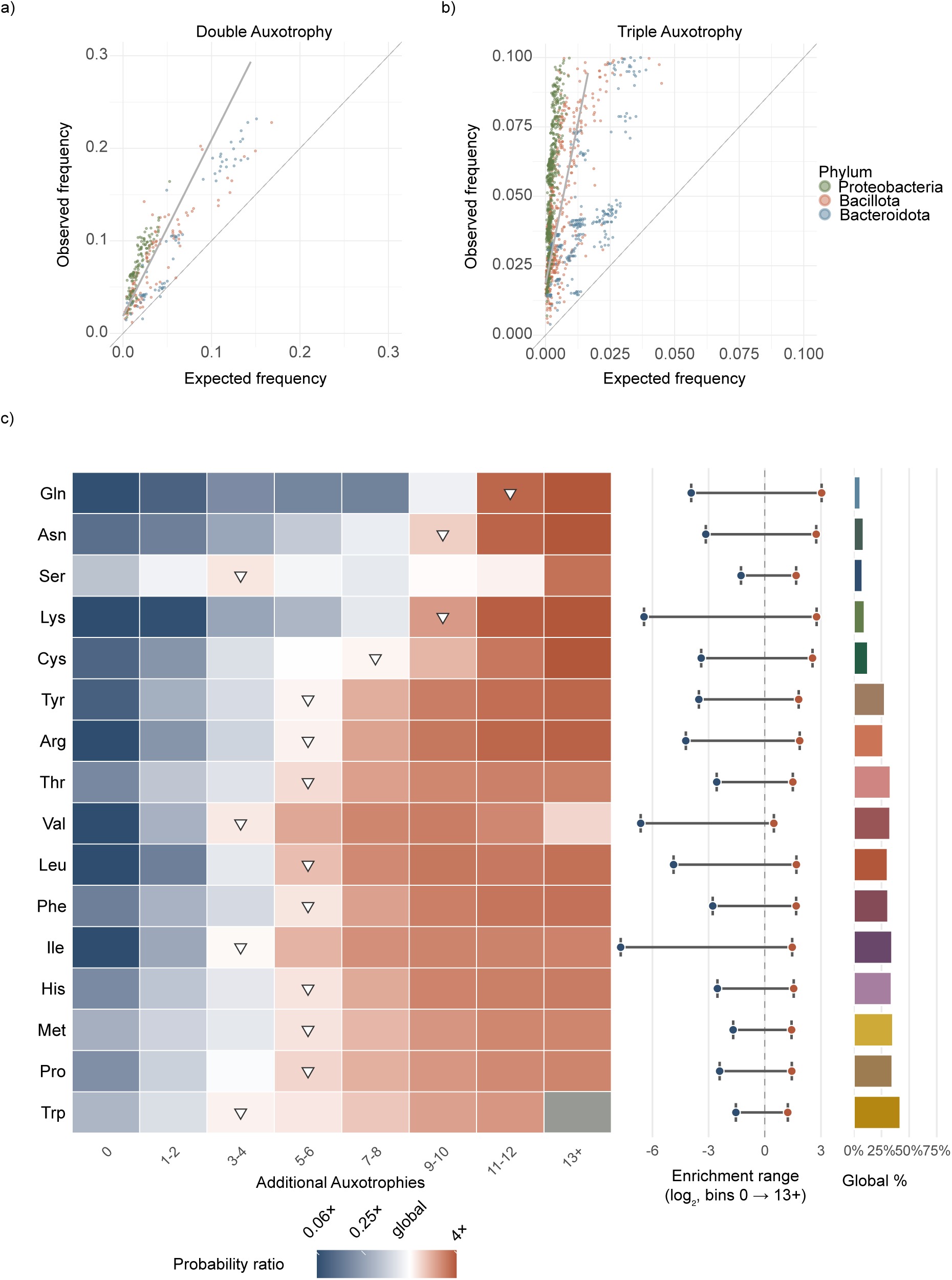
Auxotrophies accumulate non-independently. a) Observed versus expected double co-auxotrophy frequency across amino acid pairs. For each pair, the expected co-auxotrophy frequency is calculated as the product of individual amino acid frequencies (independence assumption); the dashed diagonal indicates perfect independence. Points are coloured by phylum (Proteobacteria, Bacillota, Bacteroidota); the solid grey line shows the linear regression across all phyla combined. A slope greater than 1 indicates that double co-auxotrophies are more common than expected by chance b) As in a), but for triple auxotrophy. c) Heatmap showing the predicted loss rate in each bin relative to the genome-wide loss rate for that amino acid (probability ratio; white = 1×, global average). Triangles mark the bin when the auxotrophy first becomes overrepresented (where the auxotrophy probability surpasses the global probability). Segments show the enrichment range from bin 0 (genome has no other amino acid losses) to the highest available bin; a longer segment indicates that the predicted loss probability changes substantially across the spectrum of genome reduction, while a shorter segment indicates that the probability remains relatively constant regardless of how many other auxotrophies are present. The bar chart shows global auxotrophy prevalence for each amino acid.

To characterize how individual amino acid losses depend on the overall number of auxotrophies, we modelled the predicted probability of each amino acid auxotrophy as a function of the number of other auxotrophies present in the genome (grouped into bins), then expressed this as a log₂ enrichment ratio relative to each amino acid’s auxotrophy frequency across the dataset (Figure 5c). The reference category (bin 0) represents genomes in which the focal amino acid is the only auxotrophy, providing an estimate of the probability of losing a given pathway in an otherwise prototrophic background. This baseline probability is low across all amino acids (median: 2.4%), indicating that most losses do not arise in isolation.

Across amino acids, auxotrophies are generally underrepresented when there are none or few other auxotrophies and overrepresented when there are many other auxotrophies, further confirming that auxotrophies typically co-occur. For each amino acid auxotrophy, the transition from under- to overrepresented as a function of other auxotrophies already present is marked by the crossover point at which it becomes overrepresented (diamonds in Figure 5c, left panel). This crossover point broadly mirrors global prevalence of the auxotrophy (Figure 5c, right panel): The most frequent auxotrophies tend to cross at low auxotrophy counts, while the rarest are restricted to nearly eroded genomes, suggesting that crossover position and global prevalence both reflect the same underlying property, potentially the intrinsic dispensability of each biosynthetic pathway, and that losses follow a consistent priority ordering rather than accumulating stochastically. This hierarchy likely reflects a combination of biosynthetic cost, pathway length, and metabolic network centrality rather than any single factor, consistent with streamlining acting on the overall cost of maintaining a pathway rather than any single property.

In addition to the crossover point at which amino acid auxotrophies become overrepresented, we can also characterise how strongly the loss probability changes across the range of auxotrophy burden. We quantify this as the enrichment range: the difference in predicted loss probability between genomes with no other auxotrophies (bin 0) and genomes with the maximum observed number of other auxotrophies (see Methods). A large enrichment range indicates that the amino acid is rarely lost in isolation but becomes increasingly likely to be lost as genomes become more eroded; a small range indicates that the probability of loss is relatively constant across all genomic backgrounds.

For instance, the branched-chain amino acids valine and isoleucine have near-zero probabilities of occurring as sole auxotrophies (0.3% and 0.2%) but once other auxotrophies are present they soon become overrepresented, with a large enrichment range consistent with a shared biosynthetic pathway that mechanistically couples their co-loss (Kaleta et al. 2013). In contrast, tryptophan (42% global auxotrophy) has a relatively high probability of occurring as a sole auxotrophy (14.5% at bin 0) and a small enrichment range, indicating that its loss is much less dependent on genomic context — consistent with the high biosynthetic cost of tryptophan synthesis creating selection for its loss across a wide range of genomic backgrounds. Serine also crosses early (bin 3–4) despite low prevalence, but its small enrichment range likely reflects metabolic model misclassification rather than a genuine biological signal.

Most amino acids become overrepresented at intermediate auxotrophy counts, representing a core set of amino acids whose losses are preferentially associated with genomes of moderate-to-high overall auxotrophy. Crucially, the rarest auxotrophies, such as lysine and glutamine (excluding serine), become overrepresented only at the highest observed auxotrophy counts. For instance, glutamine (the rarest globally at 5.9%) becomes overrepresented only when there are more than 10 other auxotrophies, while lysine (9.7%) when there are more than 8 other auxotrophies. Notably, lysine is almost never lost as a sole auxotrophy (0.1% at bin 0) and has one of the largest enrichment ranges in the dataset, suggesting that lysine biosynthesis loss is strongly dependent on the presence of other amino acid auxotrophies.

This consistent priority ordering of losses, with rare auxotrophies restricted to genomes already carrying many other losses, is consistent with sequential accumulation, whether driven by genomic cascades (where each loss reshapes the metabolic background, rendering further losses more likely) or ecological feedback (where initial auxotrophies drive niche shifts that progressively relax selection on additional pathways). Distinguishing between these mechanisms would require tracing the order of auxotrophy acquisition across phylogenetic lineages, which is beyond the scope of this study (see Supplementary Discussion 5). The enrichment profiles and priority ordering were highly stable when log genome size was added as a fixed-effect covariate (Pearson r = 0.955 across all amino acid and bin combinations; Supplementary Figure 18), ruling out undifferentiated gene loss in smaller genomes as an explanation for the observed structure.

Taken together, the universal excess of auxotrophy co-occurrence across phyla and environments, the consistent priority ordering of amino acid biosynthesis losses, and the increase of co-occurrence from pairs to triples is difficult to reconcile with the BQH as the primary driver of auxotrophy. The BQH predicts ecologically contingent, environment-specific co-occurrence patterns dependent on local community provisioning. Therefore, under the BQH, we would not expect the consistent excess of co-occurrence we observe across all environments and phyla. Furthermore, the ordering across all amino acids correlates only partially with published biosynthetic cost rankings (Akashi and Gojobori 2002; Kaleta et al. 2013), suggesting that the cost-benefit framework is one contributor to the dispensability hierarchy but does not fully determine it. The broad, environment-independent accumulation we observe instead points toward a genome-level process consistent with streamlining, in which losses accumulate progressively along a shared trajectory of increasing metabolic dependency.

## Conclusions

Our findings provide a coherent picture of amino acid auxotrophy as a phylogenetically structured genomic state rather than an environment-driven, amino acid-specific response to nutrient availability. By applying phylogenetically corrected models to an extensive MAG-derived dataset, we demonstrate that evolutionary history and genomic functional composition are the primary predictors of auxotrophy, often masking or absorbing signals previously attributed to the environment. The coordinated nature of these shifts — characterized by a monotonic enrichment of translational machinery alongside the systematic depletion of regulatory and biosynthetic breadth — suggests that auxotrophy is not a collection of independent adaptive events. Instead, the universal excess of co-occurring losses and the consistent hierarchical ordering of pathway erosion align most closely with genome streamlining theory. In this framework, metabolic dependency is a progressive, directional trajectory where the probability of losing a pathway is governed by the existing genome content. The hierarchical ordering we describe is consistent with any process driving progressive genome reduction, whether adaptive streamlining in large free-living populations or drift-driven erosion in obligate host-associated lineages, and may reflect a general trajectory of genome minimisation.

The global analysis performed here is necessarily insensitive to processes operating at finer phylogenetic or ecological scales. At the level of individual clades or specific environments, biosynthetic cost and Black Queen drivers are not excluded, and they may be the mechanism by which individual losses are initiated: a lineage colonising an amino-acid-rich niche may lose the most expensive pathway first, or a lineage embedded in a community may first lose a particular pathway because a dominant community member reliably leaks that metabolite. However, these frameworks alone cannot account for the global structure we observe, in which the auxotrophic state is predictable from phylogenetic relationships and genomic signatures across the bacterial tree as a whole. What our global models detect is the aggregate outcome of these local processes: a phylogenetically conserved, progressively accumulating pattern of gene loss consistent with genome streamlining as the overarching trajectory.

Overall, this study provides a new framework for interpreting metabolic dependencies and the evolutionary drivers of genome minimization in natural microbial systems.

## Methods

### Genome selection and quality control

Bacterial genomes were retrieved from the proGenomes3 database, which includes genomic sequences that are assigned to species-specific identifier clusters (specI) according to similarity of 40 marker genes. We selected 18,608 representative genomes from each of these clusters that included environmental information (Fullam et al. 2023). Genome quality was assessed using CheckM2 (v1.0.1).

To account for potential overprediction of auxotrophies due to genome fragmentation, we performed a correlation analysis between completeness and auxotrophy (predicted via the metabolic modeling pipeline described below). A statistically significant negative relationship was observed (Spearman’s ρ = −0.35, p < 2.2E−16; Supplementary Fig. S1). To mitigate this bias, genomes were filtered using a strict quality-control threshold of ≥85% completeness and ≤2% contamination, in accordance with the standards established by Starke et al. (2023). This high-quality subset of 17,392 genomes was used for all subsequent metabolic analyses.

### Genome-scale metabolic modelling

Genome-scale metabolic models (GEMs) were reconstructed using the gapseq pipeline (v1.2, commit e3210c0f) (Zimmermann et al. 2021). The ‘gapseq doall’ command was utilized to automate pathway prediction, reaction network assembly, and iterative gap-filling based on the predicted protein sequences. For taxa identified as obligate anaerobes in the proGenomes3 metadata, oxygen was removed from the simulated medium by setting the oxygen concentration to zero (cpd00007:0) during the medium specification and gap-filling stages.

Following the methodology validated by Starke and coworkers (Starke et al. 2023), we utilized genome-specific gap-filling media rather than a standardized medium for all models. This approach ensures that the gap-filling process is informed by the unique metabolic potential of each genome, reducing the risk of erroneous predictions (see Starke et al. 2023 for details). After reconstruction, we maintained only those models capable of achieving a predicted baseline growth rate of at least 0.01 h^−1^ when growth is simulated in a nutrient complete medium (see details on metabolic modelling below) resulting in a final high-quality dataset of 16,398 functional metabolic models.

### Assigning genome metadata

We retrieved isolation source metadata for each genome using the Genome Taxonomy Database (GTDB) metadata (Release 10, R10-RS226, (Parks et al. 2022). Environmental information was extracted from the GTDB’s aggregation of NCBI BioSample records (ncbi_isolate_source). Due to the lack of standardization in free-text metadata, we implemented a custom rule-based classification system. Environmental sources were mapped to 16 granular categories (e.g., animal_digestive, engineered_system) using an extensive library of habitat-specific keywords (Supplementary Information). These were further aggregated into four broad categories: Aquatic, Animal-associated, Soil, and Plant-associated. Records that did not match any terms in the habitat description library were assigned to an “Other” category.

Additionally, we distinguished metagenome-assembled genomes (MAGs) from isolates using the ncbi_genome_category field. We classified a genome as a MAG if the field contained “derived from metagenome” and as “Isolate/Other” in all other cases. Of the 16,398 functional models, this yielded 8,249 MAGs and 8,149 isolate genomes.

### Predicting auxotrophies

Metabolic simulations were executed using flux balance analysis (FBA) with an objective function set to the flux through the biomass formation reaction. Simulations were conducted in R (v4.3.1) using the sybil (v2.2.1) package, with the IBM ILOG CPLEX optimizer via the cplexAPI library (v22.1.2) (Gelius-Dietrich et al. 2013; “IBM ILOG CPLEX Optimization Studio” 2024). To identify amino acid dependencies, we established a baseline biomass flux for each model in a simulated rich medium containing all amino acids. We then performed systematic amino acid dropout simulations by iteratively setting the lower bound of specific exchange reactions (EX_[compound]_e0) to zero. A genome was classified as an auxotroph if the biomass flux dropped below an absolute threshold of 0.01 h^−1^, representing a near-total cessation of growth (≤1% of typical baseline flux).

### Benchmarking against experimental data

To evaluate the reliability of our metabolic reconstructions, we benchmarked the gapseq pipeline described above against a curated dataset of 223 bacterial genomes with published phenotypic auxotrophy data (Ramoneda et al. 2023; Starke et al. 2023; Price et al. 2020). Some of these genomes were experimentally screened for auxotrophy for each amino acid and others were screened for a subset of amino acids, yielding 3,939 total observations of auxotrophy or prototrophy (3,797 prototrophies and 142 auxotrophies). Notably, this dataset is very unbalanced, with 3,797 of these observations corresponding to prototrophy and only 142 to auxotrophy. To evaluate the performance of GEM-based auxotrophy predictions, we calculated several key metrics that describe the model’s reliability: models achieved an overall accuracy of 97.36%, with a specificity of 98.27% and a recall (sensitivity) of 78.02%.

### Phylogenetic tree

The GTDB bac120 phylogenetic tree (R226) was pruned to the taxa in our filtered dataset, rooted on Fusobacteria, and rescaled to unit height (total tree height = 1). Rescaling ensures that the phylogenetic rate parameter α and derived half-life values are on a comparable scale across datasets (Supplementary Methods S2).

### Phylogenetic signal: evolutionary half-life

For each of the 16 amino acids where auxotrophies were predicted, an intercept-only phylogenetic logistic regression was fitted using phyloglm (logistic_MPLE; phylolm v2.6.5) across 4,507 MAGs from aquatic, animal-, soil-, and plant-associated environments. The evolutionary rate parameter α was converted to a phylogenetic half-life t₁/₂ = ln(2)/α, representing the branch length at which phylogenetic correlation decays to 50%. The half-life was validated against a null distribution from 1,000 random permutations of trait assignments. Full model specification is in Supplementary Methods S2.

### Prediction of effect of environment

For each amino acid, binary auxotrophy status was modelled as a function of broad environment class using phylogenetic logistic regression (phyloglm, logistic_MPLE). Environment was encoded with sum-to-zero contrasts so each coefficient represents a deviation from the grand-mean log-odds of auxotrophy. The L-Glutamine model used only three environment classes (animal, aquatic, soil) because no plant-associated genomes in the dataset were glutamine auxotrophs. Null (intercept-only) models were fitted for each amino acid for ΔAIC and likelihood ratio test (LRT) comparisons. Explanatory power was quantified as partial likelihood-based R² (R²_lik_) using the rr2 package. All p-values were adjusted using the Benjamini–Hochberg (BH) procedure. Full model specification and post-hoc contrasts are described in Supplementary Methods S3.

### Prediction of effect of genome content (COG composition)

Functional gene content was characterised using COG category annotations from eggNOG-mapper. Per-genome COG fractions were calculated as the percentage of annotated genes (excluding unknown/unannotated genes) in each category; genes with multiple COG assignments were split before counting. COG categories absent from ≥1% of genes in ≥100 genomes, and five eukaryote-specific or rare categories (Y, A, B, W, Z), were removed prior to analysis (18 categories retained). COG E (amino acid transport and metabolism) and COG J (translation) were excluded from multivariate analyses to avoid circularity and because of boundary convergence issues, respectively (see Supplementary Methods S4).

Marginal associations were estimated by fitting a univariate phylogenetic logistic regression (phyloglm, logistic_MPLE) for each amino acid × COG combination (16 × 18 = 288 models; BH-adjusted p-values). To decompose unique contributions of genomic composition and environment, full phyloglm models (environment + all COG fractions) were compared against models omitting each predictor block, and against standard GLMs lacking phylogenetic correction, using ΔAIC, partial R², and LRTs (Supplementary Methods S5).

### COG composition shifts with auxotrophy count

To test whether genomic functional composition shifts systematically with total auxotrophy count, we binned genomes by total auxotrophy count into paired groups (0, 1–2, 3–4, …, 13–14), with fully prototrophic genomes (bin 0) as reference. For each COG category, a phylogenetic linear model (phylolm, Pagél’s λ) was fitted with auxotrophy bin as the categorical predictor. Coefficients represent percentage-point shifts in COG fraction relative to the prototrophic baseline. For visualisation, shifts are expressed as percentage change from baseline: δ/μ₀ × 100%, where δ is the bin coefficient and μ₀ is the estimated prototrophic baseline fraction (Supplementary Methods S6).

### Co-auxotrophy: observed versus expected

For all pairwise (*n* = 120) and three-way (*n* = 560) combinations of the 16 amino acids, expected co-auxotrophy frequency was calculated as the product of individual amino acid frequencies under statistical independence (e.g., freq(A) × freq(B) for A–B double auxotrophs). We then compared these expected frequencies to the actual observed co-occurrence frequencies in our dataset using linear regression. Observed co-auxotrophy frequency was regressed on expected frequency using ordinary least squares, and tested against a null model of perfect independence (slope = 1, intercept = 0) using one-sample t-tests. Analyses were performed globally and stratified by broad environment class and phylum (Supplementary Methods S7).

### Auxotrophy identity predicted by auxotrophy count

To test whether total auxotrophy count predicts the loss of specific amino acids, we fitted a separate phylogenetic GLMM (PGLMM; phyr::pglmm, binomial family) for each amino acid. Because total auxotrophy count includes the focal amino acid’s own loss, using it directly as a predictor would be partially circular. We therefore subtracted the focal amino acid’s contribution from each genome’s total count before fitting: for a genome auxotrophic for the focal amino acid, the predictor was total count minus one; for a prototrophic genome, the predictor was the total count unchanged. This corrected count was grouped into eight bins (0, 1–2, 3–4, 5–6, 7–8, 9–10, 11–12, 13+), with bin “0” as the reference level, representing genomes with no other amino acid losses. Bins in which all genomes shared the same auxotrophy status for the focal amino acid were excluded before fitting, as no coefficient can be estimated from a response with no variation. Model significance was assessed by likelihood ratio test against an intercept-only PGLMM fitted on the same observations (χ², df = up to 7, fewer for amino acids where one or more bins were excluded). Model-predicted probabilities were converted to log₂ enrichment ratios relative to each amino acid’s global loss prevalence. Full model specification, enrichment-range and crossover-point definitions are provided in Supplementary Methods S8.

### Software

All analyses were performed in R (v4.4.1)(R Core Team 2024) using the following key packages: phylolm (v2.6.5)(Tung Ho and Ané 2014) for phylogenetic logistic and linear regression, phyr (v1.1.3)(Li et al. 2020) for PGLMMs, rr2 (v1.1.1)(Ives and Li 2018) for likelihood-based R², ape (v5.8-1)(Paradis and Schliep 2019) for tree manipulation, and tidyverse (v2.0.0)(Wickham et al. 2019) for data wrangling. Figures were produced with ggplot2 (v3.5.0)(Wickham 2016) and exported as SVG via svglite (v2.2.2)(Wickham H et al. 2025). Claude (Anthropic) and Gemini (Google DeepMind) were utilized to debug analysis code, and to assemble supplementary materials. All outputs were reviewed, verified, and edited by the authors, who take full responsibility for the accuracy and integrity of the published work.

## Data Availability

The analysis code is currently hosted in a private repository and is available from the corresponding author upon reasonable request. The repository will be made publicly available upon peer-reviewed publication.

## Acknowledgements

This work was supported by the Simons Collaboration on Principles of Microbial Ecosystems (PriME, 542379FY22 and LS-PRIME-00011858 to M.A.). R.E.S. was supported by an EMBO Postdoctoral Fellowship (ALTF 867-2024) and an ETH Zurich Postdoctoral Fellowship (24-2 FEL-068). This research was also supported as a part of the NCCR Microbiomes, funded by the Swiss National Science Foundation (grant nos. 51NF40_180575 and 51NF40_225148). We thank Astrid Stubbusch for helpful discussions and Joshua Bloom for feedback on the manuscript. Computational resources were provided by the Euler cluster at ETH Zurich.

