## Supplementary Materials for "Amino acid auxotrophy is a feature of a distinct bacterial lifestyle across environments"

Alyssa Henderson *et al.*

**Contents**

1. Supplementary Methods (pages 2–20)
2. Supplementary Figures S1–S21 (pages 21–41)
3. Supplementary Tables S1–S8 (pages 42–74)
4. Supplementary Discussion 1–7 (pages 75–79)

### Supplementary Methods

---

#### S1. Genome Quality Filtering and Environment Classification

Auxotrophy predictions from gapseq were merged with GTDB bac120 genome metadata. Genomes were retained if they satisfied both quality thresholds:

$$\text{Completeness} \geq 85\% \quad \text{and} \quad \text{Contamination} \leq 2\%$$

All downstream analyses were further restricted to metagenome-assembled genomes (MAGs; `metagenome == "YES"`) to avoid the systematic biases in isolate collections (laboratory cultivation selects against obligate auxotrophs). A separate MAG vs. isolate comparison is shown in Figure S1.

**Environment classification.** NCBI isolation source free-text strings were classified into fine-grained environment categories using prioritised regular-expression rules evaluated by `case_when` (i.e., earlier rules take precedence). Categories were then rolled up into four broad classes used in all phylogenetic models:

| Broad class | Fine-grained categories included |
|---|---|
| **animal** | animal_digestive, animal_other |
| **aquatic** | marine, freshwater, aquatic_other |
| **soil** | soil |
| **plant** | plant_associated |

Genomes not matching any of the four broad classes (e.g., sediment, engineered systems, extreme environments) were assigned to an "other" category and excluded from environment-modelling analyses.

##### S1.2. Genome-scale metabolic model reconstruction

Genome-scale metabolic models (GEMs) were reconstructed using gapseq (v1.2, commit e3210c0f; Zimmermann et al. 2021) with the `doall` command, which automates: (i) pathway prediction from predicted protein sequences, (ii) reaction network assembly, and (iii) iterative gap-filling against a genome-specific medium. Using genome-specific gap-filling media rather than a universal medium reduces erroneous pathway additions caused by pathways that are broadly present in bacteria but absent from the focal genome (see Starke et al. 2023 for validation).

For taxa classified as obligate anaerobes in the proGenomes3 metadata, oxygen was excluded from the gap-filling medium by setting the oxygen exchange lower bound to zero (`cpd00007:0`). Models were retained only if they achieved a baseline biomass flux of $\geq$0.01 h$^{-1}$ in a nutrient-complete environment; this threshold excludes reconstructions in which the gap-filling failed to produce a coherent metabolic network. Of the 17,392 high-quality genomes, 16,398 passed this growth check and were used in all subsequent analyses.

##### S1.3. Auxotrophy prediction via flux balance analysis

Amino acid auxotrophies were predicted using flux balance analysis (FBA) implemented in R (v4.3.1) with the `sybil` package (v2.2.1) and the IBM ILOG CPLEX optimiser (v22.1.2; cplexAPI library). For each model, a baseline biomass flux was established in a simulated rich medium containing all amino acids. Auxotrophies were identified by systematic single-amino-acid dropout: the exchange reaction lower bound for each amino acid was set to zero in turn, and the biomass flux was re-optimised. A genome was classified as auxotrophic for amino acid $a$ if removing $a$ from the medium reduced predicted biomass flux below 0.01 h$^{-1}$ ($\leq$1% of the nutrient-replete baseline), representing a near-complete cessation of growth.

**Input data coding.** The resulting per-genome auxotrophy matrix uses the convention: 0 = auxotrophic, 1 = prototrophic. Where model coefficients are reported as log-odds of *auxotrophy*, the raw model output (which is on the prototrophy scale by convention) is negated; this negation is noted throughout the methods where applicable.

##### S1.4. Benchmarking against experimental data

To evaluate the accuracy of the gapseq-based predictions, we benchmarked against a curated dataset of 223 bacterial genomes with published phenotypic auxotrophy data (Ramoneda et al. 2023; Starke et al. 2025), comprising 3,939 total observations (3,797 prototrophies and 142 auxotrophies). Model performance metrics:

| Metric | Value |
|---|---|
| Accuracy | 97.4% |
| Specificity (true negative rate) | 98.3% |
| Recall / Sensitivity (true positive rate) | 78.0% |

False negatives (auxotrophies predicted as prototrophic) are the dominant error mode. They likely arise because GEMs rely on gene presence/absence: some phenotypic auxotrophies are caused by regulatory variation, non-synonymous SNPs, or pathways requiring only minimal gap-filling, all of which leave near-complete biosynthetic gene sets intact. Because non-functional biosynthetic pathways are typically subject to gene decay and eventual pseudogenisation over evolutionary timescales, gene-content-based predictions are expected to be conservative estimates of auxotrophy — that is, the true population-level auxotrophy frequencies are at least as high as those reported here.

---

#### S2. Phylogenetic Signal Strength: Evolutionary Half-Life

To quantify how rapidly auxotrophy states evolve independently of phylogenetic relatedness, we fitted an intercept-only phylogenetic logistic regression for each amino acid separately using `phyloglm` (logistic_MPLE; Ho & Ané 2014):

$$\text{logit}(P(\text{auxotrophic}_i)) = \mu_a + \varepsilon_i$$

**Data coding note.** In the input data, binary trait columns are coded 1 = prototrophic, 0 = auxotrophic. Internally, `phyloglm` therefore fits log-odds of prototrophy; all reported coefficients and intercepts are negated relative to the raw model output so that the equations and plots throughout this document are on the **auxotrophy** scale (positive values = higher probability of auxotrophy). This negation does not affect the $\alpha$ parameter, goodness-of-fit metrics (AIC, $R^2$_lik), or LRT statistics, which are invariant to the labelling of the two binary states.

where $\varepsilon_i$ follows a phylogenetic error structure governed by the evolutionary rate parameter $\alpha$. Under the logistic MPLE model, the covariance between taxa $i$ and $j$ decays exponentially with phylogenetic distance $d_{ij}$:

$$\text{Cov}(\varepsilon_i, \varepsilon_j) \propto e^{-\alpha\, d_{ij}}$$

The **phylogenetic half-life** is the phylogenetic distance at which this covariance decays to half its initial value:

$$t_{1/2} = \frac{\ln 2}{\hat{\alpha}}$$

A large $t_{1/2}$ (small $\hat{\alpha}$) indicates that phylogenetically distant relatives tend to share the same auxotrophy state (evolutionary stasis). A small $t_{1/2}$ (large $\hat{\alpha}$) indicates rapid, lineage-independent evolution. Branch lengths were rescaled to unit tree height before fitting so that $t_{1/2}$ values are comparable across datasets and to the $[0, 1]$ scale of the tree.

---

#### S3. Environment Predicts Auxotrophy: Phylogenetic Logistic Regression

##### S3.1 Model specification

For each of the 16 amino acids, we modelled binary auxotrophy as a function of broad environment class using `phyloglm` (method `logistic_MPLE`; see §S2 for data coding note):

$$\operatorname{logit}\big(P(\text{auxotrophic}_i)\big) = \mu_a + \sum_{k=1}^{K-1} \beta_k \cdot x_{ik} + \varepsilon_i$$

where $x_{ik}$ are sum-to-zero (effects) coded indicator variables for broad environment class (animal, aquatic, soil, plant; $K = 4$), and $\varepsilon_i$ represents phylogenetic error as in §S2. Under sum-to-zero coding, each $\beta_k$ represents the deviation of environment $k$ from the grand mean log-odds of auxotrophy, and the final level's coefficient is not estimated directly but is recovered algebraically:

$$\hat{\beta}_K = -\sum_{k=1}^{K-1} \hat{\beta}_k$$

The variance of $\hat{\beta}_K$ is derived from the full variance-covariance matrix $\boldsymbol{\Sigma}$ of the estimated coefficients:

$$\operatorname{Var}(\hat{\beta}_K) = \operatorname{Var}\left(-\sum_{k=1}^{K-1} \hat{\beta}_k\right) = \sum_{j=1}^{K-1}\sum_{l=1}^{K-1} \operatorname{Cov}(\hat{\beta}_j, \hat{\beta}_l) = \mathbf{1}^\top \boldsymbol{\Sigma}_{\text{env}} \mathbf{1}$$

where $\boldsymbol{\Sigma}_{\text{env}}$ is the $(K-1) \times (K-1)$ covariance submatrix for the environment coefficients.

**L-Glutamine special case.** No plant-associated genomes in the dataset were glutamine auxotrophs, causing complete separation and non-convergence of the four-environment model. L-Glutamine was therefore modelled separately using only three environment classes (animal, aquatic, soil), with sum-to-zero coding over $K = 3$ levels.

A null (intercept-only) model was also fitted for each amino acid for the purposes of ΔAIC and LRT comparisons (§S3.2, §S9).

Parallel standard GLMs (no phylogenetic correction; `glm(family = binomial)`) were fitted for supplementary comparison.

##### S3.2 Model comparison: ΔAIC

Model fit was compared using the Akaike Information Criterion:

$$\text{AIC} = -2\,\hat{\ell} + 2K_p$$

where $\hat{\ell}$ is the maximised log-likelihood and $K_p$ is the number of estimated parameters. Improvement in fit from adding the environment predictor was quantified as:

$$\Delta\text{AIC}_{\text{env}} = \text{AIC(null)} - \text{AIC(full)}$$

A positive $\Delta\text{AIC}_{\text{env}}$ indicates that the environment model fits better than the null; values $> 2$ are conventionally considered meaningful.

##### S3.3 Multiple testing correction

p-values from the z-tests on individual environment coefficients (two-sided, $z_k = \hat{\beta}_k/\hat{\sigma}_k$) were adjusted for multiple comparisons within each amino acid's coefficient table using the Benjamini–Hochberg (BH) false discovery rate procedure.

##### S3.4 Sensitivity analysis: fine-grained environment labels

To test whether the broad four-class environment scheme (S3.1) attenuates the measured environment signal through measurement imprecision (attenuation bias), we repeated the phyloglm analysis using fine-grained habitat labels (`class` column in the metadata). Fine classes were retained only if they contained $\geq 30$ MAGs to ensure stable coefficient estimation; 12 classes passed this threshold (animal_digestive, freshwater, engineered_system, marine, sediment, other, soil, extreme_environment, animal_other, coastal, plant_associated, other_host). Three classes were too small and excluded (built_environment, terrestrial_other, food_associated).

**Per-amino-acid filtering.** For each amino acid, fine classes in which all genomes were prototrophic (zero auxotrophs) were additionally excluded before model fitting to prevent complete separation and non-convergence. Classes that fell below the 30-genome threshold after this per-AA filtering were also removed. The predictor (`class`) was encoded with sum-to-zero (effects) coding over the retained classes for each amino acid independently.

The resulting full and intercept-only phyloglm models were fitted using the same specification as S3.1 (logistic_MPLE, btol = 50, unit-height-rescaled tree). Model fit was compared using:

$$\Delta\text{AIC}_{\text{fine}} = \text{AIC}(\text{intercept-only}_{\text{fine}}) - \text{AIC}(\text{full}_{\text{fine}})$$

This is compared directly to $\Delta\text{AIC}_{\text{broad}}$ from S3.2. Both are computed within their own dataset (fine models on the filtered subset, broad models on the full MAG set), so each $\Delta$AIC measures the improvement from adding environment over intercept-only on the same observations — a valid like-for-like comparison of label informativeness despite the differing sample sizes.

Under the attenuation hypothesis, finer labels should **consistently** increase ΔAIC across amino acids. Results are shown in Supplementary Figure L (script `10_sensitivity_fine_env.Rmd`).

---

#### S4. COG Category Composition

##### S4.1 Calculation of COG fractions

Per-genome COG functional category assignments were obtained from eggNOG-mapper applied to predicted protein sequences. Some genes carry multiple COG assignments (e.g., "EK"); these were split into individual categories before counting, so that each gene contributes proportionally to all applicable categories. The fraction of annotated genes in category $c$ for genome $g$ is:

$$f_{g,c} = \frac{n_{g,c}}{N_g^*} \times 100\%$$

where $n_{g,c}$ is the count of genes assigned to category $c$ (after splitting), and $N_g^*$ is the total number of annotated genes in genome $g$ after excluding function-unknown (COG S) and unannotated (–) genes.

##### S4.2 Excluded COG categories

Five COG categories were excluded globally as irrelevant or erroneous in prokaryotic genomes: **Y** (nuclear structure; eukaryote-specific), **A** (RNA processing; eukaryote-specific), **B** (chromatin; eukaryote-specific), **W** (extracellular structures; rare in bacteria), and **Z** (cytoskeleton; rare in bacteria).

In the multivariate importance analysis (§S5), two additional categories were excluded to avoid circularity: **E** (amino acid transport and metabolism) and **J** (translation, ribosomal structure). COG E is mechanistically coupled to auxotrophy by definition (auxotrophs have fewer amino acid biosynthesis genes); its inclusion would inflate the apparent predictive power of genomic composition. COG J similarly conflates cause and consequence of genome streamlining. Both categories are retained in the univariate analysis (§S5.1) for reference, where the btol-boundary warning flag indicates model unreliability.

> **Note:** COG E and COG J are included in the univariate models (script 05, `fit_univari` chunk) but excluded from the multivariate importance analysis (`get_AIC_rr2_per_predictor` chunk in script 05). The plot of univariate results therefore shows values for all 18 categories, while the ΔAIC decomposition (Figure 4b) uses 16 categories. This asymmetry is intentional: COG E is excluded to avoid circularity and COG J is excluded due to boundary convergence issues.

#### S5. COG Composition Predicts Auxotrophy: Multivariate Importance

##### S5.1 Univariate COG associations

**COG predictor preprocessing.** Because COG fractions are compositional data (they sum to approximately 100% per genome), the individual fractions are not statistically independent. To address this, COG fractions were transformed using the centred log-ratio (CLR) prior to z-scaling. CLR transforms each fraction relative to the geometric mean of all fractions for that genome:

$$\mathrm{CLR}(f_{i,c}) = \log\left(\frac{f_{i,c}}{g_i}\right), \quad g_i = \exp\left(\frac{1}{C}\sum_{c'} \log f_{i,c'}\right)$$

where $g_i$ is the row geometric mean across all $C$ COG fractions for genome $i$. Zeros were replaced with a pseudocount of 0.001% before log-transformation. CLR values were then z-scaled to mean 0 and SD 1 across genomes to make coefficients comparable in magnitude across categories. The resulting predictor $\tilde{f}_{i,c} = z(\mathrm{CLR}(f_{i,c}))$ is used in all phyloglm models. Note: after CLR, back-transformation to raw percentage points is not straightforward; coefficients are interpreted as the log-odds change per 1-SD increase in CLR-transformed COG fraction.

For each combination of amino acid (16) $\times$ COG category (18, including E and J), a separate phylogenetic logistic regression was fitted (see §S2 for data coding note):

$$\mathrm{logit}(P(\mathrm{auxotrophic}_i)) = \mu_a + \beta_c \cdot \tilde{f}_{i,c} + \varepsilon_i$$

where $\tilde{f}_{i,c}$ is the CLR-transformed and z-scaled COG fraction for genome $i$ in category $c$, and $\beta_c$ is the log-odds of **auxotrophy** per 1-SD increase in $\tilde{f}_{i,c}$. A positive $\beta_c$ means that a higher relative representation of genes in category $c$ is associated with greater probability of auxotrophy; a negative value indicates the opposite. Because the raw model estimate is on the prototrophy scale (§S2), the reported coefficient is `LogOdds = -estimate` — this is consistent across all figures and tables in the paper.

p-values were corrected across all COG $\times$ amino acid combinations using the BH procedure.

##### S5.2 Multivariate importance decomposition

For each amino acid, a full phylogenetic logistic regression was fitted with both broad environment class and all CLR-transformed, z-scaled COG fractions (excluding E and J; see §S5.1 for preprocessing) as predictors (see §S2 for data coding note):

$$\text{logit}(P(\text{auxotrophic}_i)) = \mu_a + \sum_k \beta_k x_{ik} + \sum_c \gamma_c \tilde{f}_{i,c} + \varepsilon_i$$

Two reduced models were compared to the full model to decompose predictor importance (within-class comparisons using phyloglm only, so that likelihood functions are matched):

| Comparison | Reduced model | ΔAIC formula | Interpretation |
|---|---|---|---|
| COG contribution | Full − COGs (env only) | AIC(no-COGs) − AIC(full) | Unique variance explained by genomic composition |
| Environment contribution | Full − environment (COGs only) | AIC(no-env) − AIC(full) | Unique variance explained by habitat |

A positive ΔAIC means the reduced model is worse — i.e., the dropped predictor(s) genuinely improve fit. Note: comparison of phyloglm to a standard GLM (to quantify the phylogenetic signal contribution) is not valid because the two model classes use different likelihood functions (MPLE vs. exact binomial MLE); this comparison is therefore not reported. The contribution of phylogeny is instead shown via $R^2_{\text{lik}}$ (Figure 3c).

**Partial $\mathbf{R}^2$ ($\mathbf{R}^2$_lik).** The partial coefficient of determination based on likelihood was computed using the `rr2` package:

$$R^2_{\text{lik}}(\text{full}, \text{reduced}) = 1 - \exp\left(\frac{2}{n}\left(\hat{\ell}_{\text{reduced}} - \hat{\ell}_{\text{full}}\right)\right)$$

where $\hat{\ell}$ denotes the maximised log-likelihood and $n$ is the sample size. Values near zero indicate the dropped predictor(s) contribute little beyond the retained predictors; values near 1 indicate they explain nearly all the residual variation.

**Likelihood ratio tests.** For the COG and environment contributions, a LRT was computed:

$$\Lambda = -2\left(\hat{\ell}_{\text{reduced}} - \hat{\ell}_{\text{full}}\right) \sim \chi^2_{(df)}$$

where *df* equals the number of additional fixed-effect parameters in the full model. For the phylogeny vs. GLM comparison, no LRT was computed because the two model classes (logistic_MPLE vs. standard binomial GLM) are not nested in the standard sense; ΔAIC serves as the comparison metric instead.

**Correctness note on LRT df for environment.** The code computes `df = length(coef(full)) - length(coef(no_broad))`. With sum-to-zero coding and

$K = 4$ environment levels, broadclass contributes $K-1 = 3$ coefficients. However, note that `phyloglm` with `logistic_MPLE` may report coefficients differently depending on whether the intercept is parameterised separately. Verify that the df for the broadclass LRT is consistently 3 across all amino acids.

---

#### S6. COG Composition Shifts with Total Auxotrophy Burden

##### S6.1 Auxotrophy count binning

Total per-genome auxotrophy count was computed as:

$$A_i = \sum_{a=1}^{16} \mathbf{1}[\mathrm{AA}_{i,a} = 0]$$

where $\mathrm{AA}_{i,a} = 0$ indicates that genome $i$ is auxotrophic for amino acid $a$ (using the input data coding where 0 = auxotrophic, 1 = prototrophic). $A_i$ therefore ranges from 0 (fully prototrophic) to 16 (auxotrophic for all modelled amino acids). Genomes were grouped into bins of width 2:

$$B_i \in \{0,\ 1\text{–}2,\ 3\text{–}4,\ 5\text{–}6,\ 7\text{–}8,\ 9\text{–}10,\ 11\text{–}12,\ 13\text{–}14\}$$

with bin "0" (fully prototrophic genomes; $A_i = 0$) as the reference level.

##### S6.2 Phylogenetic linear model

For each COG category $c$, a phylogenetic linear model (phylolm, Pagel's $\lambda$ covariance model) was fitted with auxotrophy bin as a categorical predictor:

$$f_{i,c} = \mu_c + \sum_{b \neq 0} \delta_{c,b} \cdot \mathbf{1}[B_i = b] + \varepsilon_i$$

where $\varepsilon_i \sim \mathcal{N}(0, \sigma^2 \boldsymbol{V}(\lambda))$, and $\boldsymbol{V}(\lambda)$ is the phylogenetic variance-covariance matrix under Pagel's $\lambda$ model:

$$V_{ij}(\lambda) = \lambda \cdot t_{ij} + (1 - \lambda) \cdot \mathbf{1}[i = j]$$

with $t_{ij}$ the shared branch length between taxa $i$ and $j$.

The intercept $\hat{\mu}_c$ is the estimated COG fraction for bin-0 (prototrophic) genomes; each $\hat{\delta}_{c,b}$ is the percentage-point shift relative to this baseline for genomes in bin $b$.

p-values were adjusted across all COG $\times$ bin combinations using the BH procedure.

**Why phylolm (lambda) rather than phyloglm?** COG fractions are continuous (non-binary) responses, so the Gaussian phylogenetic linear model is the appropriate choice. The logistic_MPLE model used elsewhere in the paper applies to binary traits only.

##### S6.3 Visualisation: relative percentage change

For plotting, model coefficients are back-transformed from z-scored units to percentage points and then expressed as **relative change from the prototrophic baseline**:

$$\Delta_{c,b}^{\text{rel}} = \frac{\hat{\delta}_{c,b}}{\hat{\mu}_c} \times 100\%$$

where $\hat{\delta}_{c,b}$ is first rescaled to percentage points by multiplying by $\text{SD}_c$ (the cross-genome standard deviation of COG fraction $c$ before z-scoring), and $\hat{\mu}_c$ is the back-transformed intercept (prototrophic baseline in %). A value of +20, for example, means the COG fraction is 20% higher than in fully prototrophic genomes. The zero line in each facet is annotated with $\hat{\mu}_c$ to provide the absolute context.

**Compositionality note.** COG fractions sum to approximately 100% per genome, so they are compositional data. Because the fractions are the *response* variable (not predictors), this does not cause collinearity or bias within any individual model. However, a positive shift in one COG category necessarily implies compensatory negative shifts in others on average. Results from different COG facets should therefore be interpreted jointly rather than in isolation. Centred log-ratio (CLR) transformation was not applied to the response because it changes the quantity being modelled from an absolute fraction to a log-ratio, complicating biological interpretation of percentage-point shifts; z-scoring prior to model fitting controls for scale differences without altering the additive structure of the linear predictor.

---

#### S7. Co-Auxotrophy Associations: Observed vs. Expected

##### S7.1 Independence null

For each pair of amino acids $(a, b)$, the expected co-auxotrophy frequency under the independence assumption is:

$$E_{ab} = p_a \cdot p_b$$

where $p_a = \Pr(\text{auxotrophic for } a)$ is the marginal auxotrophy frequency estimated from the data. For triples $(a, b, c)$:

$$E_{abc} = p_a \cdot p_b \cdot p_c$$

The observed co-auxotrophy frequency is the proportion of genomes simultaneously auxotrophic for all members of the group.

##### S7.2 Regression and null hypothesis tests

Observed frequencies were regressed on expected frequencies using ordinary least squares:

$$O_{ab} = \alpha_0 + \alpha_1 E_{ab} + \epsilon_{ab}$$

Under perfect independence, $\alpha_0 = 0$ and $\alpha_1 = 1$. Two one-sample t-tests were performed:

$$H_0 : \alpha_0 = 0 \qquad t_0 = \frac{\hat{\alpha}_0}{\mathrm{SE}(\hat{\alpha}_0)}, \quad df = n_{\mathrm{pairs}} - 2$$

$$H_0 : \alpha_1 = 1 \qquad t_1 = \frac{\hat{\alpha}_1 - 1}{\mathrm{SE}(\hat{\alpha}_1)}, \quad df = n_{\mathrm{pairs}} - 2$$

Both use two-sided p-values from the t-distribution. A slope $\hat{\alpha}_1 > 1$ indicates that co-auxotrophies accumulate faster than expected under independence (positive synergy among auxotrophies).

These regressions were computed globally and stratified by broad environment class (animal, aquatic, soil, plant) and by phylum (Proteobacteria, Firmicutes_A, Bacteroidota).

> **Correctness note.** The OLS regression of observed on expected co-auxotrophy frequencies assumes that expected frequencies are measured without error. In practice, both $O_{ab}$ and $E_{ab}$ are estimated from the same dataset, introducing correlated sampling error. The regression therefore slightly underestimates slope uncertainty. This approach is standard in the microbiome co-occurrence literature and is appropriate here given the large sample sizes, but should be noted as an approximation.
>
> **Correctness note on phylogenetic non-independence.** The doubles/triples analysis uses raw observed frequencies without phylogenetic correction. Genomes are not independent observations, so the regression standard errors are likely underestimated. This analysis is best treated as an exploratory description of co-auxotrophy patterns rather than a formal statistical test.

---

#### S8. Auxotrophy Identity Predicted by Total Auxotrophy Count

##### S8.1 Phylogenetic GLMM

**Leave-one-out (LOO) predictor.** Naively using total auxotrophy count $A_i$ as the predictor for amino acid $a$'s loss introduces circular self-prediction: a genome that is auxotrophic for $a$ has $A_i$ inflated by 1 relative to an otherwise identical genome that is prototrophic for $a$. To remove this circularity, the predictor for each per-AA model is the *other-auxotrophies* count:

$$O_{i,a} = A_i - \mathbf{1}[\mathrm{AA}_{i,a} = 0]$$

That is, for a genome that is auxotrophic for $a$, its own auxotrophy is subtracted; for a genome that is prototrophic for $a$, $O_{i,a} = A_i$. This ensures that the predictor is never inflated by the response variable being modelled.

**Inclusion criterion.** All genomes are included, including those with $O_{i,a} = 0$ (i.e., fully prototrophic genomes and genomes auxotrophic only for the focal amino acid). The "0" bin contains both prototrophs ($O_{i,a} = 0$ because $A_i = 0$) and genomes auxotrophic only for $a$ ($O_{i,a} = 0$ because $A_i = 1$ and the one loss is $a$ itself). This bin has genuine variation in the response and provides a meaningful baseline: the estimated probability of losing amino acid $a$ in a genome that has no other amino acid losses. Bins in which all genomes share the same auxotrophy status for the focal amino acid (monomorphic bins) are excluded before fitting, as no coefficient can be estimated from a response with no variation.

For each amino acid, a phylogenetic generalised linear mixed model (PGLMM; `phyr::pglmm`) was fitted to test whether the LOO auxotrophy count predicts which specific amino acids are lost:

$$\mathrm{logit}(P(\text{auxotrophic for } a)_i) = \mu_a + \sum_{b \neq \mathrm{ref}} \beta_{a,b} \cdot \mathbf{1}[B'_{i,a} = b] + u_i$$

where $B'_{i,a}$ is the binned LOO count (paired bins of width 2, with bin "0" as the reference level):

$$B'_{i,a} \in \{0,\ 1\text{–}2,\ 3\text{–}4,\ 5\text{–}6,\ 7\text{–}8,\ 9\text{–}10,\ 11\text{–}12,\ 13+\}$$

and $u_i \sim \mathcal{N}(0, \sigma_u^2 \boldsymbol{C})$ is a phylogenetic random intercept with covariance matrix $\boldsymbol{C}$ derived from the species tree.

##### S8.2 Log2 enrichment ratio

To compare enrichment profiles across amino acids with different global loss frequencies, raw PGLMM-predicted probabilities were converted to log2 enrichment ratios:

$$L_{a,b} = \log_2\left(\frac{\hat{p}_{a,b}}{g_a}\right)$$

where $\hat{p}_{a,b}$ is the back-transformed predicted probability of auxotrophy for amino acid $a$ in bin $b$ (obtained by inverting the logit):

$$\hat{p}_{a,b} = \frac{1}{1 + e^{-(\hat{\mu}_a + \hat{\beta}_{a,b})}}$$

(with $\hat{\beta}_{a,\text{ref}} = 0$ for the reference bin "0"), and $g_a$ is the global (unconditional) auxotrophy prevalence for amino acid $a$ across all genomes in the LOO-filtered dataset for that AA. Values $L_{a,b} > 0$ indicate that amino acid $a$ is lost more often than expected at count bin $b$; values $< 0$ indicate depletion. Infinite values (arising when $\hat{p}_{a,b} = 0$) are replaced with 0.

##### S8.3 Significance testing: likelihood ratio test

A LRT was used to test whether the bin predictor significantly improves model fit over a phylogenetic intercept-only null for each amino acid:

$$\Lambda_a = -2\left(\hat{\ell}_{\text{null},a} - \hat{\ell}_{\text{full},a}\right) \sim \chi^2_{(K_b-1)}$$

where $K_b = 7$ is the number of bins $(1\text{–}2, 3\text{–}4, \ldots, 13+)$, so $df = 6$. Null and full models are fitted on identical observations (same LOO-filtered genome set per AA) to ensure valid log-likelihood comparison. The log-likelihoods $\hat{\ell}$ are taken directly from the `$logLik` field of the fitted `pglmm` objects (phyr stores this as a scalar; the `logLik()` S3 generic is not dispatched for objects of class `communityPGLMM`).

**AIC for pglmm objects.** Because `AIC()` does not dispatch on `pglmm` objects, AIC was computed manually:

$$\text{AIC} = -2\,\hat{\ell} + 2\,K_p$$

where $K_p = \text{nrow}(\boldsymbol{B})$ is the number of rows in the fixed-effects coefficient matrix. Note that this counts only fixed-effect parameters; the phylogenetic variance parameter ($\sigma_u^2$) is not included in $K_p$.

**Correctness note on AIC parameter count.** The complete AIC formula should count all free parameters including variance components. For pglmm, the random effect variance is stored in `$ss` (square-root parameterisation). The full count would be $K_p = \text{nrow}(\boldsymbol{B}) + \text{length}(\hat{\sigma})$. However, because the full and null models share the same random-effect structure (one phylogenetic variance component), the $\text{length}(\hat{\sigma})$ term cancels in $\Delta\text{AIC} = \text{AIC(null)} - \text{AIC(full)}$. **The ΔAIC values used for the wildcard analysis are therefore unaffected by this simplification.** The absolute AIC values (printed in `sig_table`) would increase by 2 per model if the variance component were included, but significance and ΔAIC conclusions are unchanged.

##### S8.4 Wildcard quantification

Two complementary measures were computed to characterise each amino acid's relationship to total auxotrophy count:

**ΔAIC** (from §S8.3): higher values indicate that the total count bin is a stronger predictor of that amino acid's loss probability ("orderly" losses).

**Enrichment fold-range:**

$$\text{FoldRange}_a = 2^{\max_b L_{a,b} - \min_b L_{a,b}} = \frac{\max_b \hat{p}_{a,b}}{\min_b \hat{p}_{a,b}}$$

i.e. the ratio of the highest to the lowest predicted probability across all bins, equivalently the span of the $\log_2$ enrichment profile exponentiated to a probability ratio. Higher fold-range indicates a more vivid enrichment profile — strong enrichment at high count bins and depletion at low bins. A fold-range of 1 means the probability is identical across all bins (flat profile). Amino acids with both high ΔAIC and high fold-range show the most "orderly" loss pattern tightly coupled to total auxotrophy burden; those with low values on both metrics are lost largely independently of count ("wildcards").

**Enrichment range.** For the enrichment-range visualisation (Figure 5C), each amino acid's enrichment range is the span of log2 enrichment from the lowest to the highest bin:

$$\text{Range}_a = L_{a,\,13+} - L_{a,\,1\text{–}2}$$

This is plotted as a horizontal segment connecting the log2 enrichment at bin "1–2" (left endpoint, blue) to the log2 enrichment at bin "13+" (right endpoint, red). A large positive range indicates that the amino acid is strongly enriched for loss in highly streamlined genomes but relatively protected at low auxotrophy burden.

**Crossover point.** The crossover bin is the lowest bin $b^*$ at which the log2 enrichment first reaches or exceeds zero (i.e., the amino acid is at least as likely to be lost as the global average):

$$b_a^* = \min\{b : L_{a,b} \geq 0\}$$

If no bin reaches zero, the amino acid is considered enriched at all count levels examined. The crossover point is annotated as a filled downward-pointing triangle (▼) on the heatmap.

---

#### S9. Environment ΔAIC in Env-Only vs. Multivariate Context

To test whether the apparent environment signal is mediated through genomic composition (COG fractions), we compared the environment ΔAIC in two modelling contexts:

$$\Delta\text{AIC}_{\text{env-only}} = \text{AIC}(\text{intercept-only}) - \text{AIC}(\text{env-only phyloglm})$$

$$\Delta\text{AIC}_{\text{multivariate}} = \text{AIC}(\text{full} - \text{broadclass}) - \text{AIC}(\text{full phyloglm})$$

The difference (ΔΔAIC) quantifies how much the environment's predictive contribution changes once COG composition is controlled:

$$\Delta\Delta\text{AIC} = \Delta\text{AIC}_{\text{multivariate}} - \Delta\text{AIC}_{\text{env-only}}$$

A negative ΔΔAIC means environment explains **less** once COG composition is accounted for, suggesting that some of the apparent environment signal is mediated by (or confounded with) variation in gene content. A positive ΔΔAIC means environment and COG composition are complementary — each carries unique information.

> **Correctness note on sample scope.** The environment-only models (script 03) were fitted on all high-quality MAGs using all four environment classes (or three for L-Glutamine). The multivariate models (script 05) were also fitted on MAGs but after further filtering to genomes with COG data (`na.omit()` on the merged table). The two ΔAIC values are therefore computed on slightly different genome sets. The comparison is approximate. A stricter like-for-like comparison would refit the environment-only models on the same COG-filtered genome set. This is flagged as a caveat in the script (`09_compare_environment_aic.Rmd`, "Note on sample scope") but the current analysis does not implement the refit.

#### S10. General Statistical Notes

**Tree rescaling.** All phylogenetic models used a bac120 phylogeny rooted on Fusobacteria. Prior to fitting, branch lengths were rescaled to unit tree height:

$$d'_{ij} = \frac{d_{ij}}{\max_k \operatorname{depth}(k)}$$

This ensures that the $\alpha$ parameter (and derived $t_{1/2}$) is on a comparable [0,1] scale, and avoids numerical issues from very large or very small branch lengths.

**Multiple testing.** Unless otherwise stated, p-values from multiple model comparisons or coefficient tests within an analysis were adjusted using the Benjamini–Hochberg FDR procedure. Significance was declared at FDR-adjusted $p < 0.05$.

**Model convergence.** `phyloglm` with `logistic_MPLE` can encounter a "btol boundary" warning when the linear predictor approaches the numerical boundary during optimisation. All models tracked this warning; affected models should be treated as conservative (coefficient magnitudes are lower bounds). Convergence status is recorded in `convergence_log` (script 03) and the `BtolReached` column (scripts 05 and 06).

---

#### S12. Leave-One-Class-Out Cross-Validation: Environment Signal (AUC)

To assess whether the environment → auxotrophy signal generalises beyond the lineages used to estimate it, we performed leave-one-class-out (LOCO) cross-validation at the GTDB class level (script `11_loco_env_auc.Rmd`). Classes with $\geq 50$ MAGs in the environment-model dataset were eligible as holdouts; classes below this threshold were excluded to ensure stable phyloglm estimation on training data.

For each held-out class:

1. The phyloglm model (broadclass predicting auxotrophy, `logistic_MPLE`, sum-to-zero contrasts) was refitted on all genomes from the remaining classes.
2. Predicted auxotrophy probabilities were computed for the held-out class using fixed effects only — the intercept and environment coefficients. The phylogenetic error term $\varepsilon_i$ is specific to the tips used for fitting and cannot be extrapolated to out-of-sample taxa.
3. Area under the ROC curve (AUC) was computed against observed auxotrophy status using the `pROC` package. Because auxotrophy is coded 0 (auxotrophic) and 1 (prototrophic), predicted probabilities were inverted ($P(\text{auxo}) = 1 - \hat{p}$) before computing AUC.

AUC $> 0.5$ indicates that environment coefficients estimated from the rest of the tree predict the direction of auxotrophy variation in the held-out class better than chance. AUC $\approx 0.5$ indicates the environment signal does not transfer out-of-sample. L-Glutamine was modelled with three environment levels (animal, aquatic, soil) as in the main analysis.

Results are summarised per amino acid (mean, median, min, max AUC across held-out classes) and shown in Supplementary Figure M.

---

#### S13. Leave-One-Class-Out Cross-Validation: COG Directional Concordance

To test whether the auxotrophy burden $\rightarrow$ COG composition shifts generalise across lineages, we performed LOCO cross-validation of the phylolm COG-shift models at the GTDB class level (script `12_loco_cog_directional.Rmd`). The same eligibility threshold of $\geq 50$ MAGs per class was applied.

For each held-out class:

1. COG fractions were computed and z-scored within the training set only (mean and SD from training genomes, applied to scale both training and held-out genomes).
2. `phylolm` (Pagel's $\lambda$) was refitted for each COG category with auxotrophy burden bin as a categorical predictor (reference = bin "0", prototrophic genomes). BH-adjusted p-values were computed across all bin terms for each COG.
3. Significant (COG $\times$ bin) pairs were identified (BH-adjusted $p < 0.05$). For each significant pair, the sign of the training coefficient was recorded.
4. In the held-out class, the raw mean COG fraction was compared between the focal burden bin and bin "0". Pairs were skipped if either bin had fewer than 5 genomes in the held-out class.
5. A pair was scored as **concordant** if the sign of the held-out mean difference matched the sign of the training coefficient, and **discordant** otherwise.

Directional concordance is reported as the fraction of testable (COG $\times$ bin $\times$ class) combinations that are concordant. Because this approach tests each significant bin separately rather than summarising to a single trend direction, it correctly handles COG categories whose shifts are restricted to specific burden levels rather than monotonically increasing or decreasing across the full range.

Results are summarised per class, per COG category, and per burden bin, and shown in Supplementary Figures N and O.

---

#### S14. Jackknife Confidence Intervals for $R^2$_lik Variance Partitioning

To quantify uncertainty in the $R^2$_lik variance partitioning shown in Figure 3c, we computed 95% confidence intervals using a delete-one-GTDB-class jackknife (script `13_r2_jackknife.Rmd`). The same set of eligible classes used in the LOCO cross-validations ($\geq$ 50 MAGs; n = 13 classes) served as jackknife units.

For each held-out class, the class's genomes were removed from the dataset and the tree was pruned accordingly. Three models were then refitted on the remaining genomes for each amino acid:

1. **Full phyloglm** — `AA ~ broadclass` with phylogenetic correction (logistic_MPLE)
2. **Intercept-only phyloglm** — `AA ~ 1` with phylogenetic correction
3. **Intercept-only GLM** — `AA ~ 1` without phylogenetic correction (null model)

All three models within each replicate were fitted on an identical genome subset, ensuring that $R^2$_lik comparisons are valid (matched observations). $R^2$_lik was then computed as:

- **$R^2$_total** = $R^2$_lik(full phyloglm, intercept GLM)
- **$R^2$_phylo** = $R^2$_lik(intercept phyloglm, intercept GLM)
- **$R^2$_env** = $R^2$_total − $R^2$_phylo

This yielded a 15 × 16 matrix of jackknife estimates for each component. Jackknife standard errors were computed as:

$$\mathrm{SE}_{\mathrm{jack}} = \sqrt{\frac{n-1}{n}\sum_{i=1}^{n}(\hat{\theta}_i - \bar{\theta})^2}$$

and 95% confidence intervals were derived using the t-distribution with n − 1 = 14 degrees of freedom. The full-data $R^2$_lik values (fitted on all genomes, from script 03) are used as point estimates in Figure 3c; the jackknife CIs are overlaid as error bars to show sensitivity to clade composition.

---

#### S11. Software and Packages

| Software | Version | Purpose |
|---|---|---|
| R | v4.3.1 | FBA simulations (auxotrophy prediction) |
| R | v4.4.1 | All phylogenetic and statistical analyses |

| Software | Version | Purpose |
|---|---|---|
| gapseq | v1.2 (commit e3210c0f) | Genome-scale metabolic model reconstruction |
| sybil | v2.2.1 | Flux balance analysis (FBA) in R |
| phylolm | v2.6.5 | Phylogenetic linear and logistic regression (phyloglm, logistic_MPLE; phylolm, Pagel's $\lambda$) |
| phyr | v1.1.3 | Phylogenetic GLMM (pglmm, binomial family) |
| rr2 | v1.1.1 | Likelihood-based partial $R^2$ ($R^2$_lik) |
| ape | v5.8-1 | Tree manipulation, cophenetic distances |
| tidyverse | v2.0.0 | Data wrangling |
| ggplot2 | v3.5.0 | Figure production |
| svglite | v2.2.2 | SVG figure output with editable text |

**References:** Ho LST & Ané C (2014) A linear-time algorithm for Gaussian and non-Gaussian trait evolution models. *Systematic Biology* 63:397–408.

### Supplementary Figures

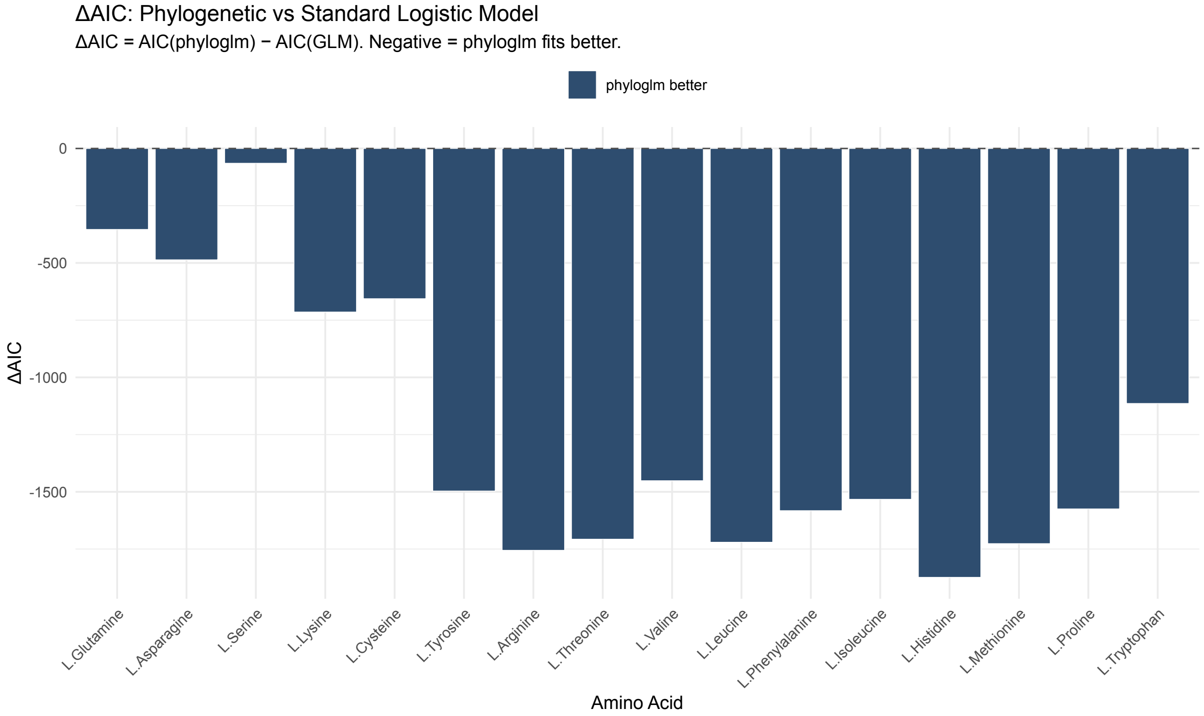


**Figure S1.** Phylogenetic correction improves model fit across amino acids. ΔAIC (AIC[GLM] − AIC[phyloglm]) per amino acid. Negative values indicate that the phylogenetically corrected model fits substantially better than the uncorrected GLM, justifying the use of phyloglm throughout. All amino acids show negative ΔAIC, confirming that phylogenetic correction is necessary for valid inference.

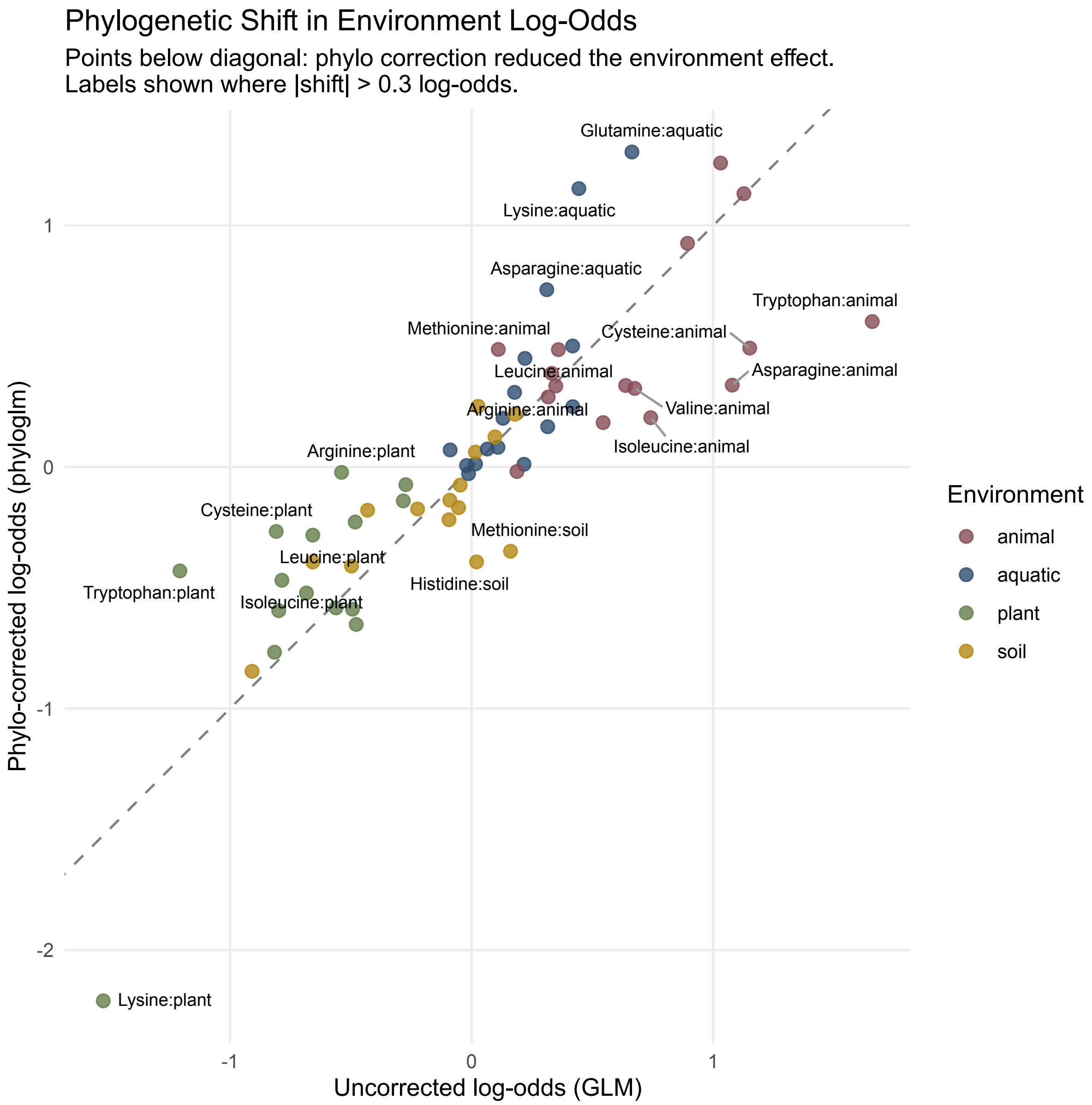


**Figure S2.** Phylogenetic correction alters estimated environment effects. Scatter of per-environment log-odds estimates from GLM ($x$-axis) versus phyloglm ($y$-axis) for each amino acid–environment pair. Points above the diagonal indicate cases where phylogenetic correction revealed a higher environment effect; points below indicate cases where correction reduced the apparent effect. Dashed line indicates the 1:1 diagonal.

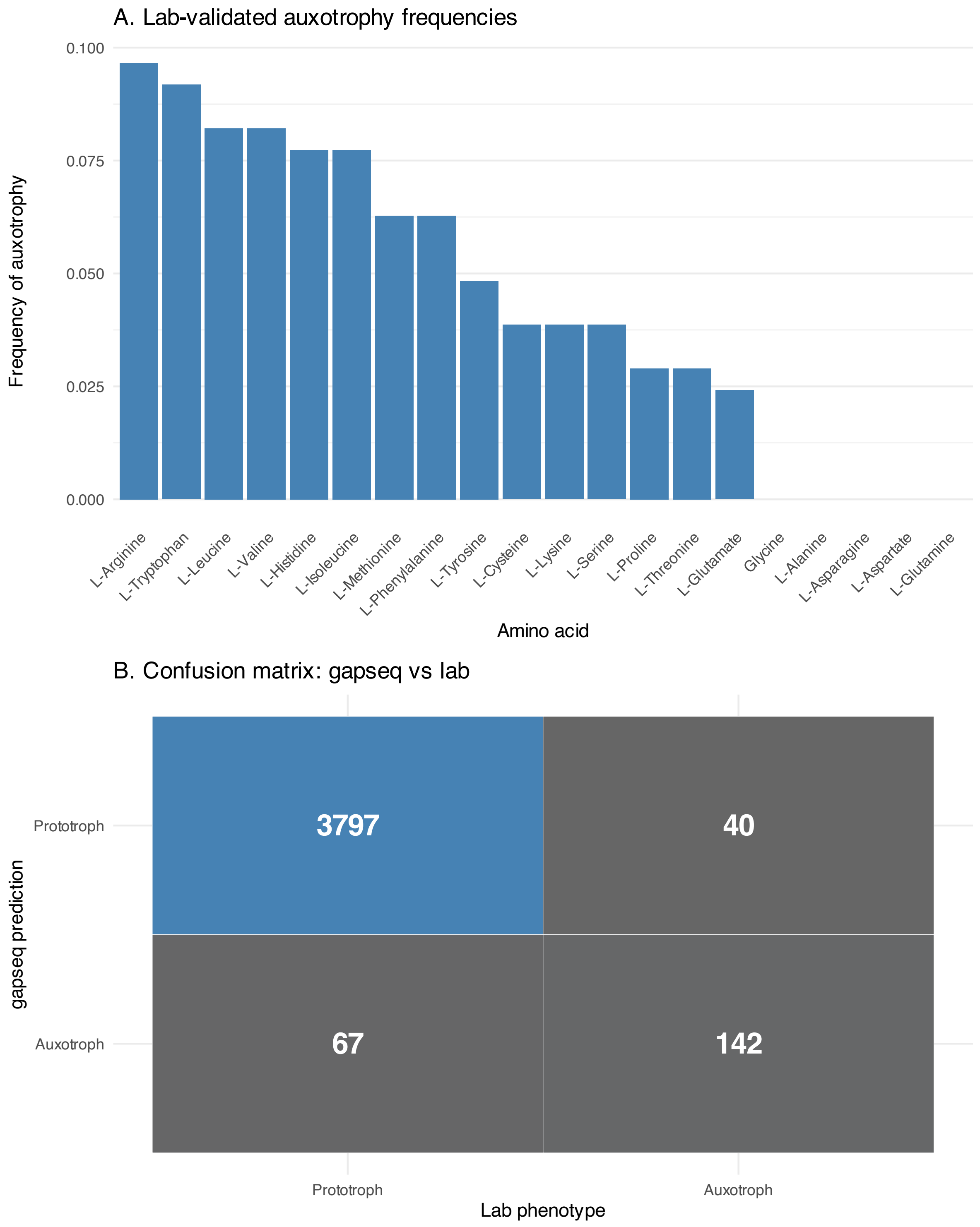


**Figure S3.** Benchmarking gapseq auxotrophy predictions against experimental data. Auxotrophy frequencies in lab-phenotyped strains (Ramoneda et al. 2023; Starke et al. 2025) alongside gapseq predictions for the same genomes. The confusion matrix summarises prediction accuracy across amino acids. False negatives (auxotrophies predicted as prototrophic) are the dominant error mode, consistent with gene-content-based predictions being conservative estimates.

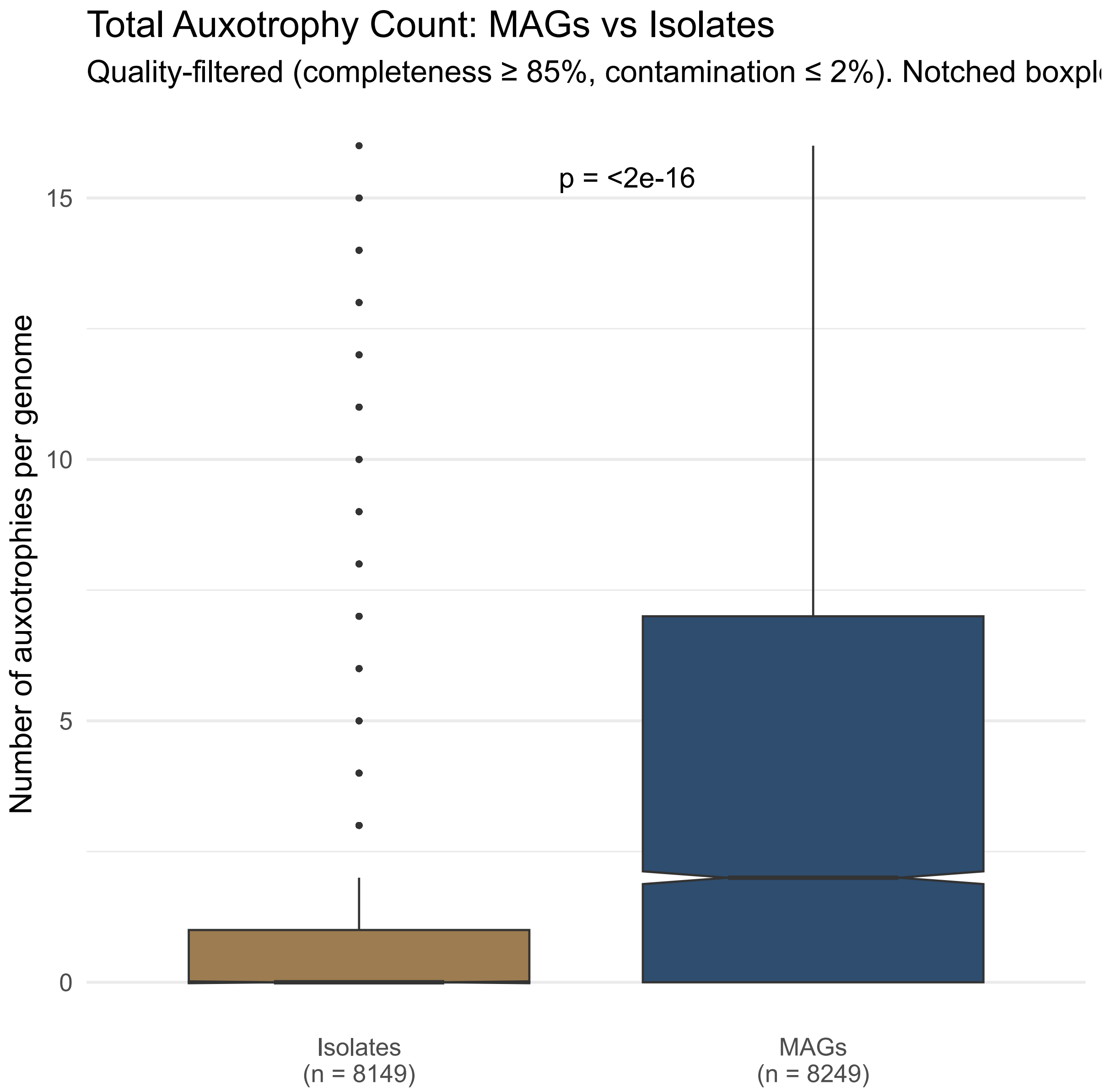


**Figure S4.** Total auxotrophy count: MAGs versus isolates. Boxplot comparing the distribution of total per-genome auxotrophy count (summed across 16 amino acids) between MAGs and isolates. MAGs show consistently higher auxotrophy counts, consistent with systematic underrepresentation of auxotrophic bacteria in culture collections.

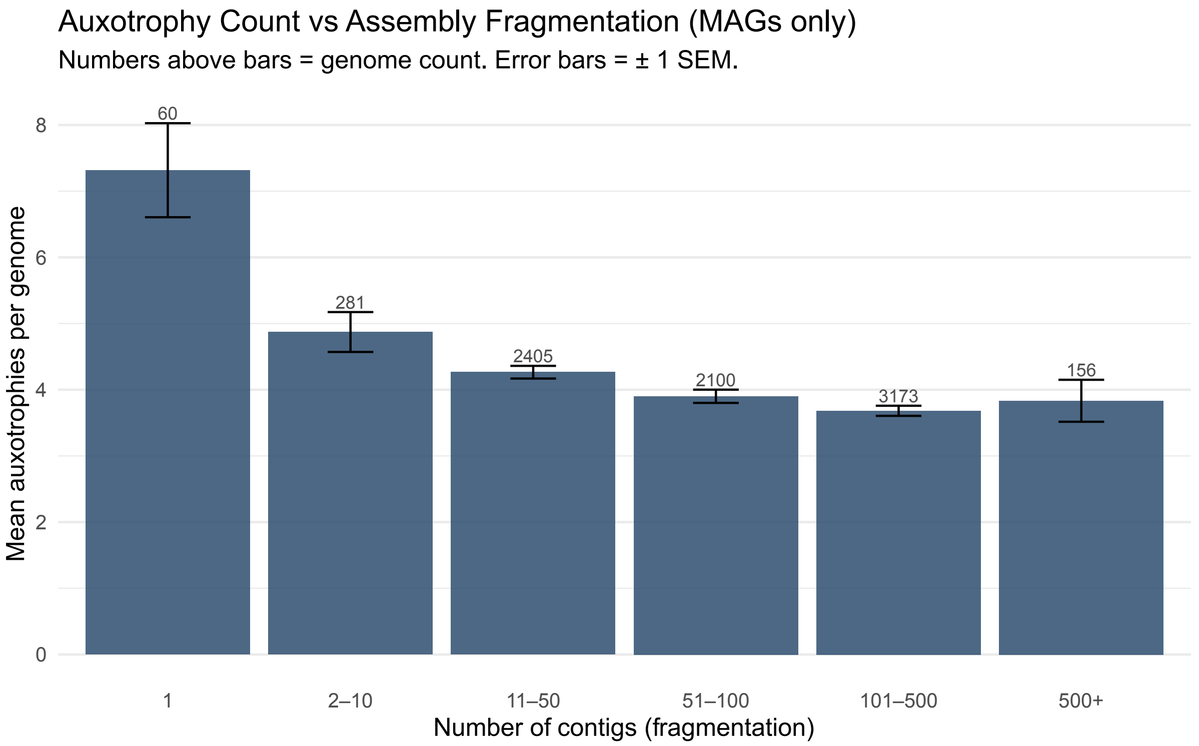


**Figure S5.** Auxotrophy count versus genome fragmentation. Total auxotrophy count versus contig count bins, MAGs only. Quality-control check confirming that auxotrophy inference is robust to assembly fragmentation — highly fragmented assemblies do not show systematically elevated or reduced auxotrophy counts.

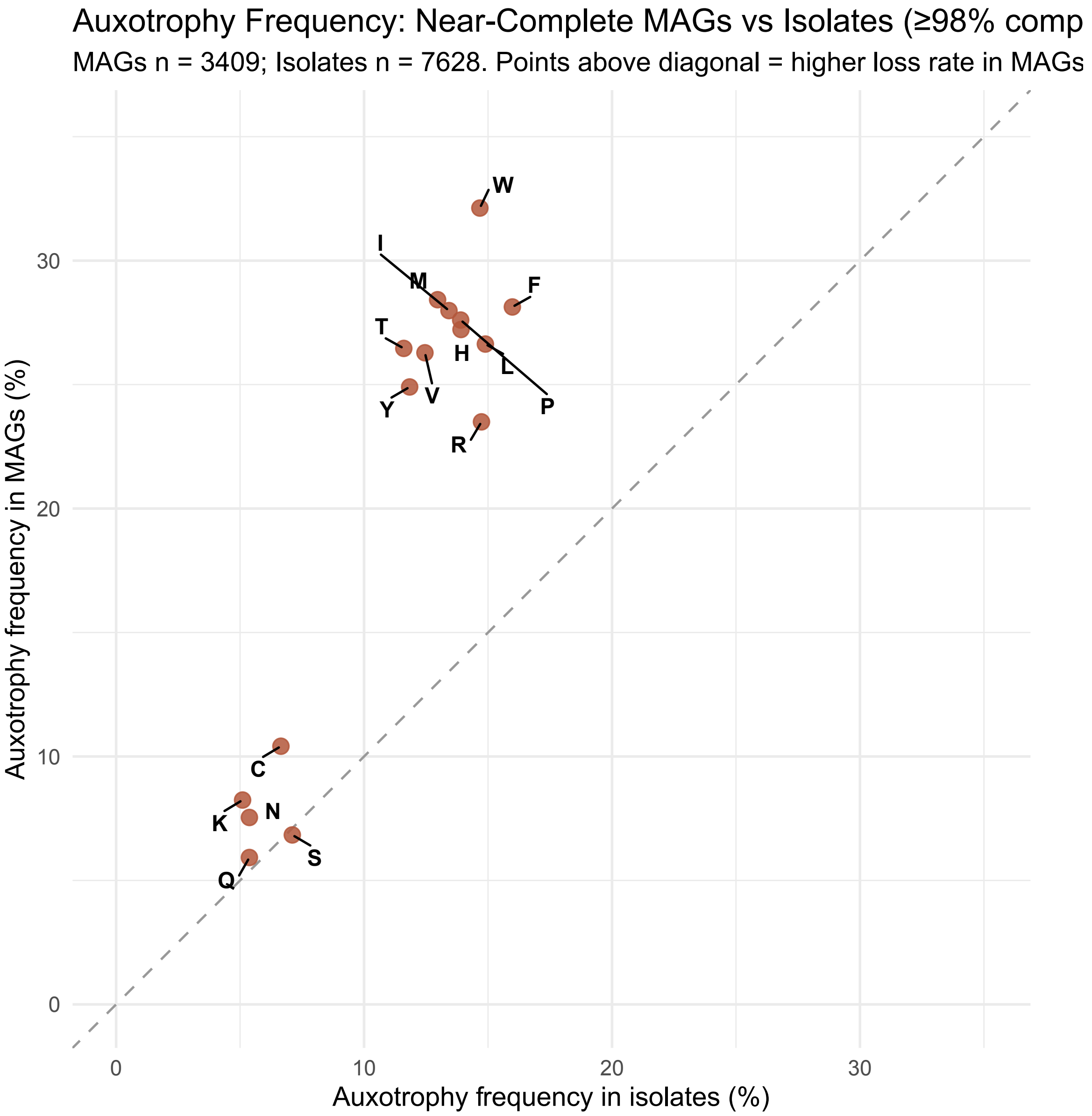


**Figure S6.** Per-amino-acid auxotrophy frequency: near-complete MAGs versus isolates. Per-amino-acid auxotrophy frequency for near-complete MAGs (≥98% completeness) versus near-complete isolates (≥98%). Stringent comparison confirming MAG–isolate concordance at the highest quality threshold, and that the higher auxotrophy rates in MAGs are not an artefact of lower assembly completeness.

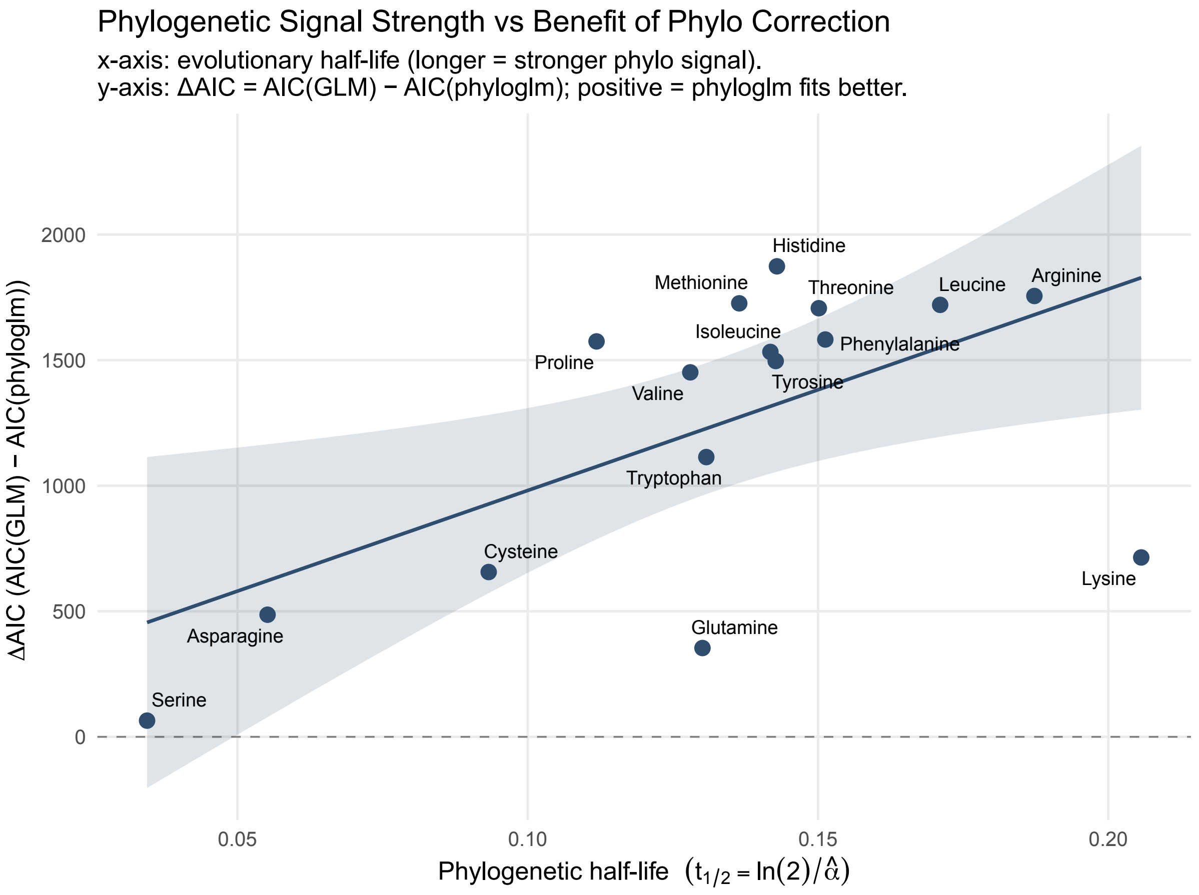


**Figure S7.** Phylogenetic correction benefit versus evolutionary half-life. ΔAIC benefit of phylogenetic correction (AIC[GLM] − AIC[phyloglm]) versus phylogenetic half-life ($t_{1/2}$) per amino acid. Tests whether traits with stronger evolutionary signal (longer half-life) benefit more from phylogenetic correction.

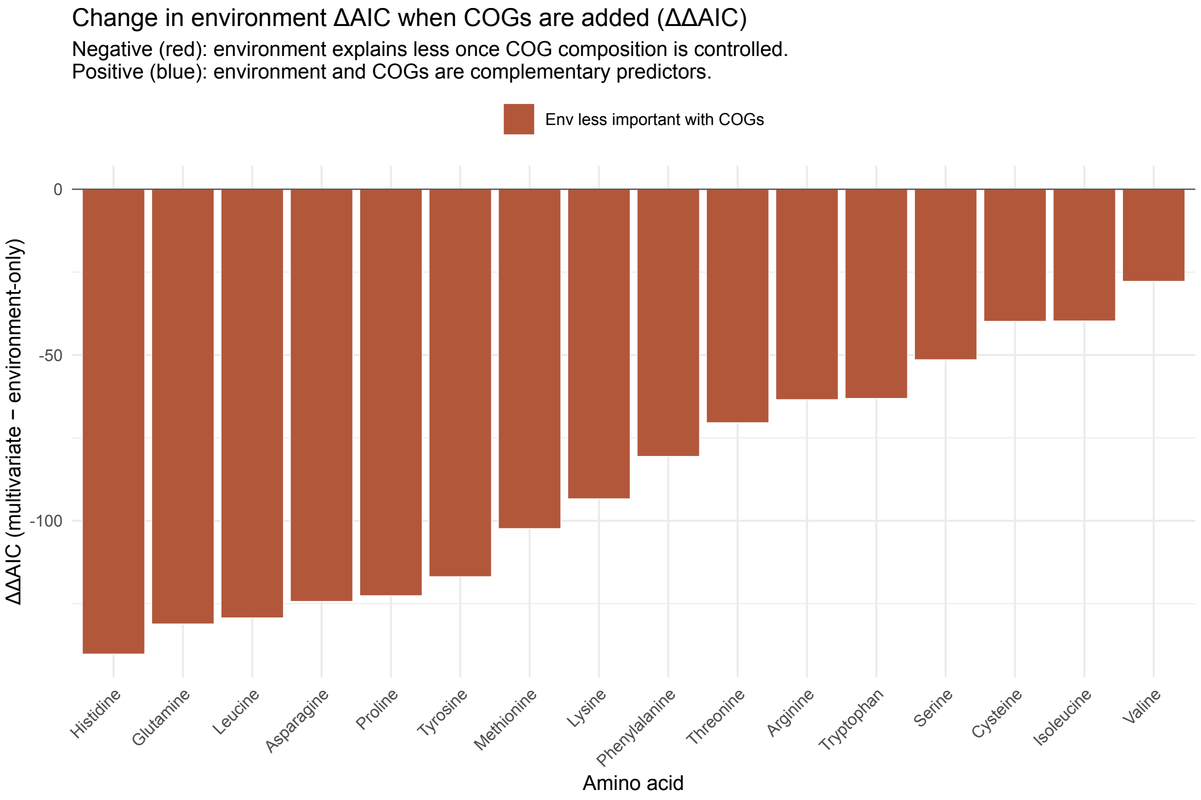


**Figure S8.** Environment ΔAIC: environment-only versus multivariate model. Environment ΔAIC in the environment-only model versus the full multivariate model (broad environment class + COG composition) for each amino acid. Assesses whether the environment signal is mediated by genomic functional composition (broad environment class) — a reduction in ΔAIC in the multivariate context suggests the environment signal is partly confounded with gene content.

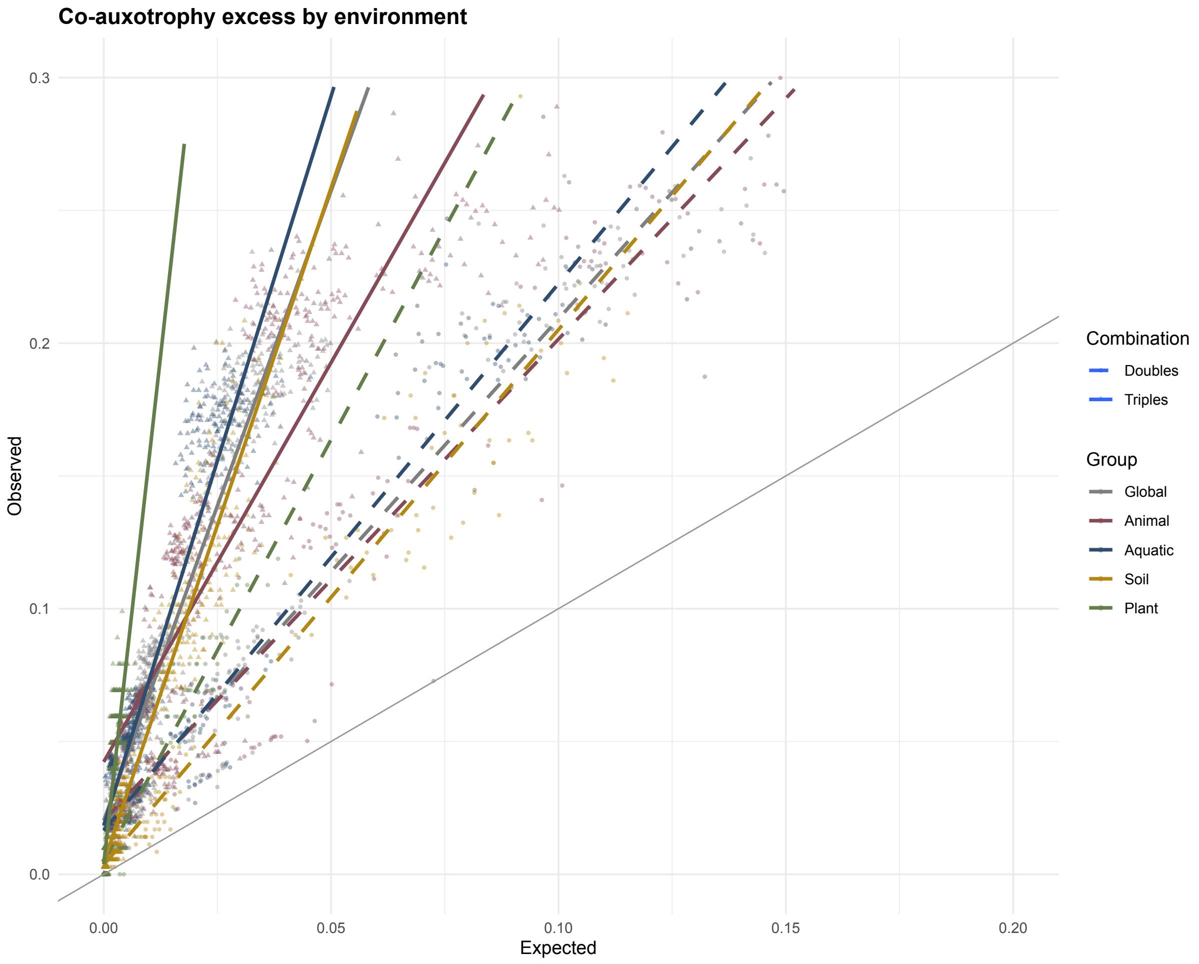


**Figure S9.** Co-auxotrophy doubles and triples by environment. Observed versus expected co-auxotrophy frequencies for doubles (dashed) and triples (solid), with per-environment trend lines shown alongside the global trend. Tests whether the tendency for auxotrophies to co-occur beyond chance differs across broad habitat classes.

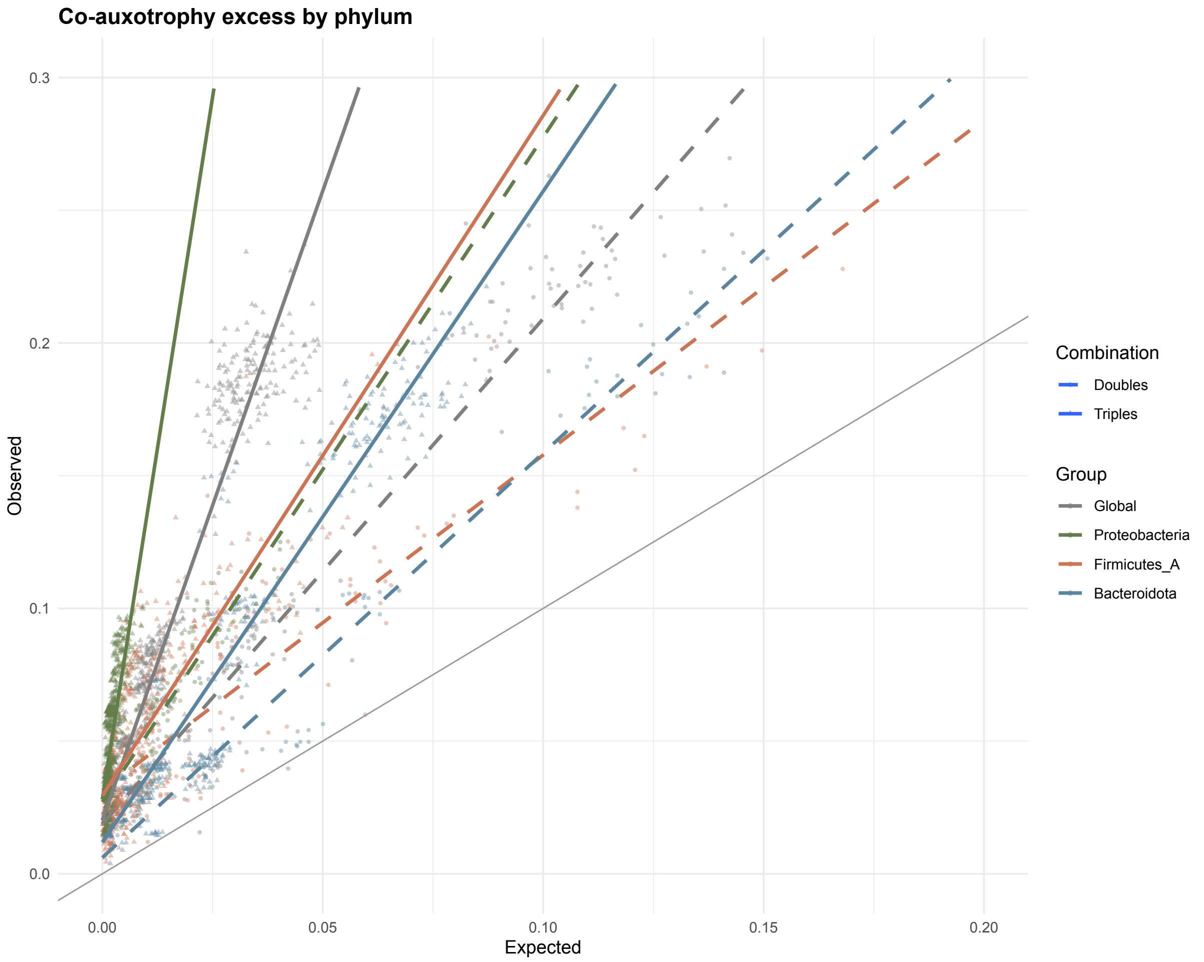


**Figure S10.** Co-auxotrophy doubles and triples by phylum. Observed versus expected co-auxotrophy frequencies for doubles (dashed) and triples (solid), with per-phylum trend lines shown alongside the global trend (Proteobacteria, Firmicutes_A, Bacteroidota). Tests whether the co-auxotrophy excess is consistent across major bacterial phyla.

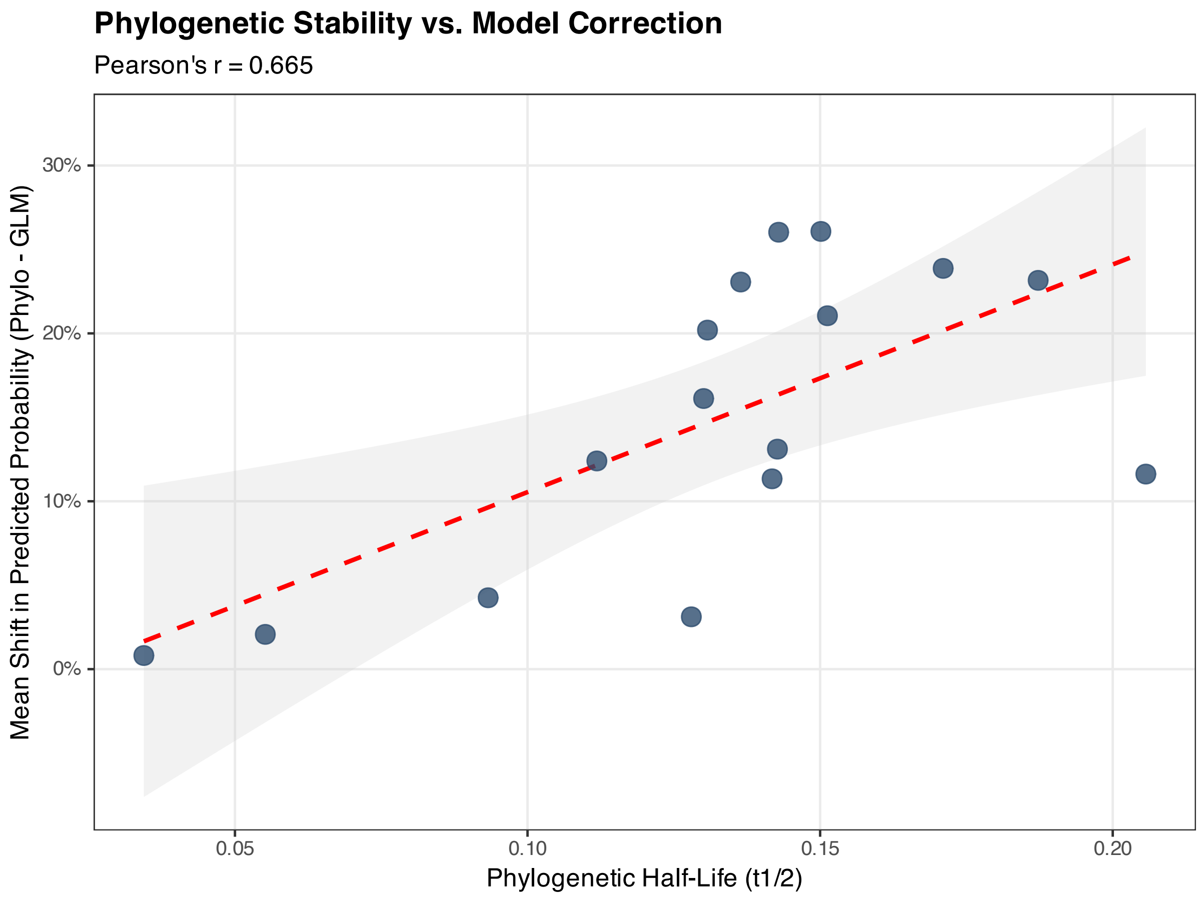


**Figure S11.** Phylogenetic stability versus magnitude of model correction. Phylogenetic half-life ($t_{1/2}$) versus mean shift in predicted auxotrophy probability (phyloglm − GLM) per amino acid. Tests whether evolutionarily more conserved traits (longer half-life) are corrected more by the phylogenetic model, consistent with the expectation that conserved traits are most affected by phylogenetic non-independence.

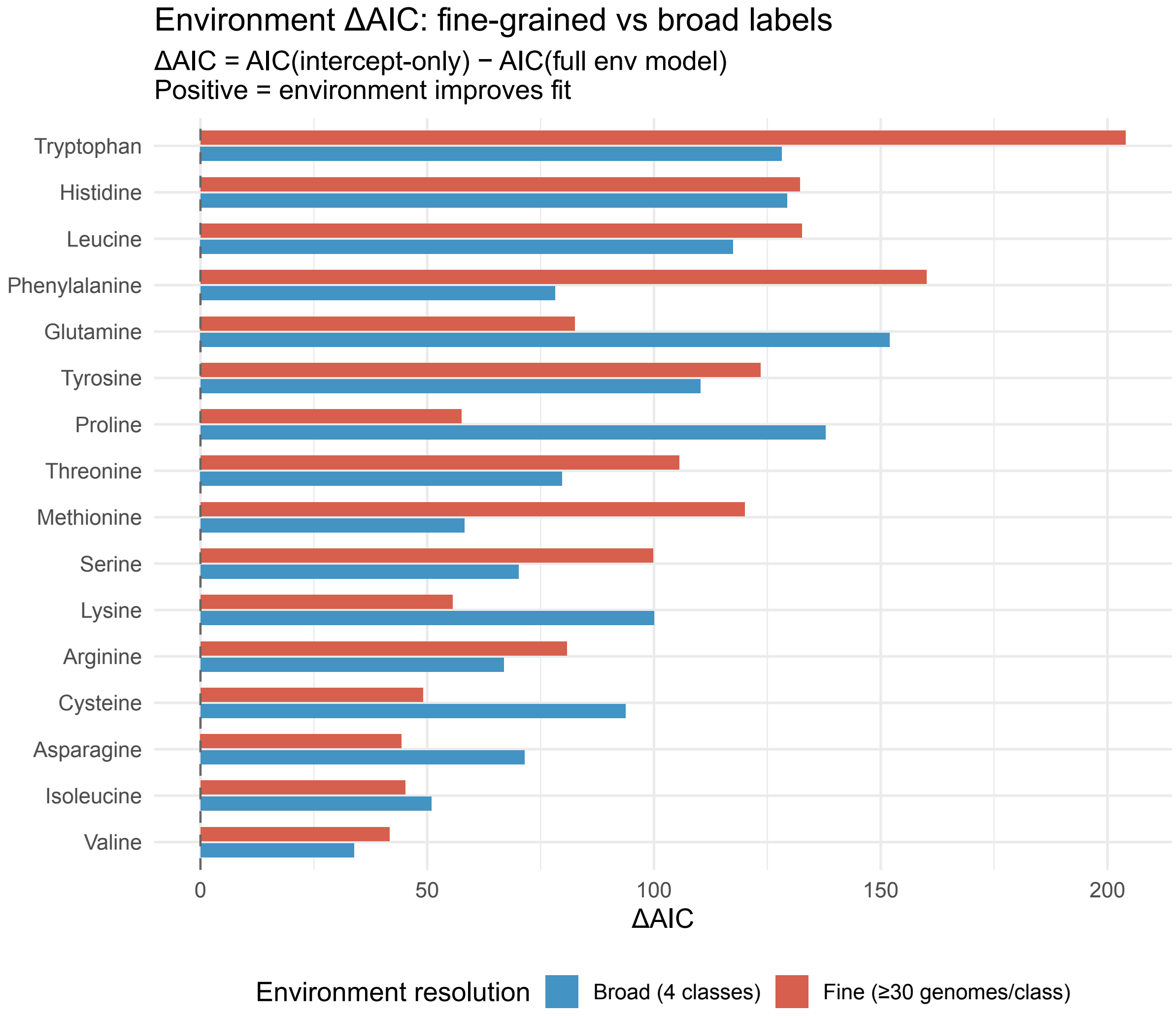


**Figure S12.** Sensitivity analysis: ΔAIC under fine-grained versus broad environment labels. ΔAIC = AIC(intercept-only) − AIC(full environment model) for each amino acid, comparing broad (4-class) and fine-grained (12 classes, ≥30 MAGs each) environment classifications shown side-by-side. Environment showed strong support for all 16 amino acids under both schemes (ΔAIC $> 7$ in all cases), with mean ΔAIC of 92.4 under broad labels versus 95.9 under fine labels. The absence of a uniform improvement with finer resolution argues against systematic attenuation bias in the broad-class analysis. See Supplementary Methods for details.

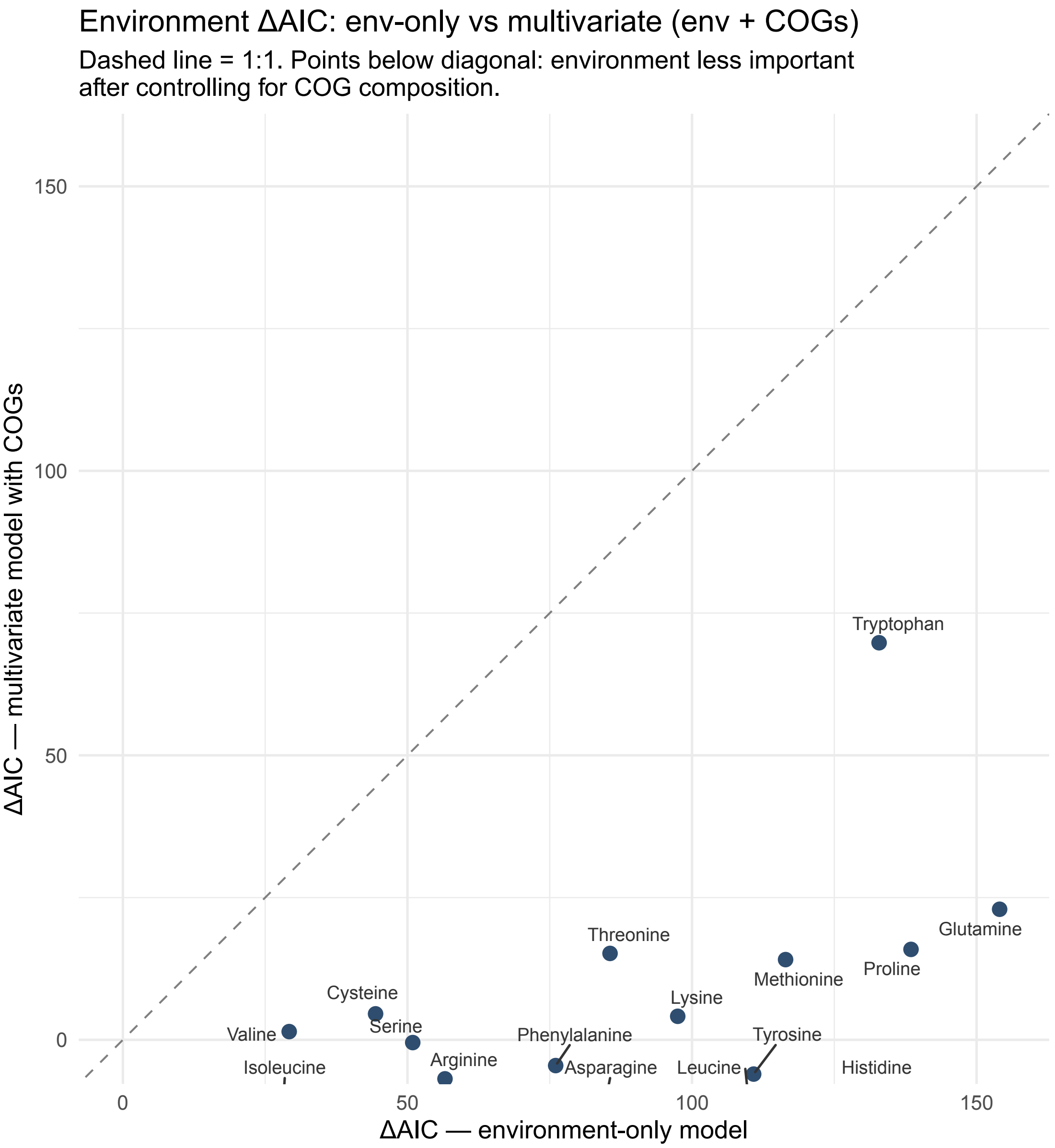


**Figure S13.** LOCO cross-validation: environment predicting auxotrophy (AUC). Leave-one-class-out cross-validation AUC for the environment predicting auxotrophy signal, evaluated at the GTDB class level. Each point represents one held-out class; the crossbar shows the median AUC across held-out classes. The dashed line at AUC = 0.5 indicates chance performance. AUC was computed using fixed-effects-only predictions applied to held-out genomes. See Supplementary Methods.

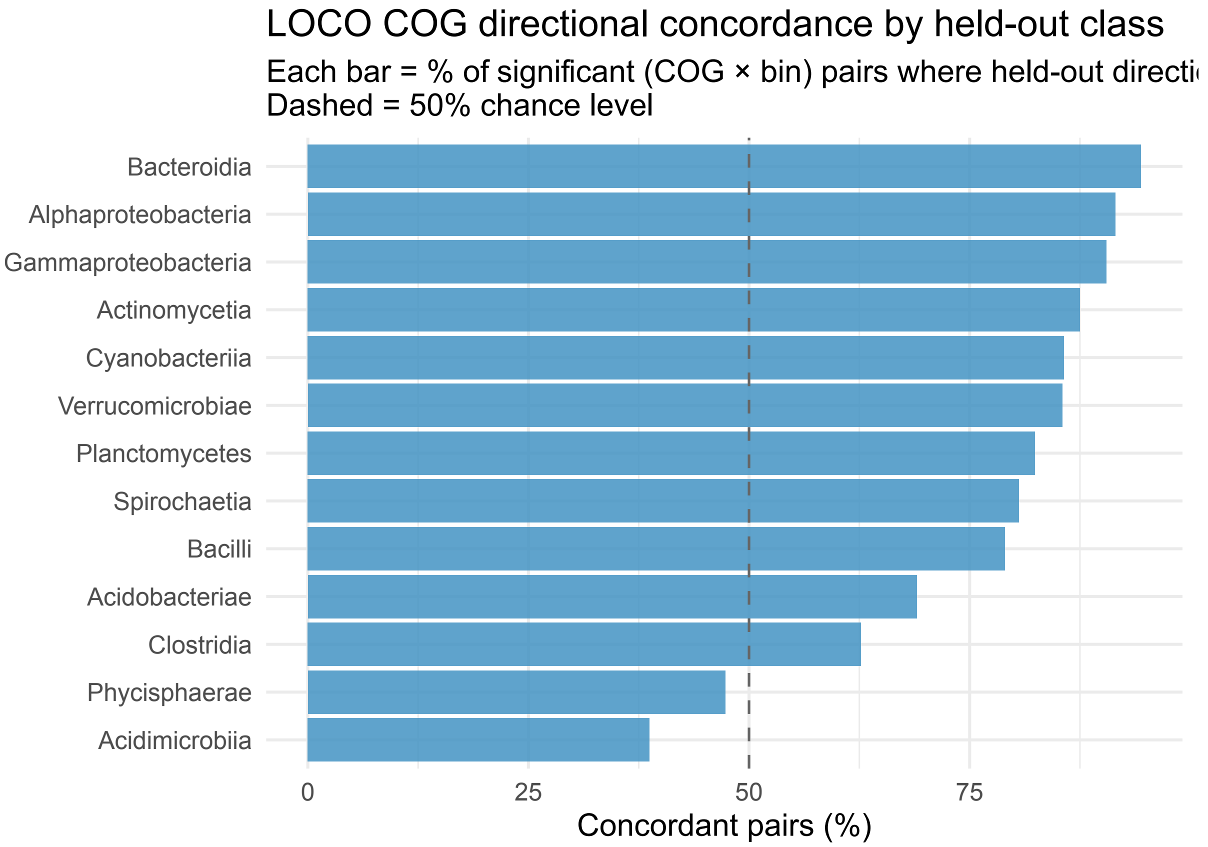


**Figure S14.** LOCO cross-validation: COG directional concordance by class. Percentage of significant (COG category × burden bin) pairs in which the direction of the COG fraction shift in the held-out class matches the sign of the training phylolm coefficient. Each bar represents one held-out class; the dashed line at 50% indicates chance. Overall concordance was 76.7% (493/643 pairs) across 13 held-out classes. See Supplementary Methods.

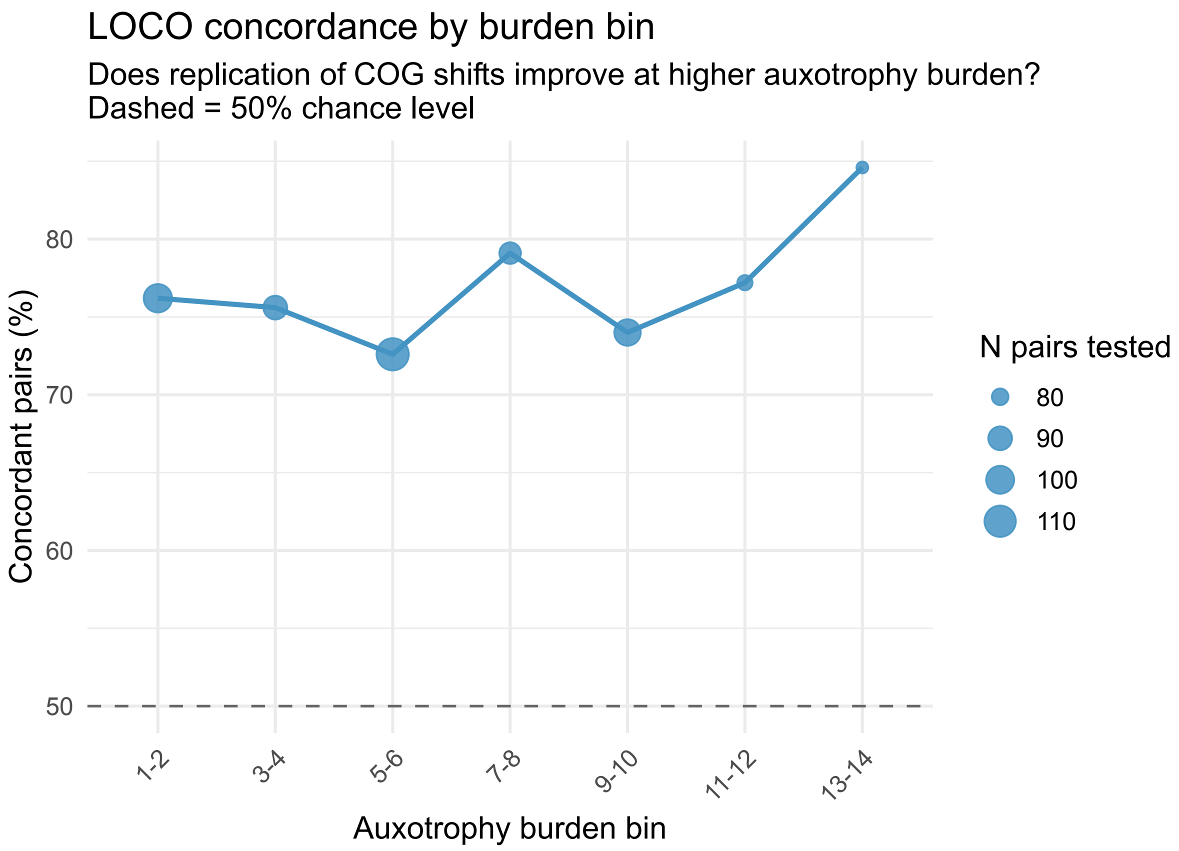


**Figure S15.** LOCO cross-validation: COG directional concordance by burden bin. Percentage of concordant (COG × class) pairs pooled across all held-out classes, shown separately for each auxotrophy burden bin. Point size indicates the number of testable pairs. Concordance is consistently above chance across all bins (73–85%), with the highest replication at bin 13–14 (84.6%). See Supplementary Methods.

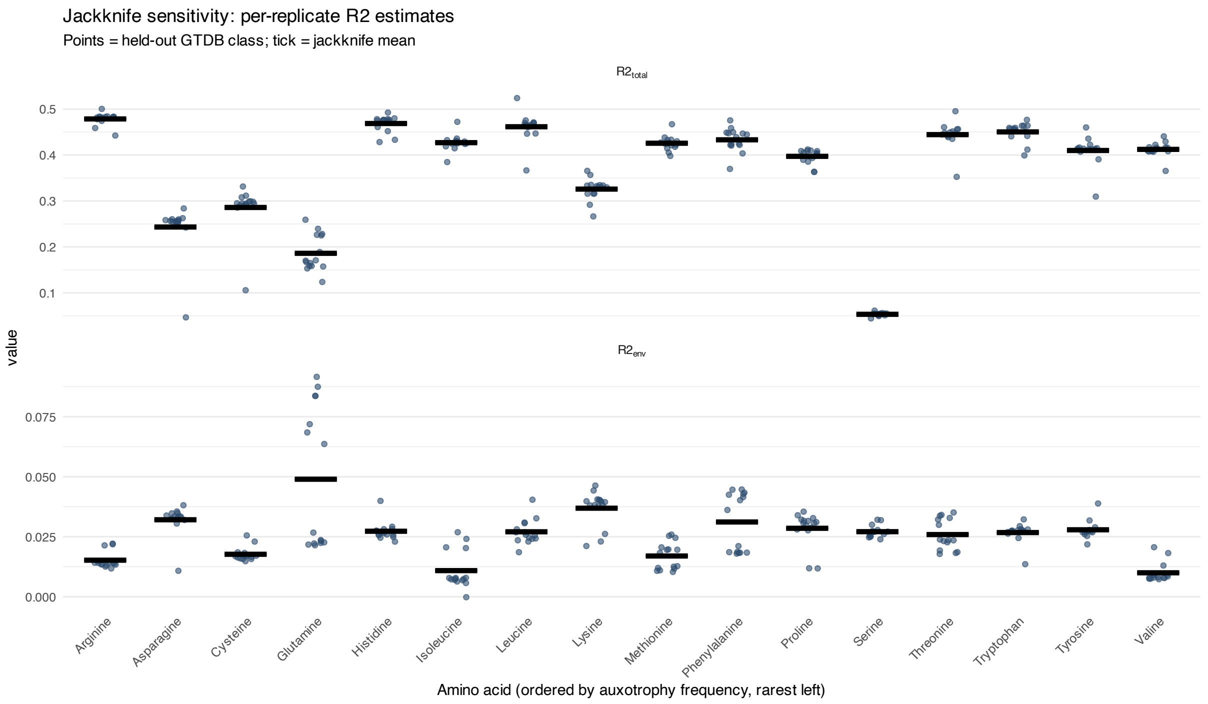


**Figure S16.** Per-replicate $R^2_{\text{lik}}$ estimates from the delete-one-GTDB-class jackknife. $R^2_{\text{lik}}$ for total variance explained ($R^2_{\text{total}}$; upper panel) and unique environment contribution ($R^2_{\text{env}}$; lower panel) for each amino acid, computed separately for each of the 13 jackknife replicates (one point per held-out GTDB class, $n = 13$). The horizontal tick marks the full-data estimate. Amino acids are ordered by global auxotrophy frequency (rarest left). Wide spread for glutamine, asparagine, and serine reflects the phylogenetic concentration of auxotrophs within a small number of lineages.

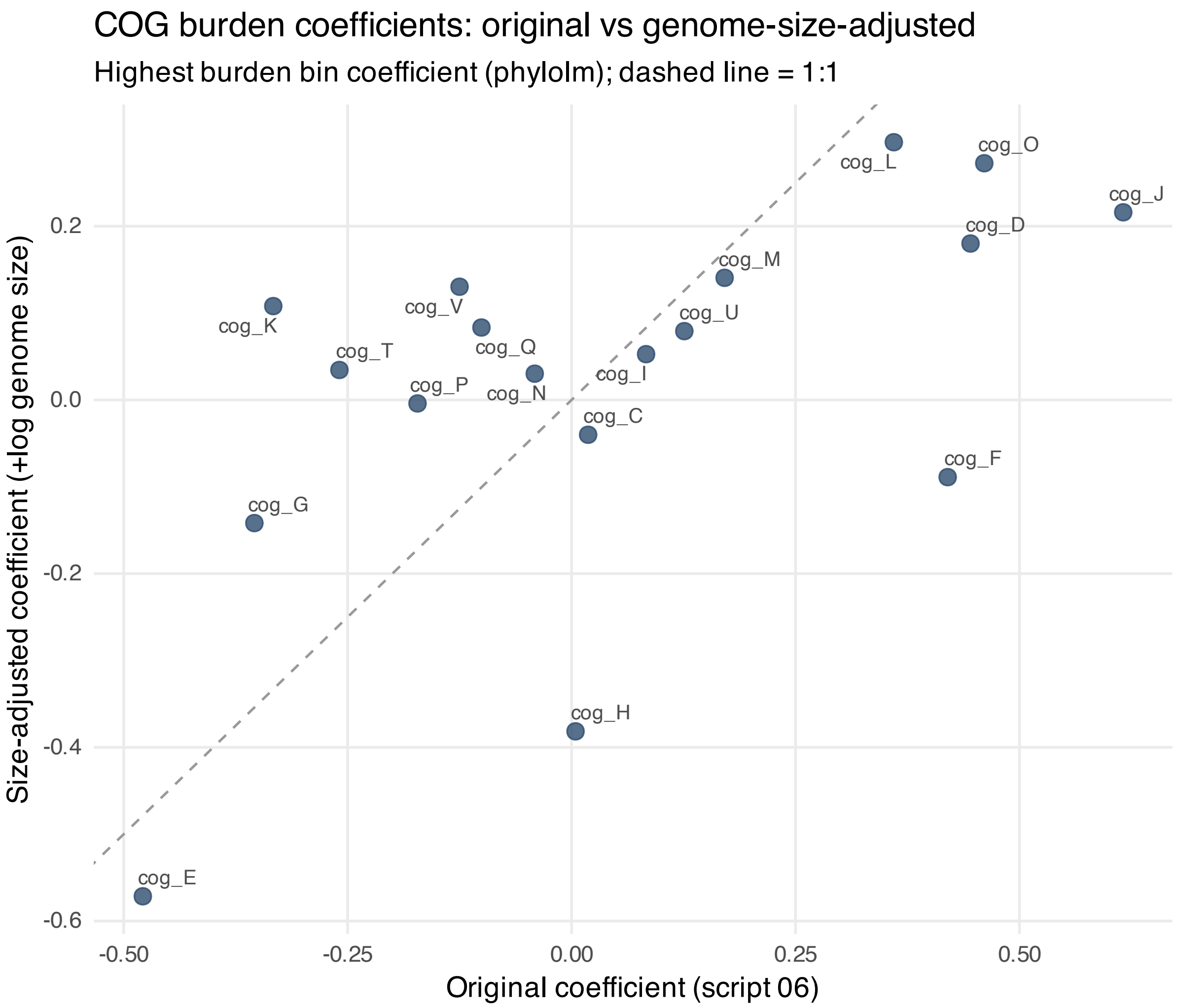


**Figure S17.** COG-burden coefficients: original versus genome-size-adjusted. Scatter of highest-burden-bin phylolm coefficients for each COG category from models without ($x$-axis) and with ($y$-axis) log(Genome_Size) as an additional fixed-effect covariate. Dashed line indicates 1:1. Pearson $r = 0.596$; mean $|\Delta\text{coefficient}| = 0.17$. See Supplementary Discussion 7.

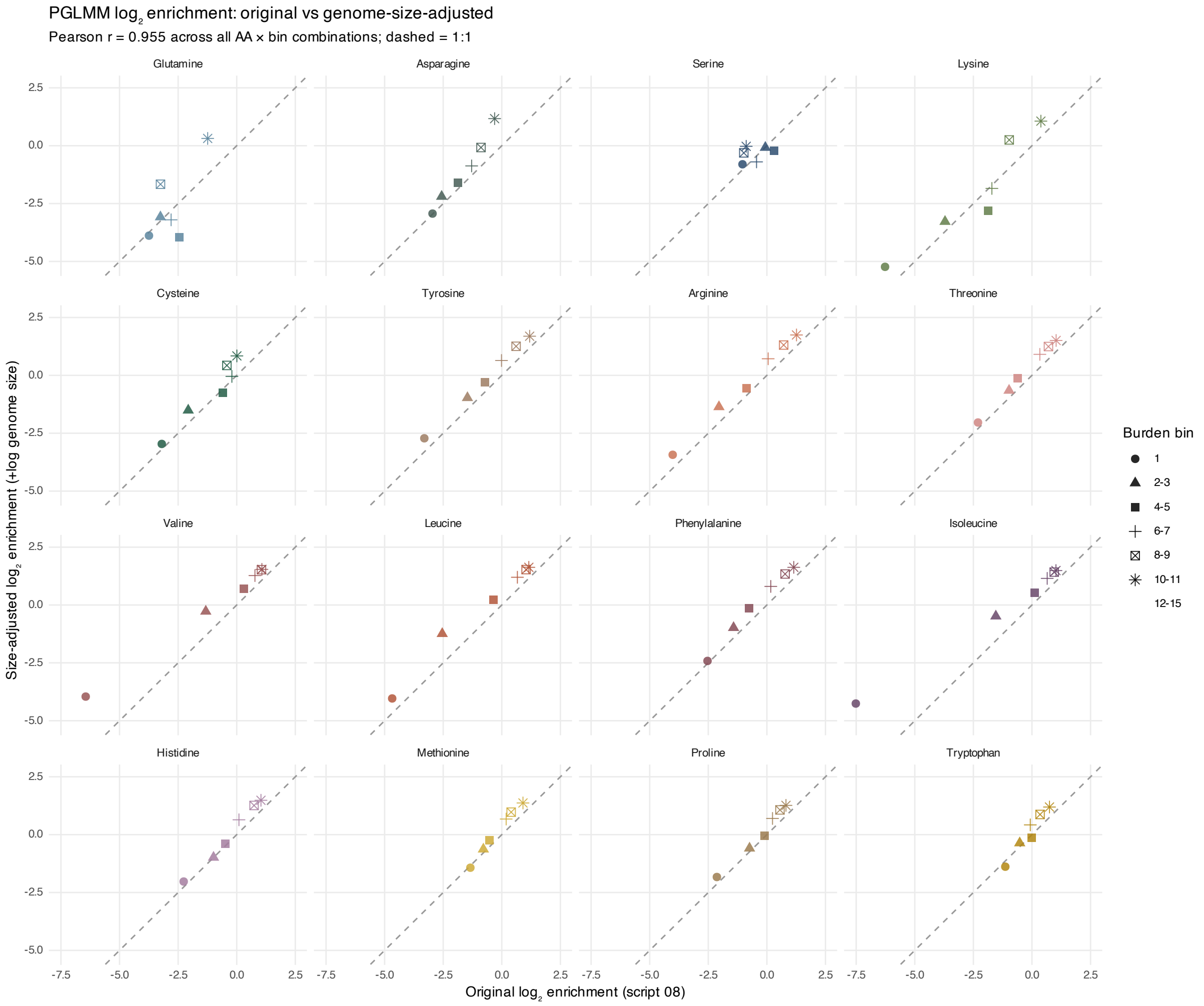


**Figure S18.** Auxotrophy identity $\log_2$ enrichment profiles: original versus genome-size-adjusted. Scatter of per-amino-acid × per-burden-bin $\log_2$ enrichment values from models without ($x$-axis) and with ($y$-axis) log(Genome_Size) as a fixed-effect covariate. Pearson $r = 0.955$; mean $|\Delta \log_2 \text{ enrichment}| = 0.54$. The priority ordering and crossover structure are preserved across amino acids. See Supplementary Discussion 7.

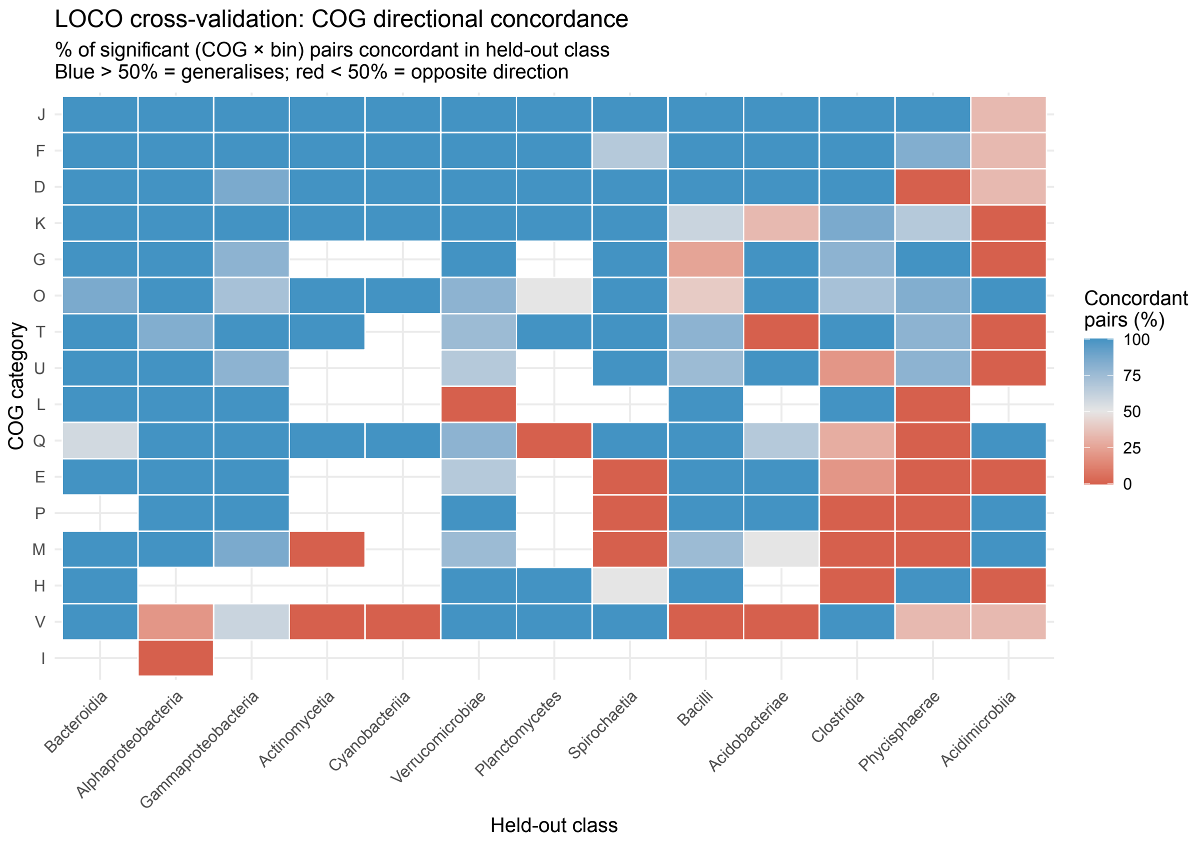


**Figure S19.** LOCO cross-validation: COG directional concordance per category and class. Heatmap showing the percentage of significant (COG category × burden bin) pairs in which the direction of the COG fraction shift in the held-out class matches the sign of the training phylolm coefficient. Each cell shows one COG category (rows) × held-out GTDB class (columns) combination. Blue (>50%) indicates that the training-set direction generalises to the held-out class; red (<50%) indicates reversal. Overall concordance was 76.7% across all held-out classes. See Supplementary Methods S13.

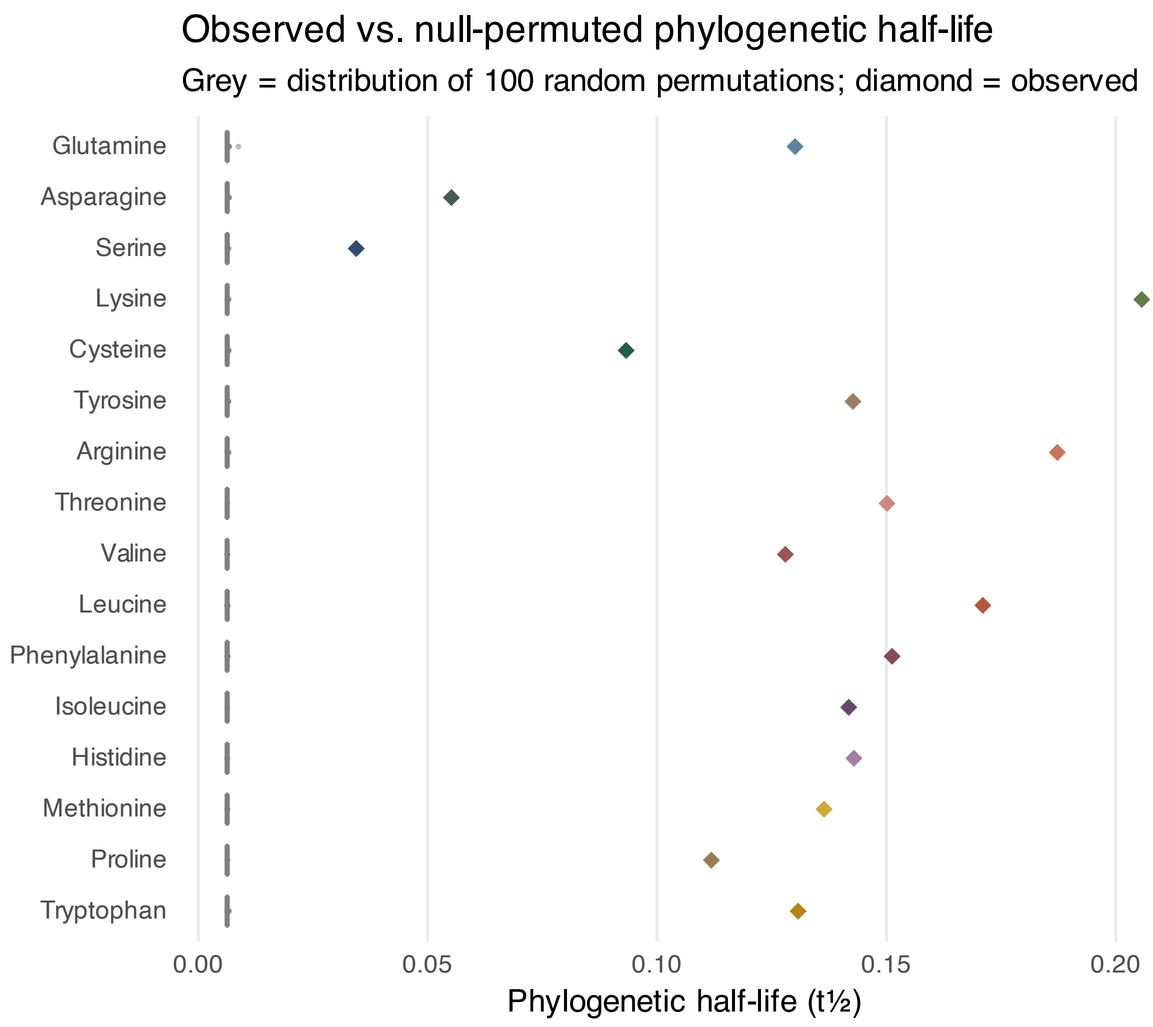


**Figure S20.** Null permutation validation of phylogenetic half-life estimates. Grey boxplots show the distribution of half-lives ($t_{1/2} = \ln(2)/\hat{\alpha}$) from $N = 100$ random permutations of auxotrophy assignments across tree tips, per amino acid. Coloured diamonds show the observed half-life. Under random permutation, $\hat{\alpha}$ consistently converged to its upper bound, yielding half-lives near zero for all amino acids and all permutations. Observed half-lives are substantially larger than any permuted value, confirming that the reported phylogenetic signal reflects genuine evolutionary structure rather than a statistical artefact of tree topology or taxon sampling. See Supplementary Methods S2.

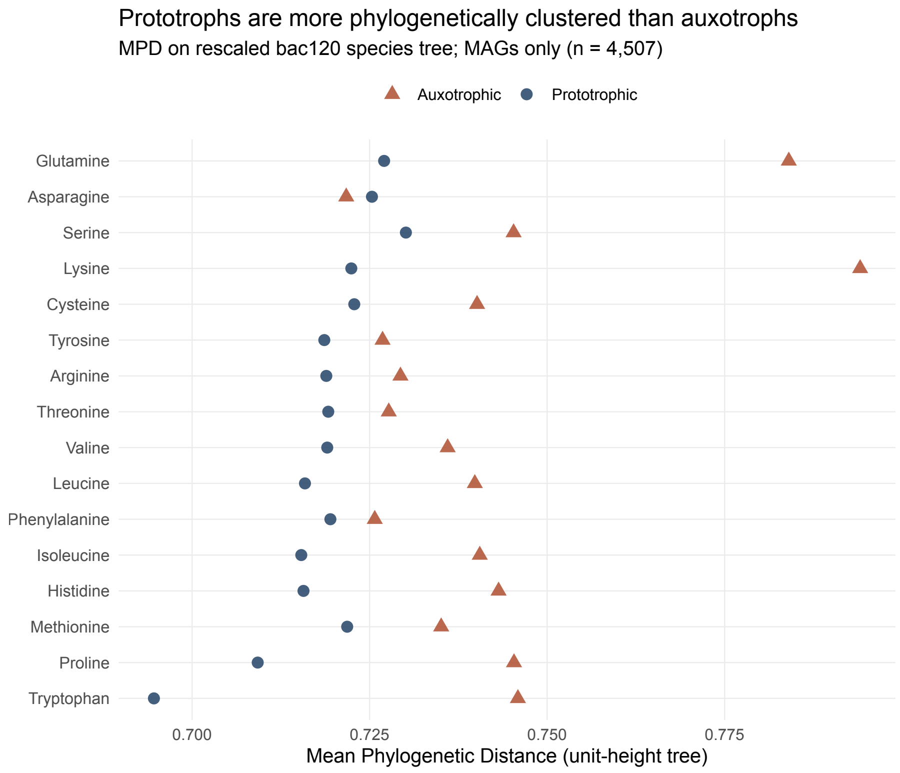


**Figure S21.** Prototrophs are more phylogenetically clustered than auxotrophs. Mean Phylogenetic Distance (MPD) computed from all pairwise cophenetic distances on the rescaled bac120 species tree, separately for prototrophic (blue circles) and auxotrophic (red triangles) genomes for each amino acid. Amino acids are ordered by global auxotrophy frequency (rarest at top). A lower MPD indicates that members of the group are, on average, more closely related. Prototrophs consistently show lower MPD than auxotrophs across nearly all amino acids, indicating that prototrophic lineages are phylogenetically clustered rather than scattered across the tree. This clustering explains why standard GLMs underestimate auxotrophy prevalence: prototrophic observations are not independent, and counting closely related prototrophic lineages as separate data points inflates the apparent prevalence of prototrophy. MAGs only; $n = 4{,}507$ genomes.

**Table S1a.** Phyloglm environment model: intercepts (log-odds scale).

| Amino acid | Predictor | Estimate (log-odds) | Std. error | Z value | P value | P adj (BH) | Sig |
|---|---|---|---|---|---|---|---|
| Lysine | (Intercept) | 1.944 | 0.471 | 4.127 | 3.680e-5 | 6.140e-5 | *** |
| Isoleucine | (Intercept) | 0.365 | 0.192 | 1.899 | 0.0575 | 0.0719 | |
| Arginine | (Intercept) | 0.223 | 0.279 | 0.799 | 0.425 | 0.708 | |
| Serine | (Intercept) | 2.748 | 0.191 | 14.426 | 3.540e-47 | 1.770e-46 | *** |
| Methionine | (Intercept) | -0.244 | 0.238 | -1.026 | 0.305 | 0.305 | |
| Tryptophan | (Intercept) | -0.173 | 0.199 | -0.872 | 0.383 | 0.479 | |
| Phenylalanine | (Intercept) | 0.112 | 0.251 | 0.445 | 0.656 | 0.656 | |
| Tyrosine | (Intercept) | 0.472 | 0.256 | 1.843 | 0.0653 | 0.109 | |
| Cysteine | (Intercept) | 1.865 | 0.205 | 9.108 | 8.410e-20 | 4.210e-19 | *** |
| Leucine | (Intercept) | 0.033 | 0.27 | 0.122 | 0.903 | 0.903 | |
| Histidine | (Intercept) | -0.175 | 0.261 | -0.67 | 0.503 | 0.503 | |
| Proline | (Intercept) | 0.229 | 0.215 | 1.068 | 0.285 | 0.476 | |
| Asparagine | (Intercept) | 2.427 | 0.283 | 8.577 | 9.760e-18 | 4.880e-17 | *** |
| Valine | (Intercept) | 0.751 | 0.185 | 4.056 | 4.990e-5 | 2.490e-4 | *** |
| Threonine | (Intercept) | -0.236 | 0.248 | -0.95 | 0.342 | 0.342 | |
| Glutamine | (Intercept) | 2.077 | 0.693 | 2.995 | 0.0027 | 0.0027 | ** |

Significance: *** $p < 0.001$, ** $p < 0.01$, * $p < 0.05$, · $p < 0.1$. P adj = BH-corrected $p$-value.

**Table S1b.** Phyloglm environment model: environment coefficients (log-odds scale).

| Amino acid | Predictor | Estimate (log-odds) | Std. error | Z value | P value | P adj (BH) | Sig |
|---|---|---|---|---|---|---|---|
| Lysine | Animal | -1.132 | 0.233 | -4.855 | 1.210e-6 | 3.020e-6 | *** |
| Lysine | Aquatic | -1.153 | 0.229 | -5.045 | 4.540e-7 | 2.270e-6 | *** |
| Lysine | Soil | 0.075 | 0.251 | 0.298 | 0.766 | 0.766 | |
| Lysine | Plant | 2.21 | 0.659 | 3.355 | 7.940e-4 | 9.920e-4 | *** |
| Isoleucine | Animal | -0.204 | 0.093 | -2.189 | 0.0286 | 0.0477 | * |
| Isoleucine | Aquatic | -0.013 | 0.071 | -0.182 | 0.856 | 0.856 | |
| Isoleucine | Soil | -0.251 | 0.095 | -2.653 | 0.008 | 0.0199 | * |
| Isoleucine | Plant | 0.469 | 0.157 | 2.984 | 0.0029 | 0.0142 | * |
| Arginine | Animal | -0.184 | 0.081 | -2.277 | 0.0228 | 0.0834 | |
| Arginine | Aquatic | -0.011 | 0.056 | -0.203 | 0.839 | 0.84 | |
| Arginine | Soil | 0.174 | 0.082 | 2.128 | 0.0334 | 0.0834 | |
| Arginine | Plant | 0.022 | 0.109 | 0.201 | 0.84 | 0.84 | |
| Serine | Animal | -0.926 | 0.2 | -4.64 | 3.490e-6 | 8.720e-6 | *** |
| Serine | Aquatic | -0.25 | 0.193 | -1.297 | 0.194 | 0.194 | |
| Serine | Soil | 0.41 | 0.273 | 1.498 | 0.134 | 0.168 | |
| Serine | Plant | 0.767 | 0.495 | 1.549 | 0.121 | 0.168 | |
| Methionine | Animal | -0.487 | 0.096 | -5.062 | 4.150e-7 | 1.040e-6 | *** |
| Methionine | Aquatic | -0.45 | 0.088 | -5.095 | 3.490e-7 | 1.040e-6 | *** |
| Methionine | Soil | 0.348 | 0.102 | 3.422 | 6.220e-4 | 7.770e-4 | *** |
| Methionine | Plant | 0.588 | 0.162 | 3.639 | 2.730e-4 | 4.550e-4 | *** |
| Tryptophan | Animal | -0.602 | 0.097 | -6.195 | 5.810e-10 | 2.910e-9 | *** |
| Tryptophan | Aquatic | -0.006 | 0.072 | -0.089 | 0.929 | 0.929 | |
| Tryptophan | Soil | 0.179 | 0.097 | 1.842 | 0.0655 | 0.109 | |
| Tryptophan | Plant | 0.43 | 0.149 | 2.879 | 0.004 | 0.01 | ** |
| Phenylalanine | Animal | -0.388 | 0.089 | -4.381 | 1.180e-5 | 5.900e-5 | *** |
| Phenylalanine | Aquatic | -0.202 | 0.074 | -2.731 | 0.0063 | 0.0105 | * |
| Phenylalanine | Soil | -0.062 | 0.091 | -0.682 | 0.495 | 0.619 | |
| Phenylalanine | Plant | 0.652 | 0.165 | 3.951 | 7.790e-5 | 1.950e-4 | *** |
| Tyrosine | Animal | -0.336 | 0.092 | -3.66 | 2.520e-4 | 0.0013 | ** |
| Tyrosine | Aquatic | 0.027 | 0.065 | 0.422 | 0.673 | 0.673 | |
| Tyrosine | Soil | 0.168 | 0.091 | 1.842 | 0.0655 | 0.109 | |
| Tyrosine | Plant | 0.14 | 0.132 | 1.064 | 0.287 | 0.359 | |
| Cysteine | Animal | -0.493 | 0.14 | -3.512 | 4.440e-4 | 0.0011 | ** |
| Cysteine | Aquatic | -0.167 | 0.111 | -1.507 | 0.132 | 0.165 | |
| Cysteine | Soil | 0.393 | 0.168 | 2.34 | 0.0193 | 0.0322 | * |
| Cysteine | Plant | 0.267 | 0.245 | 1.087 | 0.277 | 0.277 | |
| Leucine | Animal | -0.338 | 0.085 | -3.967 | 7.270e-5 | 3.630e-4 | *** |
| Leucine | Aquatic | -0.082 | 0.066 | -1.228 | 0.219 | 0.274 | |
| Leucine | Soil | 0.137 | 0.087 | 1.584 | 0.113 | 0.189 | |
| Leucine | Plant | 0.282 | 0.135 | 2.092 | 0.0365 | 0.0912 | |
| Histidine | Animal | -0.486 | 0.098 | -4.962 | 6.980e-7 | 1.750e-6 | *** |
| Histidine | Aquatic | -0.501 | 0.09 | -5.561 | 2.670e-8 | 1.340e-7 | *** |
| Histidine | Soil | 0.393 | 0.105 | 3.757 | 1.720e-4 | 2.860e-4 | *** |
| Histidine | Plant | 0.595 | 0.164 | 3.627 | 2.870e-4 | 3.590e-4 | *** |
| Proline | Animal | 0.018 | 0.09 | 0.205 | 0.838 | 0.838 | |
| Proline | Aquatic | -0.309 | 0.064 | -4.8 | 1.590e-6 | 7.930e-6 | *** |
| Proline | Soil | 0.218 | 0.093 | 2.334 | 0.0196 | 0.0489 | * |
| Proline | Plant | 0.073 | 0.131 | 0.554 | 0.58 | 0.725 | |

*Continued on next page*

*. . . continued*

| **Amino acid** | **Predictor** | **Estimate (log-odds)** | **Std. error** | **Z value** | **P value** | **P adj (BH)** | **Sig** |
|---|---|---|---|---|---|---|---|
| Asparagine | Animal | -0.339 | 0.175 | -1.939 | 0.0525 | 0.0656 | |
| Asparagine | Aquatic | -0.734 | 0.141 | -5.198 | 2.010e-7 | 5.030e-7 | *** |
| Asparagine | Soil | 0.846 | 0.251 | 3.374 | 7.400e-4 | 0.0012 | ** |
| Asparagine | Plant | 0.228 | 0.313 | 0.728 | 0.467 | 0.467 | |
| Valine | Animal | -0.326 | 0.102 | -3.182 | 0.0015 | 0.0037 | ** |
| Valine | Aquatic | -0.071 | 0.078 | -0.911 | 0.362 | 0.362 | |
| Valine | Soil | -0.125 | 0.103 | -1.212 | 0.225 | 0.282 | |
| Valine | Plant | 0.521 | 0.179 | 2.907 | 0.0037 | 0.0061 | ** |
| Threonine | Animal | -0.29 | 0.088 | -3.278 | 0.001 | 0.0026 | ** |
| Threonine | Aquatic | -0.075 | 0.068 | -1.104 | 0.27 | 0.337 | |
| Threonine | Soil | -0.218 | 0.092 | -2.373 | 0.0177 | 0.0294 | * |
| Threonine | Plant | 0.583 | 0.154 | 3.782 | 1.550e-4 | 7.760e-4 | *** |
| Glutamine | Animal | -1.259 | 0.199 | -6.31 | 2.790e-10 | 3.720e-10 | *** |
| Glutamine | Aquatic | -1.304 | 0.193 | -6.757 | 1.410e-11 | 5.640e-11 | *** |
| Glutamine | Soil | 2.563 | 0.386 | 6.646 | 3.000e-11 | 6.010e-11 | *** |

Significance: *** $p < 0.001$, ** $p < 0.01$, * $p < 0.05$, · $p < 0.1$. P adj = BH-corrected $p$-value.

**Table S2a.** $R^2_{\text{lik}}$ variance partitioning. $R^2_{\text{total}}$: total variance explained; $R^2_{\text{phylo}}$: phylogenetic baseline; $R^2_{\text{env}}$: unique environment contribution. Standard errors from jackknife ($n = 13$ classes). Amino acids ordered by auxotrophy frequency (rarest first).

| Amino acid | R2_total (SE) | R2_phylo | R2_env unique (SE) | R2_env % of total |
|---|---|---|---|---|
| L-Glutamine | 0.224 (0.142) | 0.137 | 0.087 (0.107) | 38.9 |
| L-Asparagine | 0.252 (0.199) | 0.218 | 0.034 (0.022) | 13.5 |
| L-Serine | 0.054 (0.013) | 0.026 | 0.027 (0.009) | 50.4 |
| L-Lysine | 0.329 (0.086) | 0.289 | 0.04 (0.027) | 12.1 |
| L-Cysteine | 0.291 (0.185) | 0.274 | 0.017 (0.01) | 5.8 |
| L-Tyrosine | 0.412 (0.114) | 0.385 | 0.027 (0.013) | 6.6 |
| L-Arginine | 0.479 (0.047) | 0.465 | 0.014 (0.013) | 2.9 |
| L-Threonine | 0.44 (0.105) | 0.421 | 0.019 (0.023) | 4.3 |
| L-Valine | 0.411 (0.058) | 0.403 | 0.008 (0.015) | 1.9 |
| L-Leucine | 0.463 (0.114) | 0.438 | 0.025 (0.018) | 5.4 |
| L-Phenylalanine | 0.42 (0.092) | 0.402 | 0.018 (0.044) | 4.3 |
| L-Isoleucine | 0.424 (0.063) | 0.417 | 0.007 (0.029) | 1.7 |
| L-Histidine | 0.471 (0.064) | 0.443 | 0.028 (0.014) | 5.9 |
| L-Methionine | 0.434 (0.056) | 0.409 | 0.025 (0.021) | 5.8 |
| L-Proline | 0.404 (0.057) | 0.373 | 0.031 (0.026) | 7.7 |
| L-Tryptophan | 0.455 (0.074) | 0.427 | 0.027 (0.014) | 5.9 |

**Table S2b.** ΔAIC predictor importance: COG composition and broad environment class.

| Amino acid | Predictor block | ΔAIC | LRT P value |
|---|---|---|---|
| Lysine | COG composition | 128.25 | 7.330e-26 |
| Lysine | Broad environment class | 4.13 | 0.0164 |
| Isoleucine | COG composition | 512.61 | 1.240e-105 |
| Isoleucine | Broad environment class | -11.57 | 1.0 |
| Arginine | COG composition | 431.87 | 1.380e-88 |
| Arginine | Broad environment class | -6.86 | 0.888 |
| Serine | COG composition | 64.32 | 1.690e-13 |
| Serine | Broad environment class | -0.48 | 0.111 |
| Methionine | COG composition | 328.85 | 5.640e-67 |
| Methionine | Broad environment class | 14.1 | 1.920e-4 |
| Tryptophan | COG composition | 435.94 | 1.910e-89 |
| Tryptophan | Broad environment class | 69.78 | 5.130e-16 |
| Phenylalanine | COG composition | 316.36 | 2.280e-64 |
| Phenylalanine | Broad environment class | -4.53 | 0.482 |
| Tyrosine | COG composition | 295.98 | 3.980e-60 |
| Tyrosine | Broad environment class | -6.02 | 0.74 |
| Cysteine | COG composition | 186.34 | 1.520e-37 |
| Cysteine | Broad environment class | 4.57 | 0.0136 |
| Leucine | COG composition | 363.22 | 3.650e-74 |
| Leucine | Broad environment class | -18.54 | 1.0 |
| Histidine | COG composition | 324.68 | 4.180e-66 |
| Histidine | Broad environment class | -9.37 | 1.0 |
| Proline | COG composition | 404.34 | 8.550e-83 |
| Proline | Broad environment class | 15.89 | 8.390e-5 |
| Asparagine | COG composition | 94.31 | 3.330e-19 |
| Asparagine | Broad environment class | -46.67 | 1.0 |
| Valine | COG composition | 279.19 | 1.220e-56 |
| Valine | Broad environment class | 1.45 | 0.0509 |
| Threonine | COG composition | 241.96 | 6.130e-49 |
| Threonine | Broad environment class | 15.19 | 1.160e-4 |
| Glutamine | COG composition | 101.77 | 1.190e-20 |
| Glutamine | Broad environment class | 22.94 | 3.140e-6 |

**Table S3.** Univariate COG model coefficients (phyloglm, log-odds per 1 SD of CLR-transformed COG fraction). Translation (COG J) and Amino Acid Transport (COG E) excluded from multivariate models. All 16 Translation models reached the btol boundary.

| Amino acid | COG category | COG class | Log-odds (per 1 SD) | Std. error | Z value | P value | P adj (BH) | Sig | Convergence warning |
|---|---|---|---|---|---|---|---|---|---|
| Lysine | Energy Production | Metabolism | -0.107 | 0.036 | 2.949 | 0.0032 | 0.0039 | ** | |
| Lysine | Cell Cycle | Cellular Processes | 0.109 | 0.038 | -2.887 | 0.0039 | 0.0047 | ** | |
| Lysine | Amino Acid Transport | Metabolism | -0.125 | 0.037 | 3.346 | 8.190e-4 | 0.0011 | ** | |
| Lysine | Nucleotide Transport | Metabolism | 0.244 | 0.047 | -5.244 | 1.570e-7 | 4.280e-7 | *** | |
| Lysine | Carbohydrate Transport | Metabolism | 0.069 | 0.046 | -1.522 | 0.128 | 0.131 | | |
| Lysine | Coenzyme Transport | Metabolism | -0.108 | 0.032 | 3.402 | 6.690e-4 | 9.400e-4 | *** | |
| Lysine | Lipid Transport | Metabolism | -0.089 | 0.035 | 2.503 | 0.0123 | 0.0139 | * | |
| Lysine | Translation | Information Storage | 0.459 | 0.059 | -7.742 | 9.790e-15 | 7.340e-14 | *** | btol limit reached |
| Lysine | Transcription | Information Storage | 0.064 | 0.036 | -1.789 | 0.0736 | 0.0774 | | |
| Lysine | Replication/Repair | Information Storage | 0.196 | 0.038 | -5.21 | 1.890e-7 | 5.080e-7 | *** | |
| Lysine | Cell Wall | Cellular Processes | 0.138 | 0.037 | -3.677 | 2.360e-4 | 3.770e-4 | *** | |
| Lysine | Motility | Cellular Processes | -0.108 | 0.039 | 2.771 | 0.0056 | 0.0066 | ** | |
| Lysine | Posttrans. Mod | Cellular Processes | 0.212 | 0.039 | -5.418 | 6.020e-8 | 1.840e-7 | *** | |
| Lysine | Inorganic Ion Transport | Metabolism | 0.037 | 0.03 | -1.24 | 0.215 | 0.216 | | |
| Lysine | Secondary Metabolites | Metabolism | -0.129 | 0.041 | 3.147 | 0.0016 | 0.0021 | ** | |
| Lysine | Signal Transduction | Cellular Processes | -0.168 | 0.04 | 4.224 | 2.400e-5 | 4.640e-5 | *** | |
| Lysine | Intracellular Traffic | Cellular Processes | 0.087 | 0.039 | -2.228 | 0.0259 | 0.0282 | * | |
| Lysine | Defense | Cellular Processes | 0.055 | 0.042 | -1.306 | 0.192 | 0.193 | | |
| Isoleucine | Energy Production | Metabolism | -0.116 | 0.032 | 3.65 | 2.630e-4 | 4.110e-4 | *** | |
| Isoleucine | Cell Cycle | Cellular Processes | 0.196 | 0.033 | -5.978 | 2.260e-9 | 8.790e-9 | *** | |
| Isoleucine | Amino Acid Transport | Metabolism | -0.327 | 0.038 | 8.636 | 5.840e-18 | 8.010e-17 | *** | |
| Isoleucine | Nucleotide Transport | Metabolism | 0.229 | 0.035 | -6.563 | 5.260e-11 | 2.440e-10 | *** | |
| Isoleucine | Carbohydrate Transport | Metabolism | -0.097 | 0.035 | 2.735 | 0.0062 | 0.0074 | ** | |
| Isoleucine | Coenzyme Transport | Metabolism | -0.124 | 0.032 | 3.911 | 9.200e-5 | 1.610e-4 | *** | |
| Isoleucine | Lipid Transport | Metabolism | -0.072 | 0.033 | 2.158 | 0.031 | 0.0336 | * | |
| Isoleucine | Translation | Information Storage | 0.388 | 0.038 | -10.093 | 5.900e-24 | 1.700e-22 | *** | btol limit reached |
| Isoleucine | Transcription | Information Storage | -0.131 | 0.035 | 3.796 | 1.470e-4 | 2.460e-4 | *** | |
| Isoleucine | Replication/Repair | Information Storage | 0.142 | 0.029 | -4.862 | 1.160e-6 | 2.790e-6 | *** | |

*Continued on next page*

*. . . continued*

| Amino acid | COG category | COG class | Log-odds (per 1 SD) | Std. error | Z value | P value | P adj (BH) | Sig | Convergence warning |
|---|---|---|---|---|---|---|---|---|---|
| Isoleucine | Cell Wall | Cellular Processes | 0.296 | 0.042 | -7.126 | 1.040e-12 | 6.090e-12 | *** | |
| Isoleucine | Motility | Cellular Processes | 0.067 | 0.031 | -2.15 | 0.0316 | 0.0342 | * | |
| Isoleucine | Posttrans. Mod | Cellular Processes | 0.305 | 0.038 | -8.022 | 1.040e-15 | 9.060e-15 | *** | |
| Isoleucine | Inorganic Ion Transport | Metabolism | -0.175 | 0.032 | 5.374 | 7.690e-8 | 2.310e-7 | *** | |
| Isoleucine | Secondary Metabolites | Metabolism | -0.14 | 0.037 | 3.819 | 1.340e-4 | 2.250e-4 | *** | |
| Isoleucine | Signal Transduction | Cellular Processes | -0.122 | 0.032 | 3.838 | 1.240e-4 | 2.110e-4 | *** | |
| Isoleucine | Intracellular Traffic | Cellular Processes | 0.19 | 0.037 | -5.128 | 2.930e-7 | 7.810e-7 | *** | |
| Isoleucine | Defense | Cellular Processes | 0.082 | 0.034 | -2.431 | 0.0151 | 0.0167 | * | |
| Arginine | Energy Production | Metabolism | -0.107 | 0.029 | 3.662 | 2.500e-4 | 3.950e-4 | *** | |
| Arginine | Cell Cycle | Cellular Processes | 0.123 | 0.031 | -3.945 | 7.970e-5 | 1.420e-4 | *** | |
| Arginine | Amino Acid Transport | Metabolism | -0.116 | 0.033 | 3.533 | 4.110e-4 | 6.020e-4 | *** | |
| Arginine | Nucleotide Transport | Metabolism | 0.239 | 0.034 | -6.938 | 3.970e-12 | 2.200e-11 | *** | |
| Arginine | Carbohydrate Transport | Metabolism | -0.045 | 0.033 | 1.389 | 0.165 | 0.168 | | |
| Arginine | Coenzyme Transport | Metabolism | -0.074 | 0.028 | 2.65 | 0.0081 | 0.0094 | ** | |
| Arginine | Lipid Transport | Metabolism | -0.059 | 0.031 | 1.898 | 0.0576 | 0.0613 | | |
| Arginine | Translation | Information Storage | 0.437 | 0.042 | -10.377 | 3.170e-25 | 1.140e-23 | *** | btol limit reached |
| Arginine | Transcription | Information Storage | -0.072 | 0.031 | 2.33 | 0.0198 | 0.0217 | * | |
| Arginine | Replication/Repair | Information Storage | 0.189 | 0.029 | -6.555 | 5.550e-11 | 2.540e-10 | *** | |
| Arginine | Cell Wall | Cellular Processes | 0.261 | 0.038 | -6.812 | 9.600e-12 | 4.940e-11 | *** | |
| Arginine | Motility | Cellular Processes | -0.118 | 0.03 | 3.942 | 8.070e-5 | 1.430e-4 | *** | |
| Arginine | Posttrans. Mod | Cellular Processes | 0.284 | 0.036 | -7.773 | 7.680e-15 | 5.980e-14 | *** | |
| Arginine | Inorganic Ion Transport | Metabolism | -0.099 | 0.029 | 3.443 | 5.760e-4 | 8.210e-4 | *** | |
| Arginine | Secondary Metabolites | Metabolism | -0.14 | 0.035 | 4.001 | 6.320e-5 | 1.170e-4 | *** | |
| Arginine | Signal Transduction | Cellular Processes | -0.164 | 0.031 | 5.309 | 1.100e-7 | 3.180e-7 | *** | |
| Arginine | Intracellular Traffic | Cellular Processes | 0.088 | 0.034 | -2.615 | 0.0089 | 0.0103 | * | |
| Arginine | Defense | Cellular Processes | 0.096 | 0.031 | -3.092 | 0.002 | 0.0025 | ** | |
| Serine | Energy Production | Metabolism | -0.262 | 0.052 | 4.996 | 5.870e-7 | 1.480e-6 | *** | |
| Serine | Cell Cycle | Cellular Processes | 0.386 | 0.052 | -7.414 | 1.220e-13 | 8.000e-13 | *** | |
| Serine | Amino Acid Transport | Metabolism | -0.215 | 0.054 | 3.995 | 6.460e-5 | 1.180e-4 | *** | |
| Serine | Nucleotide Transport | Metabolism | 0.418 | 0.051 | -8.12 | 4.650e-16 | 4.790e-15 | *** | |

*Continued on next page*

*. . . continued*

| Amino acid | COG category | COG class | Log-odds (per 1 SD) | Std. error | Z value | P value | P adj (BH) | Sig | Convergence warning |
|---|---|---|---|---|---|---|---|---|---|
| Serine | Carbohydrate Transport | Metabolism | 0.15 | 0.06 | -2.505 | 0.0122 | 0.0138 | * | |
| Serine | Coenzyme Transport | Metabolism | 0.285 | 0.058 | -4.949 | 7.480e-7 | 1.860e-6 | *** | |
| Serine | Lipid Transport | Metabolism | -0.152 | 0.044 | 3.481 | 4.990e-4 | 7.190e-4 | *** | |
| Serine | Translation | Information Storage | 0.368 | 0.043 | -8.542 | 1.320e-17 | 1.730e-16 | *** | btol limit reached |
| Serine | Transcription | Information Storage | 0.144 | 0.056 | -2.577 | 0.01 | 0.0114 | * | |
| Serine | Replication/Repair | Information Storage | 0.261 | 0.043 | -6.089 | 1.140e-9 | 4.610e-9 | *** | |
| Serine | Cell Wall | Cellular Processes | 0.337 | 0.062 | -5.423 | 5.870e-8 | 1.820e-7 | *** | |
| Serine | Motility | Cellular Processes | -0.162 | 0.046 | 3.516 | 4.380e-4 | 6.370e-4 | *** | |
| Serine | Posttrans. Mod | Cellular Processes | 0.165 | 0.048 | -3.43 | 6.030e-4 | 8.550e-4 | *** | |
| Serine | Inorganic Ion Transport | Metabolism | -0.157 | 0.042 | 3.739 | 1.850e-4 | 3.050e-4 | *** | |
| Serine | Secondary Metabolites | Metabolism | -0.399 | 0.058 | 6.909 | 4.890e-12 | 2.610e-11 | *** | |
| Serine | Signal Transduction | Cellular Processes | -0.25 | 0.053 | 4.684 | 2.810e-6 | 6.230e-6 | *** | |
| Serine | Intracellular Traffic | Cellular Processes | -0.136 | 0.058 | 2.335 | 0.0195 | 0.0216 | * | |
| Serine | Defense | Cellular Processes | 0.241 | 0.068 | -3.546 | 3.910e-4 | 5.780e-4 | *** | |
| Methionine | Energy Production | Metabolism | 0.054 | 0.031 | -1.756 | 0.0791 | 0.0825 | | |
| Methionine | Cell Cycle | Cellular Processes | 0.168 | 0.032 | -5.247 | 1.540e-7 | 4.280e-7 | *** | |
| Methionine | Amino Acid Transport | Metabolism | -0.242 | 0.035 | 6.932 | 4.160e-12 | 2.260e-11 | *** | |
| Methionine | Nucleotide Transport | Metabolism | 0.226 | 0.034 | -6.708 | 1.980e-11 | 9.820e-11 | *** | |
| Methionine | Carbohydrate Transport | Metabolism | -0.105 | 0.036 | 2.922 | 0.0035 | 0.0043 | ** | |
| Methionine | Coenzyme Transport | Metabolism | 0.07 | 0.029 | -2.416 | 0.0157 | 0.0174 | * | |
| Methionine | Lipid Transport | Metabolism | -0.104 | 0.036 | 2.894 | 0.0038 | 0.0046 | ** | |
| Methionine | Translation | Information Storage | 0.344 | 0.037 | -9.3 | 1.410e-20 | 2.410e-19 | *** | btol limit reached |
| Methionine | Transcription | Information Storage | -0.169 | 0.034 | 4.898 | 9.700e-7 | 2.390e-6 | *** | |
| Methionine | Replication/Repair | Information Storage | 0.187 | 0.029 | -6.455 | 1.080e-10 | 4.720e-10 | *** | |
| Methionine | Cell Wall | Cellular Processes | 0.223 | 0.04 | -5.561 | 2.690e-8 | 9.000e-8 | *** | |
| Methionine | Motility | Cellular Processes | 0.062 | 0.031 | -2.003 | 0.0452 | 0.0486 | * | |
| Methionine | Posttrans. Mod | Cellular Processes | 0.307 | 0.038 | -8.174 | 2.990e-16 | 3.400e-15 | *** | |
| Methionine | Inorganic Ion Transport | Metabolism | -0.125 | 0.032 | 3.946 | 7.940e-5 | 1.420e-4 | *** | |
| Methionine | Secondary Metabolites | Metabolism | -0.156 | 0.037 | 4.265 | 2.000e-5 | 3.940e-5 | *** | |
| Methionine | Signal Transduction | Cellular Processes | -0.171 | 0.031 | 5.447 | 5.110e-8 | 1.600e-7 | *** | |

*Continued on next page*

*. . . continued*

| Amino acid | COG category | COG class | Log-odds (per 1 SD) | Std. error | Z value | P value | P adj (BH) | Sig | Convergence warning |
|---|---|---|---|---|---|---|---|---|---|
| Methionine | Intracellular Traffic | Cellular Processes | 0.2 | 0.037 | -5.463 | 4.680e-8 | 1.520e-7 | *** | |
| Methionine | Defense | Cellular Processes | -0.057 | 0.034 | 1.678 | 0.0933 | 0.0963 | | |
| Tryptophan | Energy Production | Metabolism | -0.116 | 0.032 | 3.602 | 3.160e-4 | 4.810e-4 | *** | |
| Tryptophan | Cell Cycle | Cellular Processes | 0.355 | 0.034 | -10.387 | 2.840e-25 | 1.140e-23 | *** | |
| Tryptophan | Amino Acid Transport | Metabolism | -0.141 | 0.036 | 3.948 | 7.870e-5 | 1.420e-4 | *** | |
| Tryptophan | Nucleotide Transport | Metabolism | 0.338 | 0.036 | -9.299 | 1.420e-20 | 2.410e-19 | *** | |
| Tryptophan | Carbohydrate Transport | Metabolism | 0.17 | 0.038 | -4.44 | 9.000e-6 | 1.830e-5 | *** | |
| Tryptophan | Coenzyme Transport | Metabolism | -0.202 | 0.034 | 5.935 | 2.940e-9 | 1.130e-8 | *** | |
| Tryptophan | Lipid Transport | Metabolism | -0.105 | 0.038 | 2.773 | 0.0056 | 0.0066 | ** | |
| Tryptophan | Translation | Information Storage | 0.705 | 0.042 | -16.848 | 1.090e-63 | 3.150e-61 | *** | btol limit reached |
| Tryptophan | Transcription | Information Storage | -0.107 | 0.036 | 3.002 | 0.0027 | 0.0033 | ** | |
| Tryptophan | Replication/Repair | Information Storage | 0.227 | 0.031 | -7.424 | 1.140e-13 | 7.640e-13 | *** | |
| Tryptophan | Cell Wall | Cellular Processes | 0.291 | 0.045 | -6.469 | 9.890e-11 | 4.380e-10 | *** | |
| Tryptophan | Motility | Cellular Processes | -0.142 | 0.035 | 4.113 | 3.900e-5 | 7.390e-5 | *** | |
| Tryptophan | Posttrans. Mod | Cellular Processes | 0.208 | 0.038 | -5.501 | 3.770e-8 | 1.250e-7 | *** | |
| Tryptophan | Inorganic Ion Transport | Metabolism | -0.19 | 0.034 | 5.607 | 2.050e-8 | 7.120e-8 | *** | |
| Tryptophan | Secondary Metabolites | Metabolism | -0.312 | 0.041 | 7.694 | 1.420e-14 | 1.020e-13 | *** | |
| Tryptophan | Signal Transduction | Cellular Processes | -0.273 | 0.033 | 8.387 | 4.990e-17 | 6.250e-16 | *** | |
| Tryptophan | Intracellular Traffic | Cellular Processes | 0.098 | 0.037 | -2.665 | 0.0077 | 0.009 | ** | |
| Tryptophan | Defense | Cellular Processes | 0.155 | 0.034 | -4.507 | 6.580e-6 | 1.370e-5 | *** | |
| Phenylalanine | Energy Production | Metabolism | -0.143 | 0.031 | 4.678 | 2.890e-6 | 6.350e-6 | *** | |
| Phenylalanine | Cell Cycle | Cellular Processes | 0.17 | 0.032 | -5.312 | 1.080e-7 | 3.150e-7 | *** | |
| Phenylalanine | Amino Acid Transport | Metabolism | -0.308 | 0.037 | 8.335 | 7.780e-17 | 9.330e-16 | *** | |
| Phenylalanine | Nucleotide Transport | Metabolism | 0.217 | 0.033 | -6.495 | 8.300e-11 | 3.730e-10 | *** | |
| Phenylalanine | Carbohydrate Transport | Metabolism | -0.153 | 0.036 | 4.257 | 2.070e-5 | 4.030e-5 | *** | |
| Phenylalanine | Coenzyme Transport | Metabolism | -0.15 | 0.032 | 4.721 | 2.350e-6 | 5.320e-6 | *** | |
| Phenylalanine | Lipid Transport | Metabolism | -0.081 | 0.033 | 2.436 | 0.0149 | 0.0166 | * | |
| Phenylalanine | Translation | Information Storage | 0.285 | 0.037 | -7.783 | 7.060e-15 | 5.650e-14 | *** | btol limit reached |
| Phenylalanine | Transcription | Information Storage | -0.129 | 0.034 | 3.826 | 1.300e-4 | 2.210e-4 | *** | |
| Phenylalanine | Replication/Repair | Information Storage | 0.19 | 0.029 | -6.62 | 3.590e-11 | 1.700e-10 | *** | |

*Continued on next page*

*... continued*

| Amino acid | COG category | COG class | Log-odds (per 1 SD) | Std. error | Z value | P value | P adj (BH) | Sig | Convergence warning |
|---|---|---|---|---|---|---|---|---|---|
| Phenylalanine | Cell Wall | Cellular Processes | 0.377 | 0.04 | -9.316 | 1.210e-20 | 2.320e-19 | *** | |
| Phenylalanine | Motility | Cellular Processes | -0.122 | 0.031 | 3.913 | 9.130e-5 | 1.600e-4 | *** | |
| Phenylalanine | Posttrans. Mod | Cellular Processes | 0.365 | 0.038 | -9.492 | 2.260e-21 | 4.640e-20 | *** | |
| Phenylalanine | Inorganic Ion Transport | Metabolism | -0.2 | 0.032 | 6.222 | 4.900e-10 | 2.020e-9 | *** | |
| Phenylalanine | Secondary Metabolites | Metabolism | -0.147 | 0.036 | 4.148 | 3.350e-5 | 6.400e-5 | *** | |
| Phenylalanine | Signal Transduction | Cellular Processes | 0.042 | 0.03 | -1.376 | 0.169 | 0.171 | | |
| Phenylalanine | Intracellular Traffic | Cellular Processes | 0.151 | 0.036 | -4.208 | 2.580e-5 | 4.960e-5 | *** | |
| Phenylalanine | Defense | Cellular Processes | 0.097 | 0.033 | -2.95 | 0.0032 | 0.0039 | ** | |
| Tyrosine | Energy Production | Metabolism | -0.129 | 0.032 | 3.987 | 6.700e-5 | 1.220e-4 | *** | |
| Tyrosine | Cell Cycle | Cellular Processes | 0.247 | 0.035 | -7.046 | 1.850e-12 | 1.040e-11 | *** | |
| Tyrosine | Amino Acid Transport | Metabolism | -0.316 | 0.039 | 8.121 | 4.620e-16 | 4.790e-15 | *** | |
| Tyrosine | Nucleotide Transport | Metabolism | 0.277 | 0.036 | -7.74 | 9.940e-15 | 7.340e-14 | *** | |
| Tyrosine | Carbohydrate Transport | Metabolism | -0.128 | 0.038 | 3.411 | 6.480e-4 | 9.150e-4 | *** | |
| Tyrosine | Coenzyme Transport | Metabolism | -0.166 | 0.033 | 5.008 | 5.490e-7 | 1.400e-6 | *** | |
| Tyrosine | Lipid Transport | Metabolism | -0.118 | 0.035 | 3.37 | 7.500e-4 | 0.001 | ** | |
| Tyrosine | Translation | Information Storage | 0.389 | 0.041 | -9.564 | 1.130e-21 | 2.720e-20 | *** | btol limit reached |
| Tyrosine | Transcription | Information Storage | -0.102 | 0.036 | 2.83 | 0.0046 | 0.0056 | ** | |
| Tyrosine | Replication/Repair | Information Storage | 0.287 | 0.033 | -8.716 | 2.880e-18 | 4.150e-17 | *** | |
| Tyrosine | Cell Wall | Cellular Processes | 0.312 | 0.041 | -7.666 | 1.770e-14 | 1.240e-13 | *** | |
| Tyrosine | Motility | Cellular Processes | -0.151 | 0.033 | 4.6 | 4.220e-6 | 9.050e-6 | *** | |
| Tyrosine | Posttrans. Mod | Cellular Processes | 0.324 | 0.04 | -8.073 | 6.860e-16 | 6.590e-15 | *** | |
| Tyrosine | Inorganic Ion Transport | Metabolism | -0.224 | 0.034 | 6.624 | 3.500e-11 | 1.680e-10 | *** | |
| Tyrosine | Secondary Metabolites | Metabolism | -0.217 | 0.039 | 5.595 | 2.210e-8 | 7.570e-8 | *** | |
| Tyrosine | Signal Transduction | Cellular Processes | -0.198 | 0.034 | 5.852 | 4.860e-9 | 1.840e-8 | *** | |
| Tyrosine | Intracellular Traffic | Cellular Processes | 0.136 | 0.038 | -3.602 | 3.150e-4 | 4.810e-4 | *** | |
| Tyrosine | Defense | Cellular Processes | 0.12 | 0.035 | -3.38 | 7.250e-4 | 0.001 | ** | |
| Cysteine | Energy Production | Metabolism | -0.126 | 0.042 | 3.013 | 0.0026 | 0.0032 | ** | |
| Cysteine | Cell Cycle | Cellular Processes | 0.235 | 0.047 | -5.01 | 5.450e-7 | 1.400e-6 | *** | |
| Cysteine | Amino Acid Transport | Metabolism | -0.198 | 0.044 | 4.454 | 8.440e-6 | 1.720e-5 | *** | |
| Cysteine | Nucleotide Transport | Metabolism | 0.446 | 0.055 | -8.056 | 7.880e-16 | 7.330e-15 | *** | |

*Continued on next page*

*... continued*

| Amino acid | COG category | COG class | Log-odds (per 1 SD) | Std. error | Z value | P value | P adj (BH) | Sig | Convergence warning |
|---|---|---|---|---|---|---|---|---|---|
| Cysteine | Carbohydrate Transport | Metabolism | 0.077 | 0.047 | -1.649 | 0.0991 | 0.102 | | |
| Cysteine | Coenzyme Transport | Metabolism | 0.129 | 0.041 | -3.157 | 0.0016 | 0.0021 | ** | |
| Cysteine | Lipid Transport | Metabolism | -0.132 | 0.041 | 3.221 | 0.0013 | 0.0017 | ** | |
| Cysteine | Translation | Information Storage | 0.606 | 0.059 | -10.316 | 5.950e-25 | 1.910e-23 | *** | btol limit reached |
| Cysteine | Transcription | Information Storage | -0.153 | 0.046 | 3.326 | 8.800e-4 | 0.0012 | ** | |
| Cysteine | Replication/Repair | Information Storage | 0.351 | 0.045 | -7.784 | 7.040e-15 | 5.650e-14 | *** | |
| Cysteine | Cell Wall | Cellular Processes | 0.341 | 0.059 | -5.793 | 6.900e-9 | 2.550e-8 | *** | |
| Cysteine | Motility | Cellular Processes | -0.155 | 0.04 | 3.857 | 1.150e-4 | 1.960e-4 | *** | |
| Cysteine | Posttrans. Mod | Cellular Processes | 0.311 | 0.05 | -6.255 | 3.980e-10 | 1.660e-9 | *** | |
| Cysteine | Inorganic Ion Transport | Metabolism | -0.117 | 0.036 | 3.233 | 0.0012 | 0.0016 | ** | |
| Cysteine | Secondary Metabolites | Metabolism | -0.303 | 0.053 | 5.758 | 8.500e-9 | 3.090e-8 | *** | |
| Cysteine | Signal Transduction | Cellular Processes | -0.257 | 0.047 | 5.449 | 5.050e-8 | 1.600e-7 | *** | |
| Cysteine | Intracellular Traffic | Cellular Processes | 0.251 | 0.053 | -4.736 | 2.180e-6 | 4.980e-6 | *** | |
| Cysteine | Defense | Cellular Processes | -0.135 | 0.038 | 3.559 | 3.730e-4 | 5.540e-4 | *** | |
| Leucine | Energy Production | Metabolism | -0.104 | 0.029 | 3.534 | 4.090e-4 | 6.010e-4 | *** | |
| Leucine | Cell Cycle | Cellular Processes | 0.211 | 0.031 | -6.868 | 6.510e-12 | 3.410e-11 | *** | |
| Leucine | Amino Acid Transport | Metabolism | -0.169 | 0.034 | 5.036 | 4.750e-7 | 1.230e-6 | *** | |
| Leucine | Nucleotide Transport | Metabolism | 0.232 | 0.032 | -7.203 | 5.880e-13 | 3.610e-12 | *** | |
| Leucine | Carbohydrate Transport | Metabolism | -0.125 | 0.034 | 3.652 | 2.600e-4 | 4.090e-4 | *** | |
| Leucine | Coenzyme Transport | Metabolism | -0.156 | 0.03 | 5.116 | 3.110e-7 | 8.230e-7 | *** | |
| Leucine | Lipid Transport | Metabolism | -0.084 | 0.033 | 2.543 | 0.011 | 0.0126 | * | |
| Leucine | Translation | Information Storage | 0.401 | 0.038 | -10.622 | 2.340e-26 | 1.350e-24 | *** | btol limit reached |
| Leucine | Transcription | Information Storage | -0.101 | 0.032 | 3.121 | 0.0018 | 0.0023 | ** | |
| Leucine | Replication/Repair | Information Storage | 0.189 | 0.028 | -6.809 | 9.850e-12 | 4.980e-11 | *** | |
| Leucine | Cell Wall | Cellular Processes | 0.212 | 0.039 | -5.475 | 4.380e-8 | 1.430e-7 | *** | |
| Leucine | Motility | Cellular Processes | -0.075 | 0.03 | 2.521 | 0.0117 | 0.0133 | * | |
| Leucine | Posttrans. Mod | Cellular Processes | 0.469 | 0.043 | -10.849 | 2.010e-27 | 1.930e-25 | *** | |
| Leucine | Inorganic Ion Transport | Metabolism | -0.161 | 0.031 | 5.265 | 1.400e-7 | 3.910e-7 | *** | |
| Leucine | Secondary Metabolites | Metabolism | -0.165 | 0.035 | 4.714 | 2.430e-6 | 5.470e-6 | *** | |
| Leucine | Signal Transduction | Cellular Processes | -0.159 | 0.03 | 5.319 | 1.040e-7 | 3.060e-7 | *** | |

*Continued on next page*

*. . . continued*

| Amino acid | COG category | COG class | Log-odds (per 1 SD) | Std. error | Z value | P value | P adj (BH) | Sig | Convergence warning |
|---|---|---|---|---|---|---|---|---|---|
| Leucine | Intracellular Traffic | Cellular Processes | 0.141 | 0.035 | -4.079 | 4.510e-5 | 8.500e-5 | *** | |
| Leucine | Defense | Cellular Processes | 0.098 | 0.031 | -3.127 | 0.0018 | 0.0023 | ** | |
| Histidine | Energy Production | Metabolism | 0.077 | 0.031 | -2.494 | 0.0126 | 0.0141 | * | |
| Histidine | Cell Cycle | Cellular Processes | 0.13 | 0.032 | -4.046 | 5.200e-5 | 9.730e-5 | *** | |
| Histidine | Amino Acid Transport | Metabolism | -0.209 | 0.035 | 6.006 | 1.900e-9 | 7.490e-9 | *** | |
| Histidine | Nucleotide Transport | Metabolism | 0.216 | 0.034 | -6.297 | 3.030e-10 | 1.280e-9 | *** | |
| Histidine | Carbohydrate Transport | Metabolism | -0.104 | 0.036 | 2.889 | 0.0039 | 0.0047 | ** | |
| Histidine | Coenzyme Transport | Metabolism | -0.082 | 0.031 | 2.691 | 0.0071 | 0.0084 | ** | |
| Histidine | Lipid Transport | Metabolism | -0.062 | 0.034 | 1.818 | 0.069 | 0.0728 | | |
| Histidine | Translation | Information Storage | 0.391 | 0.039 | -10.05 | 9.220e-24 | 2.410e-22 | *** | btol limit reached |
| Histidine | Transcription | Information Storage | -0.106 | 0.034 | 3.085 | 0.002 | 0.0026 | ** | |
| Histidine | Replication/Repair | Information Storage | 0.168 | 0.029 | -5.831 | 5.520e-9 | 2.060e-8 | *** | |
| Histidine | Cell Wall | Cellular Processes | 0.22 | 0.04 | -5.46 | 4.750e-8 | 1.520e-7 | *** | |
| Histidine | Motility | Cellular Processes | -0.101 | 0.032 | 3.171 | 0.0015 | 0.002 | ** | |
| Histidine | Posttrans. Mod | Cellular Processes | 0.275 | 0.038 | -7.294 | 3.010e-13 | 1.930e-12 | *** | |
| Histidine | Inorganic Ion Transport | Metabolism | -0.142 | 0.032 | 4.468 | 7.890e-6 | 1.620e-5 | *** | |
| Histidine | Secondary Metabolites | Metabolism | -0.119 | 0.036 | 3.28 | 0.001 | 0.0014 | ** | |
| Histidine | Signal Transduction | Cellular Processes | -0.146 | 0.031 | 4.659 | 3.170e-6 | 6.930e-6 | *** | |
| Histidine | Intracellular Traffic | Cellular Processes | 0.191 | 0.036 | -5.245 | 1.560e-7 | 4.280e-7 | *** | |
| Histidine | Defense | Cellular Processes | -0.062 | 0.034 | 1.863 | 0.0625 | 0.0662 | | |
| Proline | Energy Production | Metabolism | -0.11 | 0.032 | 3.386 | 7.100e-4 | 9.930e-4 | *** | |
| Proline | Cell Cycle | Cellular Processes | 0.181 | 0.034 | -5.272 | 1.350e-7 | 3.800e-7 | *** | |
| Proline | Amino Acid Transport | Metabolism | -0.232 | 0.037 | 6.353 | 2.110e-10 | 9.080e-10 | *** | |
| Proline | Nucleotide Transport | Metabolism | 0.341 | 0.038 | -8.992 | 2.440e-19 | 3.900e-18 | *** | |
| Proline | Carbohydrate Transport | Metabolism | -0.139 | 0.038 | 3.698 | 2.180e-4 | 3.500e-4 | *** | |
| Proline | Coenzyme Transport | Metabolism | -0.141 | 0.033 | 4.279 | 1.880e-5 | 3.730e-5 | *** | |
| Proline | Lipid Transport | Metabolism | -0.073 | 0.035 | 2.12 | 0.034 | 0.0367 | * | |
| Proline | Translation | Information Storage | 0.43 | 0.041 | -10.605 | 2.830e-26 | 1.360e-24 | *** | btol limit reached |
| Proline | Transcription | Information Storage | -0.183 | 0.037 | 4.982 | 6.300e-7 | 1.580e-6 | *** | |
| Proline | Replication/Repair | Information Storage | 0.226 | 0.031 | -7.208 | 5.670e-13 | 3.550e-12 | *** | |

*Continued on next page*

*. . . continued*

| Amino acid | COG category | COG class | Log-odds (per 1 SD) | Std. error | Z value | P value | P adj (BH) | Sig | Convergence warning |
|---|---|---|---|---|---|---|---|---|---|
| Proline | Cell Wall | Cellular Processes | 0.28 | 0.042 | -6.693 | 2.180e-11 | 1.070e-10 | *** | |
| Proline | Motility | Cellular Processes | -0.153 | 0.033 | 4.593 | 4.370e-6 | 9.260e-6 | *** | |
| Proline | Posttrans. Mod | Cellular Processes | 0.444 | 0.041 | -10.795 | 3.630e-27 | 2.620e-25 | *** | |
| Proline | Inorganic Ion Transport | Metabolism | -0.158 | 0.033 | 4.744 | 2.100e-6 | 4.830e-6 | *** | |
| Proline | Secondary Metabolites | Metabolism | -0.186 | 0.039 | 4.772 | 1.820e-6 | 4.310e-6 | *** | |
| Proline | Signal Transduction | Cellular Processes | -0.246 | 0.035 | 7.12 | 1.080e-12 | 6.240e-12 | *** | |
| Proline | Intracellular Traffic | Cellular Processes | 0.181 | 0.038 | -4.747 | 2.070e-6 | 4.810e-6 | *** | |
| Proline | Defense | Cellular Processes | 0.113 | 0.035 | -3.194 | 0.0014 | 0.0019 | ** | |
| Asparagine | Energy Production | Metabolism | 0.148 | 0.056 | -2.638 | 0.0083 | 0.0097 | ** | |
| Asparagine | Cell Cycle | Cellular Processes | 0.279 | 0.059 | -4.76 | 1.930e-6 | 4.520e-6 | *** | |
| Asparagine | Amino Acid Transport | Metabolism | -0.207 | 0.059 | 3.495 | 4.750e-4 | 6.870e-4 | *** | |
| Asparagine | Nucleotide Transport | Metabolism | 0.501 | 0.057 | -8.773 | 1.740e-18 | 2.640e-17 | *** | |
| Asparagine | Carbohydrate Transport | Metabolism | 0.076 | 0.065 | -1.169 | 0.242 | 0.242 | | |
| Asparagine | Coenzyme Transport | Metabolism | 0.183 | 0.058 | -3.182 | 0.0015 | 0.0019 | ** | |
| Asparagine | Lipid Transport | Metabolism | -0.15 | 0.045 | 3.306 | 9.480e-4 | 0.0013 | ** | |
| Asparagine | Translation | Information Storage | 0.482 | 0.051 | -9.546 | 1.350e-21 | 2.990e-20 | *** | btol limit reached |
| Asparagine | Transcription | Information Storage | -0.089 | 0.063 | 1.407 | 0.159 | 0.163 | | |
| Asparagine | Replication/Repair | Information Storage | 0.352 | 0.046 | -7.656 | 1.920e-14 | 1.320e-13 | *** | |
| Asparagine | Cell Wall | Cellular Processes | 0.216 | 0.066 | -3.246 | 0.0012 | 0.0016 | ** | |
| Asparagine | Motility | Cellular Processes | -0.211 | 0.047 | 4.5 | 6.810e-6 | 1.410e-5 | *** | |
| Asparagine | Posttrans. Mod | Cellular Processes | 0.484 | 0.059 | -8.17 | 3.070e-16 | 3.400e-15 | *** | |
| Asparagine | Inorganic Ion Transport | Metabolism | -0.189 | 0.047 | 4.027 | 5.640e-5 | 1.050e-4 | *** | |
| Asparagine | Secondary Metabolites | Metabolism | -0.314 | 0.059 | 5.345 | 9.040e-8 | 2.680e-7 | *** | |
| Asparagine | Signal Transduction | Cellular Processes | -0.292 | 0.058 | 5.038 | 4.710e-7 | 1.230e-6 | *** | |
| Asparagine | Intracellular Traffic | Cellular Processes | 0.24 | 0.062 | -3.884 | 1.030e-4 | 1.790e-4 | *** | |
| Asparagine | Defense | Cellular Processes | -0.214 | 0.04 | 5.304 | 1.130e-7 | 3.240e-7 | *** | |
| Valine | Energy Production | Metabolism | -0.118 | 0.033 | 3.618 | 2.970e-4 | 4.600e-4 | *** | |
| Valine | Cell Cycle | Cellular Processes | -0.054 | 0.032 | 1.683 | 0.0925 | 0.0961 | | |
| Valine | Amino Acid Transport | Metabolism | -0.316 | 0.039 | 8.045 | 8.620e-16 | 7.760e-15 | *** | |
| Valine | Nucleotide Transport | Metabolism | 0.165 | 0.036 | -4.553 | 5.300e-6 | 1.110e-5 | *** | |

*Continued on next page*

*. . . continued*

| Amino acid | COG category | COG class | Log-odds (per 1 SD) | Std. error | Z value | P value | P adj (BH) | Sig | Convergence warning |
|---|---|---:|---:|---:|---:|---:|---:|---|---:|
| Valine | Carbohydrate Transport | Metabolism | 0.061 | 0.036 | -1.681 | 0.0927 | 0.0961 | | |
| Valine | Coenzyme Transport | Metabolism | 0.073 | 0.031 | -2.329 | 0.0198 | 0.0217 | * | |
| Valine | Lipid Transport | Metabolism | 0.064 | 0.036 | -1.762 | 0.0781 | 0.0818 | | |
| Valine | Translation | Information Storage | 0.218 | 0.038 | -5.757 | 8.580e-9 | 3.090e-8 | *** | btol limit reached |
| Valine | Transcription | Information Storage | -0.136 | 0.036 | 3.722 | 1.970e-4 | 3.210e-4 | *** | |
| Valine | Replication/Repair | Information Storage | 0.106 | 0.03 | -3.598 | 3.210e-4 | 4.860e-4 | *** | |
| Valine | Cell Wall | Cellular Processes | 0.228 | 0.042 | -5.387 | 7.170e-8 | 2.170e-7 | *** | |
| Valine | Motility | Cellular Processes | -0.061 | 0.032 | 1.918 | 0.0552 | 0.0589 | | |
| Valine | Posttrans. Mod | Cellular Processes | 0.211 | 0.038 | -5.609 | 2.040e-8 | 7.120e-8 | *** | |
| Valine | Inorganic Ion Transport | Metabolism | -0.113 | 0.032 | 3.573 | 3.540e-4 | 5.320e-4 | *** | |
| Valine | Secondary Metabolites | Metabolism | -0.113 | 0.037 | 3.047 | 0.0023 | 0.0029 | ** | |
| Valine | Signal Transduction | Cellular Processes | -0.099 | 0.033 | 3.032 | 0.0024 | 0.003 | ** | |
| Valine | Intracellular Traffic | Cellular Processes | 0.149 | 0.038 | -3.934 | 8.370e-5 | 1.480e-4 | *** | |
| Valine | Defense | Cellular Processes | 0.119 | 0.036 | -3.279 | 0.001 | 0.0014 | ** | |
| Threonine | Energy Production | Metabolism | -0.097 | 0.031 | 3.152 | 0.0016 | 0.0021 | ** | |
| Threonine | Cell Cycle | Cellular Processes | 0.114 | 0.031 | -3.669 | 2.430e-4 | 3.870e-4 | *** | |
| Threonine | Amino Acid Transport | Metabolism | -0.191 | 0.034 | 5.57 | 2.550e-8 | 8.640e-8 | *** | |
| Threonine | Nucleotide Transport | Metabolism | 0.163 | 0.033 | -4.879 | 1.060e-6 | 2.580e-6 | *** | |
| Threonine | Carbohydrate Transport | Metabolism | -0.104 | 0.036 | 2.92 | 0.0035 | 0.0043 | ** | |
| Threonine | Coenzyme Transport | Metabolism | -0.119 | 0.031 | 3.773 | 1.610e-4 | 2.680e-4 | *** | |
| Threonine | Lipid Transport | Metabolism | 0.043 | 0.032 | -1.331 | 0.183 | 0.185 | | |
| Threonine | Translation | Information Storage | 0.26 | 0.036 | -7.181 | 6.940e-13 | 4.170e-12 | *** | btol limit reached |
| Threonine | Transcription | Information Storage | -0.127 | 0.034 | 3.743 | 1.820e-4 | 3.010e-4 | *** | |
| Threonine | Replication/Repair | Information Storage | 0.102 | 0.027 | -3.722 | 1.970e-4 | 3.210e-4 | *** | |
| Threonine | Cell Wall | Cellular Processes | 0.195 | 0.04 | -4.894 | 9.900e-7 | 2.420e-6 | *** | |
| Threonine | Motility | Cellular Processes | -0.081 | 0.031 | 2.599 | 0.0094 | 0.0108 | * | |
| Threonine | Posttrans. Mod | Cellular Processes | 0.304 | 0.038 | -8.092 | 5.890e-16 | 5.850e-15 | *** | |
| Threonine | Inorganic Ion Transport | Metabolism | -0.134 | 0.031 | 4.259 | 2.060e-5 | 4.030e-5 | *** | |
| Threonine | Secondary Metabolites | Metabolism | -0.129 | 0.036 | 3.572 | 3.550e-4 | 5.320e-4 | *** | |
| Threonine | Signal Transduction | Cellular Processes | -0.145 | 0.031 | 4.711 | 2.460e-6 | 5.500e-6 | *** | |

*Continued on next page*

*... continued*

| Amino acid | COG category | COG class | Log-odds (per 1 SD) | Std. error | Z value | P value | P adj (BH) | Sig | Convergence warning |
|---|---|---|---|---|---|---|---|---|---|
| Threonine | Intracellular Traffic | Cellular Processes | 0.123 | 0.036 | -3.456 | 5.490e-4 | 7.860e-4 | *** | |
| Threonine | Defense | Cellular Processes | 0.105 | 0.033 | -3.236 | 0.0012 | 0.0016 | ** | |
| Glutamine | Energy Production | Metabolism | -0.17 | 0.053 | 3.176 | 0.0015 | 0.0019 | ** | |
| Glutamine | Cell Cycle | Cellular Processes | 0.219 | 0.057 | -3.862 | 1.120e-4 | 1.940e-4 | *** | |
| Glutamine | Amino Acid Transport | Metabolism | -0.28 | 0.061 | 4.625 | 3.740e-6 | 8.100e-6 | *** | |
| Glutamine | Nucleotide Transport | Metabolism | 0.315 | 0.065 | -4.818 | 1.450e-6 | 3.450e-6 | *** | |
| Glutamine | Carbohydrate Transport | Metabolism | -0.136 | 0.054 | 2.516 | 0.0119 | 0.0135 | * | |
| Glutamine | Coenzyme Transport | Metabolism | -0.172 | 0.048 | 3.563 | 3.670e-4 | 5.470e-4 | *** | |
| Glutamine | Lipid Transport | Metabolism | -0.179 | 0.048 | 3.709 | 2.080e-4 | 3.370e-4 | *** | |
| Glutamine | Translation | Information Storage | 0.971 | 0.063 | -15.35 | 3.530e-53 | 5.080e-51 | *** | btol limit reached |
| Glutamine | Transcription | Information Storage | 0.106 | 0.055 | -1.917 | 0.0552 | 0.0589 | | |
| Glutamine | Replication/Repair | Information Storage | 0.455 | 0.058 | -7.839 | 4.530e-15 | 3.840e-14 | *** | |
| Glutamine | Cell Wall | Cellular Processes | 0.257 | 0.071 | -3.609 | 3.080e-4 | 4.740e-4 | *** | |
| Glutamine | Motility | Cellular Processes | -0.129 | 0.048 | 2.704 | 0.0068 | 0.0081 | ** | |
| Glutamine | Posttrans. Mod | Cellular Processes | 0.392 | 0.065 | -6.056 | 1.400e-9 | 5.590e-9 | *** | |
| Glutamine | Inorganic Ion Transport | Metabolism | -0.233 | 0.053 | 4.409 | 1.040e-5 | 2.090e-5 | *** | |
| Glutamine | Secondary Metabolites | Metabolism | -0.212 | 0.059 | 3.63 | 2.840e-4 | 4.420e-4 | *** | |
| Glutamine | Signal Transduction | Cellular Processes | -0.263 | 0.06 | 4.371 | 1.240e-5 | 2.480e-5 | *** | |
| Glutamine | Intracellular Traffic | Cellular Processes | 0.418 | 0.074 | -5.686 | 1.300e-8 | 4.620e-8 | *** | |
| Glutamine | Defense | Cellular Processes | -0.199 | 0.043 | 4.599 | 4.240e-6 | 9.050e-6 | *** | |

Significance: *** $p < 0.001$, ** $p < 0.01$, * $p < 0.05$, · $p < 0.1$. P adj = BH-corrected $p$-value.

**Table S4.** AIC comparison: environment-only versus multivariate model.

| **Amino acid** | **ΔAIC (env-only)** | **ΔAIC (multivariate)** | **ΔΔAIC** |
|---|---:|---:|---:|
| Leucine | 117.45 | -11.43 | -128.88 |
| Glutamine | 151.99 | 31.03 | -120.96 |
| Histidine | 129.35 | 16.58 | -112.78 |
| Proline | 137.83 | 28.05 | -109.78 |
| Tyrosine | 110.2 | 6.74 | -103.46 |
| Cysteine | 93.69 | -2.99 | -96.68 |
| Lysine | 99.99 | 8.71 | -91.27 |
| Tryptophan | 128.19 | 38.78 | -89.42 |
| Phenylalanine | 78.16 | -6.32 | -84.48 |
| Serine | 70.12 | -4.73 | -74.85 |
| Threonine | 79.75 | 12.09 | -67.66 |
| Arginine | 66.9 | 4 | -62.9 |
| Methionine | 58.18 | -4.33 | -62.51 |
| Isoleucine | 50.88 | 0.31 | -50.56 |
| Asparagine | 71.47 | 34.31 | -37.17 |
| Valine | 33.84 | 1.63 | -32.21 |

**Table S5a.** COG fraction x auxotrophy burden: phylolm intercepts (prototrophic baseline %).

| COG category | COG ID | Intercept (%) |
|---|---|---|
| Energy Production | cog_C | 10.141 |
| Cell Cycle | cog_D | 2.178 |
| Amino Acid Transport | cog_E | 10.051 |
| Nucleotide Transport | cog_F | 4.619 |
| Carbohydrate Transport | cog_G | 9.558 |
| Coenzyme Transport | cog_H | 7.085 |
| Lipid Transport | cog_I | 3.4 |
| Translation | cog_J | 9.665 |
| Transcription | cog_K | 8.495 |
| Replication/Repair | cog_L | 7.965 |
| Cell Wall | cog_M | 8.425 |
| Motility | cog_N | 2.1 |
| Posttrans. Mod. | cog_O | 3.886 |
| Inorganic Ion Transport | cog_P | 8.103 |
| Secondary Metabolites | cog_Q | 2.067 |
| Signal Transduction | cog_T | 5.124 |
| Intracellular Traffic | cog_U | 3.196 |
| Defense | cog_V | 2.388 |

**Table S5b.** COG fraction x auxotrophy burden: phylolm coefficients (percentage points) relative to bin 0.

| COG category | COG ID | Bin | Estimate (pp) | Std. error (pp) | P adj (BH) | Sig |
|---|---|---|---|---|---|---|
| Energy Production | cog_C | 1-2 | 0.0078 | 0.0351 | 0.857 | |
| Energy Production | cog_C | 3-4 | 0.071 | 0.0504 | 0.211 | |
| Energy Production | cog_C | 5-6 | 0.0739 | 0.0655 | 0.314 | |
| Energy Production | cog_C | 7-8 | -5e-4 | 0.0812 | 0.995 | |
| Energy Production | cog_C | 9-10 | 0.0736 | 0.0805 | 0.429 | |
| Energy Production | cog_C | 11-12 | 0.1256 | 0.081 | 0.164 | |
| Energy Production | cog_C | 13-14 | 0.0123 | 0.0886 | 0.902 | |
| Cell Cycle | cog_D | 1-2 | 0.0571 | 0.0117 | 3.030e-6 | *** |
| Cell Cycle | cog_D | 3-4 | 0.0755 | 0.0165 | 1.110e-5 | *** |
| Cell Cycle | cog_D | 5-6 | 0.1282 | 0.0214 | 7.670e-9 | *** |
| Cell Cycle | cog_D | 7-8 | 0.1617 | 0.0256 | 1.080e-9 | *** |
| Cell Cycle | cog_D | 9-10 | 0.2105 | 0.0253 | 8.080e-16 | *** |
| Cell Cycle | cog_D | 11-12 | 0.255 | 0.0253 | 1.280e-22 | *** |
| Cell Cycle | cog_D | 13-14 | 0.2645 | 0.0286 | 3.120e-19 | *** |
| Amino Acid Transport | cog_E | 1-2 | -0.0229 | 0.036 | 0.591 | |
| Amino Acid Transport | cog_E | 3-4 | -0.0309 | 0.0518 | 0.614 | |
| Amino Acid Transport | cog_E | 5-6 | -0.2413 | 0.0674 | 6.900e-4 | *** |
| Amino Acid Transport | cog_E | 7-8 | -0.5882 | 0.0837 | 1.160e-11 | *** |
| Amino Acid Transport | cog_E | 9-10 | -0.8109 | 0.083 | 2.190e-21 | *** |
| Amino Acid Transport | cog_E | 11-12 | -0.9784 | 0.0834 | 4.230e-30 | *** |
| Amino Acid Transport | cog_E | 13-14 | -0.9308 | 0.0911 | 3.480e-23 | *** |
| Nucleotide Transport | cog_F | 1-2 | 0.1126 | 0.0177 | 9.020e-10 | *** |
| Nucleotide Transport | cog_F | 3-4 | 0.1756 | 0.0252 | 1.670e-11 | *** |
| Nucleotide Transport | cog_F | 5-6 | 0.2436 | 0.0328 | 6.820e-13 | *** |
| Nucleotide Transport | cog_F | 7-8 | 0.3259 | 0.0399 | 2.430e-15 | *** |
| Nucleotide Transport | cog_F | 9-10 | 0.4179 | 0.0396 | 1.360e-24 | *** |
| Nucleotide Transport | cog_F | 11-12 | 0.4942 | 0.0397 | 1.350e-33 | *** |
| Nucleotide Transport | cog_F | 13-14 | 0.3804 | 0.0441 | 6.440e-17 | *** |
| Carbohydrate Transport | cog_G | 1-2 | -0.0675 | 0.0499 | 0.224 | |

*Continued on next page*

*. . . continued*

| COG category | COG ID | Bin | Estimate (pp) | Std. error (pp) | P adj (BH) | Sig |
|---|---|---|---|---|---|---|
| Carbohydrate Transport | cog_G | 3-4 | -0.0256 | 0.0715 | 0.769 | |
| Carbohydrate Transport | cog_G | 5-6 | -0.4594 | 0.0929 | 2.160e-6 | *** |
| Carbohydrate Transport | cog_G | 7-8 | -0.7925 | 0.1146 | 2.240e-11 | *** |
| Carbohydrate Transport | cog_G | 9-10 | -0.962 | 0.1136 | 2.210e-16 | *** |
| Carbohydrate Transport | cog_G | 11-12 | -0.8606 | 0.1141 | 3.220e-13 | *** |
| Carbohydrate Transport | cog_G | 13-14 | -0.8258 | 0.1255 | 2.150e-10 | *** |
| Coenzyme Transport | cog_H | 1-2 | 0.0677 | 0.0245 | 0.0095 | ** |
| Coenzyme Transport | cog_H | 3-4 | 0.0915 | 0.0349 | 0.0143 | * |
| Coenzyme Transport | cog_H | 5-6 | 0.0399 | 0.0454 | 0.447 | |
| Coenzyme Transport | cog_H | 7-8 | 0.0073 | 0.0555 | 0.902 | |
| Coenzyme Transport | cog_H | 9-10 | 0.0631 | 0.055 | 0.308 | |
| Coenzyme Transport | cog_H | 11-12 | 0.0749 | 0.0552 | 0.224 | |
| Coenzyme Transport | cog_H | 13-14 | 0.0111 | 0.0612 | 0.877 | |
| Lipid Transport | cog_I | 1-2 | 0.0192 | 0.0271 | 0.555 | |
| Lipid Transport | cog_I | 3-4 | 0.0079 | 0.039 | 0.867 | |
| Lipid Transport | cog_I | 5-6 | 0.0703 | 0.0507 | 0.217 | |
| Lipid Transport | cog_I | 7-8 | 0.0282 | 0.0628 | 0.709 | |
| Lipid Transport | cog_I | 9-10 | 0.1258 | 0.0623 | 0.0621 | |
| Lipid Transport | cog_I | 11-12 | 0.139 | 0.0626 | 0.0396 | * |
| Lipid Transport | cog_I | 13-14 | 0.118 | 0.0685 | 0.118 | |
| Translation | cog_J | 1-2 | 0.2383 | 0.0437 | 1.550e-7 | *** |
| Translation | cog_J | 3-4 | 0.4351 | 0.0625 | 1.680e-11 | *** |
| Translation | cog_J | 5-6 | 0.74 | 0.0813 | 9.030e-19 | *** |
| Translation | cog_J | 7-8 | 0.9549 | 0.1 | 1.730e-20 | *** |
| Translation | cog_J | 9-10 | 1.2328 | 0.0991 | 1.370e-33 | *** |
| Translation | cog_J | 11-12 | 1.627 | 0.0995 | 6.650e-57 | *** |
| Translation | cog_J | 13-14 | 1.7914 | 0.1097 | 6.650e-57 | *** |
| Transcription | cog_K | 1-2 | -0.1575 | 0.0426 | 4.600e-4 | *** |
| Transcription | cog_K | 3-4 | -0.3404 | 0.0601 | 4.660e-8 | *** |
| Transcription | cog_K | 5-6 | -0.491 | 0.0781 | 1.290e-9 | *** |

*Continued on next page*

*... continued*

| COG category | COG ID | Bin | Estimate (pp) | Std. error (pp) | P adj (BH) | Sig |
|---|---|---|---|---|---|---|
| Transcription | cog_K | 7-8 | -0.5445 | 0.0938 | 2.270e-8 | *** |
| Transcription | cog_K | 9-10 | -0.6636 | 0.093 | 5.590e-12 | *** |
| Transcription | cog_K | 11-12 | -0.8493 | 0.0929 | 6.980e-19 | *** |
| Transcription | cog_K | 13-14 | -0.6868 | 0.1047 | 2.410e-10 | *** |
| Replication/Repair | cog_L | 1-2 | 0.0226 | 0.0511 | 0.709 | |
| Replication/Repair | cog_L | 3-4 | -0.046 | 0.0707 | 0.585 | |
| Replication/Repair | cog_L | 5-6 | 0.1358 | 0.0921 | 0.188 | |
| Replication/Repair | cog_L | 7-8 | 0.2863 | 0.1079 | 0.0132 | * |
| Replication/Repair | cog_L | 9-10 | 0.3973 | 0.1067 | 4.150e-4 | *** |
| Replication/Repair | cog_L | 11-12 | 0.5575 | 0.1058 | 3.980e-7 | *** |
| Replication/Repair | cog_L | 13-14 | 0.6482 | 0.1216 | 2.900e-7 | *** |
| Cell Wall | cog_M | 1-2 | 0.0576 | 0.0361 | 0.151 | |
| Cell Wall | cog_M | 3-4 | 0.1045 | 0.0512 | 0.0596 | |
| Cell Wall | cog_M | 5-6 | 0.2813 | 0.0665 | 5.270e-5 | *** |
| Cell Wall | cog_M | 7-8 | 0.4317 | 0.0807 | 2.660e-7 | *** |
| Cell Wall | cog_M | 9-10 | 0.4908 | 0.08 | 3.140e-9 | *** |
| Cell Wall | cog_M | 11-12 | 0.4958 | 0.0801 | 2.300e-9 | *** |
| Cell Wall | cog_M | 13-14 | 0.3808 | 0.0895 | 4.810e-5 | *** |
| Motility | cog_N | 1-2 | -0.0564 | 0.03 | 0.0851 | |
| Motility | cog_N | 3-4 | -0.0754 | 0.0429 | 0.111 | |
| Motility | cog_N | 5-6 | -0.0388 | 0.0558 | 0.558 | |
| Motility | cog_N | 7-8 | 0.0164 | 0.0686 | 0.852 | |
| Motility | cog_N | 9-10 | -0.0624 | 0.068 | 0.429 | |
| Motility | cog_N | 11-12 | -0.0799 | 0.0683 | 0.299 | |
| Motility | cog_N | 13-14 | -0.057 | 0.0753 | 0.524 | |
| Posttrans. Mod. | cog_O | 1-2 | 0.0484 | 0.0185 | 0.0144 | * |
| Posttrans. Mod. | cog_O | 3-4 | 0.1138 | 0.026 | 2.800e-5 | *** |
| Posttrans. Mod. | cog_O | 5-6 | 0.1949 | 0.0338 | 2.710e-8 | *** |
| Posttrans. Mod. | cog_O | 7-8 | 0.2962 | 0.0405 | 1.520e-12 | *** |
| Posttrans. Mod. | cog_O | 9-10 | 0.4028 | 0.0401 | 1.630e-22 | *** |

*Continued on next page*

. . . continued

| COG category | COG ID | Bin | Estimate (pp) | Std. error (pp) | P adj (BH) | Sig |
|---|---|---|---|---|---|---|
| Posttrans. Mod. | cog_O | 11-12 | 0.4798 | 0.04 | 2.340e-31 | *** |
| Posttrans. Mod. | cog_O | 13-14 | 0.4655 | 0.0452 | 1.970e-23 | *** |
| Inorganic Ion Transport | cog_P | 1-2 | -0.0394 | 0.0323 | 0.276 | |
| Inorganic Ion Transport | cog_P | 3-4 | -0.0603 | 0.0462 | 0.242 | |
| Inorganic Ion Transport | cog_P | 5-6 | -0.1717 | 0.06 | 0.0072 | ** |
| Inorganic Ion Transport | cog_P | 7-8 | -0.2215 | 0.0738 | 0.0049 | ** |
| Inorganic Ion Transport | cog_P | 9-10 | -0.2574 | 0.0732 | 8.470e-4 | *** |
| Inorganic Ion Transport | cog_P | 11-12 | -0.3052 | 0.0735 | 7.200e-5 | *** |
| Inorganic Ion Transport | cog_P | 13-14 | -0.3144 | 0.081 | 2.240e-4 | *** |
| Secondary Metabolites | cog_Q | 1-2 | -0.0746 | 0.0224 | 0.0016 | ** |
| Secondary Metabolites | cog_Q | 3-4 | -0.0925 | 0.0322 | 0.0072 | ** |
| Secondary Metabolites | cog_Q | 5-6 | -0.1005 | 0.0419 | 0.0252 | * |
| Secondary Metabolites | cog_Q | 7-8 | -0.1483 | 0.0518 | 0.0072 | ** |
| Secondary Metabolites | cog_Q | 9-10 | -0.2191 | 0.0514 | 4.680e-5 | *** |
| Secondary Metabolites | cog_Q | 11-12 | -0.2389 | 0.0517 | 9.660e-6 | *** |
| Secondary Metabolites | cog_Q | 13-14 | -0.1944 | 0.0566 | 0.0011 | ** |
| Signal Transduction | cog_T | 1-2 | -0.1006 | 0.047 | 0.0473 | * |
| Signal Transduction | cog_T | 3-4 | -0.2069 | 0.0679 | 0.0042 | ** |
| Signal Transduction | cog_T | 5-6 | -0.2265 | 0.0884 | 0.0164 | * |
| Signal Transduction | cog_T | 7-8 | -0.265 | 0.1106 | 0.0252 | * |
| Signal Transduction | cog_T | 9-10 | -0.3892 | 0.1096 | 7.590e-4 | *** |
| Signal Transduction | cog_T | 11-12 | -0.7729 | 0.1105 | 1.400e-11 | *** |
| Signal Transduction | cog_T | 13-14 | -0.6887 | 0.1195 | 2.710e-8 | *** |
| Intracellular Traffic | cog_U | 1-2 | -0.0068 | 0.0233 | 0.817 | |
| Intracellular Traffic | cog_U | 3-4 | 0.0146 | 0.0326 | 0.709 | |
| Intracellular Traffic | cog_U | 5-6 | 0.109 | 0.0424 | 0.0163 | * |
| Intracellular Traffic | cog_U | 7-8 | 0.2383 | 0.0504 | 6.110e-6 | *** |
| Intracellular Traffic | cog_U | 9-10 | 0.2343 | 0.0499 | 7.000e-6 | *** |
| Intracellular Traffic | cog_U | 11-12 | 0.2215 | 0.0497 | 2.080e-5 | *** |
| Intracellular Traffic | cog_U | 13-14 | 0.2027 | 0.0566 | 6.900e-4 | *** |

*Continued on next page*

*... continued*

| COG category | COG ID | Bin | Estimate (pp) | Std. error (pp) | P adj (BH) | Sig |
|---|---|---|---|---|---|---|
| Defense | cog_V | 1-2 | -0.0441 | 0.0182 | 0.0242 | * |
| Defense | cog_V | 3-4 | -0.1132 | 0.0256 | 2.350e-5 | *** |
| Defense | cog_V | 5-6 | -0.117 | 0.0333 | 8.470e-4 | *** |
| Defense | cog_V | 7-8 | -0.0232 | 0.0398 | 0.619 | |
| Defense | cog_V | 9-10 | -0.1159 | 0.0394 | 0.0058 | ** |
| Defense | cog_V | 11-12 | -0.053900000000000003 | 0.0393 | 0.221 | |
| Defense | cog_V | 13-14 | -0.0968 | 0.0445 | 0.044 | * |

Significance: *** $p < 0.001$, ** $p < 0.01$, * $p < 0.05$, $\cdot$ $p < 0.1$. P adj = BH-corrected $p$-value.

**Table S6a.** Co-auxotrophy observed vs. expected: regression by environment.

| Group | Pair type | N | Intercept | Slope | Std. error (slope) | $R^2$ | P (slope $\neq$ 1) | Sig |
|---|---|---|---|---|---|---|---|---|
| Global | Doubles | 120 | 0.01836 | 1.9205 | 0.0464 | 0.936 | 2.080e-39 | *** |
| Animal | Doubles | 120 | 0.04674 | 1.4113 | 0.0555 | 0.846 | 2.060e-11 | *** |
| Aquatic | Doubles | 120 | 0.01641 | 2.0597 | 0.0548 | 0.923 | 2.320e-38 | *** |
| Soil | Doubles | 120 | 0.00352 | 2.0155 | 0.0466 | 0.941 | 3.260e-43 | *** |
| Plant | Doubles | 120 | 0.00487 | 3.177 | 0.1236 | 0.848 | 8.010e-35 | *** |
| Global | Triples | 560 | 0.0202 | 4.742 | 0.0644 | 0.907 | 8.810e-239 | *** |
| Animal | Triples | 560 | 0.04294 | 2.978 | 0.0715 | 0.757 | 9.000e-107 | *** |
| Aquatic | Triples | 560 | 0.01832 | 5.4935 | 0.0845 | 0.883 | 1.570e-220 | *** |
| Soil | Triples | 560 | 0.00433 | 5.0831 | 0.0615 | 0.924 | 4.640e-267 | *** |
| Plant | Triples | 560 | 0.00382 | 15.305 | 0.3261 | 0.798 | 5.550e-183 | *** |

$P$ (slope $\neq 1$): two-sided $t$-test against slope = 1.

**Table S6b.** Co-auxotrophy observed vs. expected: regression by phylum.

| **Group** | **Pair type** | **N** | **Intercept** | **Slope** | **Std. error (slope)** | $\mathbf{R^2}$ | **P (slope ≠ 1)** | **Sig** |
|---|---|---|---|---|---|---|---|---|
| Global | Doubles | 120 | 0.01836 | 1.9205 | 0.0464 | 0.936 | 2.080e-39 | *** |
| Proteobacteria | Doubles | 120 | 0.02707 | 2.5048 | 0.1254 | 0.772 | 3.510e-22 | *** |
| Firmicutes_A | Doubles | 120 | 0.03175 | 1.2741 | 0.0509 | 0.842 | 3.660e-7 | *** |
| Bacteroidota | Doubles | 120 | 0.01172 | 1.4642 | 0.0185 | 0.982 | 3.760e-49 | *** |
| Global | Triples | 560 | 0.0202 | 4.742 | 0.0644 | 0.907 | 8.810e-239 | *** |
| Proteobacteria | Triples | 560 | 0.02802 | 10.5797 | 0.3868 | 0.573 | 6.630e-92 | *** |
| Firmicutes_A | Triples | 560 | 0.02917 | 2.6124 | 0.0773 | 0.672 | 8.190e-72 | *** |
| Bacteroidota | Triples | 560 | 0.0156 | 2.3677 | 0.0167 | 0.973 | 4.330e-313 | *** |

$P$ (slope $\neq$ 1): two-sided $t$-test against slope = 1.

**Table S7.** PGLMM auxotrophy identity × burden bin coefficients (log-odds). Bin "0" is the reference level. LRT $\chi^2$ and $p$-values are model-level.

| Amino acid | Term | Estimate (log-odds) | Std. error | LRT $\chi^2$ | LRT P | LRT P adj | Sig |
|---|---|---|---|---|---|---|---|
| Lysine | Intercept (bin 0) | -6.802 | 0.708 | 1697.29 | 0 | 0 | *** |
| Lysine | other_auxos_bin1-2 | 1.758 | 0.764 | 1697.29 | 0 | 0 | *** |
| Lysine | other_auxos_bin3-4 | 3.156 | 0.74 | 1697.29 | 0 | 0 | *** |
| Lysine | other_auxos_bin5-6 | 3.435 | 0.758 | 1697.29 | 0 | 0 | *** |
| Lysine | other_auxos_bin7-8 | 4.212 | 0.739 | 1697.29 | 0 | 0 | *** |
| Lysine | other_auxos_bin9-10 | 5.546 | 0.717 | 1697.29 | 0 | 0 | *** |
| Lysine | other_auxos_bin11-12 | 6.249 | 0.712 | 1697.29 | 0 | 0 | *** |
| Lysine | other_auxos_bin13+ | 7.473 | 0.717 | 1697.29 | 0 | 0 | *** |
| Isoleucine | Intercept (bin 0) | -6.396 | 0.578 | 3885.46 | 0 | 0 | *** |
| Isoleucine | other_auxos_bin1-2 | 4.151 | 0.583 | 3885.46 | 0 | 0 | *** |
| Isoleucine | other_auxos_bin3-4 | 5.836 | 0.582 | 3885.46 | 0 | 0 | *** |
| Isoleucine | other_auxos_bin5-6 | 6.972 | 0.587 | 3885.46 | 0 | 0 | *** |
| Isoleucine | other_auxos_bin7-8 | 8.262 | 0.597 | 3885.46 | 0 | 0 | *** |
| Isoleucine | other_auxos_bin9-10 | 9.276 | 0.608 | 3885.46 | 0 | 0 | *** |
| Isoleucine | other_auxos_bin11-12 | 9.517 | 0.621 | 3885.46 | 0 | 0 | *** |
| Isoleucine | other_auxos_bin13+ | 9.467 | 0.655 | 3885.46 | 0 | 0 | *** |
| Arginine | Intercept (bin 0) | -4.237 | 0.198 | 3326.68 | 0 | 0 | *** |
| Arginine | other_auxos_bin1-2 | 1.388 | 0.222 | 3326.68 | 0 | 0 | *** |
| Arginine | other_auxos_bin3-4 | 2.441 | 0.22 | 3326.68 | 0 | 0 | *** |
| Arginine | other_auxos_bin5-6 | 3.368 | 0.224 | 3326.68 | 0 | 0 | *** |
| Arginine | other_auxos_bin7-8 | 4.471 | 0.223 | 3326.68 | 0 | 0 | *** |
| Arginine | other_auxos_bin9-10 | 5.657 | 0.223 | 3326.68 | 0 | 0 | *** |
| Arginine | other_auxos_bin11-12 | 6.981 | 0.275 | 3326.68 | 0 | 0 | *** |
| Arginine | other_auxos_bin13+ | 7.642 | 0.41 | 3326.68 | 0 | 0 | *** |
| Glutamine | Intercept (bin 0) | -5.549 | 0.379 | 1134.56 | 6.950e-242 | 7.410e-242 | *** |
| Glutamine | other_auxos_bin1-2 | 0.325 | 0.494 | 1134.56 | 6.950e-242 | 7.410e-242 | *** |
| Glutamine | other_auxos_bin3-4 | 1.002 | 0.506 | 1134.56 | 6.950e-242 | 7.410e-242 | *** |
| Glutamine | other_auxos_bin5-6 | 0.902 | 0.629 | 1134.56 | 6.950e-242 | 7.410e-242 | *** |
| Glutamine | other_auxos_bin7-8 | 0.849 | 0.693 | 1134.56 | 6.950e-242 | 7.410e-242 | *** |

*Continued on next page*

*... continued*

| Amino acid | Term | Estimate (log-odds) | Std. error | LRT $\chi^2$ | LRT P | LRT P adj | Sig |
|---|---|---|---|---|---|---|---|
| Glutamine | other__auxos__bin9-10 | 2.551 | 0.446 | 1134.56 | 6.950e-242 | 7.410e-242 | *** |
| Glutamine | other__auxos__bin11-12 | 4.216 | 0.391 | 1134.56 | 6.950e-242 | 7.410e-242 | *** |
| Glutamine | other__auxos__bin13+ | 5.491 | 0.392 | 1134.56 | 6.950e-242 | 7.410e-242 | *** |
| Serine | Intercept (bin 0) | -3.401 | 0.131 | 167.88 | 1.260e-33 | 1.260e-33 | *** |
| Serine | other__auxos__bin1-2 | 0.728 | 0.162 | 167.88 | 1.260e-33 | 1.260e-33 | *** |
| Serine | other__auxos__bin3-4 | 1.145 | 0.176 | 167.88 | 1.260e-33 | 1.260e-33 | *** |
| Serine | other__auxos__bin5-6 | 0.778 | 0.239 | 167.88 | 1.260e-33 | 1.260e-33 | *** |
| Serine | other__auxos__bin7-8 | 0.593 | 0.27 | 167.88 | 1.260e-33 | 1.260e-33 | *** |
| Serine | other__auxos__bin9-10 | 0.964 | 0.226 | 167.88 | 1.260e-33 | 1.260e-33 | *** |
| Serine | other__auxos__bin11-12 | 1.046 | 0.197 | 167.88 | 1.260e-33 | 1.260e-33 | *** |
| Serine | other__auxos__bin13+ | 2.293 | 0.171 | 167.88 | 1.260e-33 | 1.260e-33 | *** |
| Methionine | Intercept (bin 0) | -2.092 | 0.071 | 2021.82 | 0 | 0 | *** |
| Methionine | other__auxos__bin1-2 | 0.612 | 0.094 | 2021.82 | 0 | 0 | *** |
| Methionine | other__auxos__bin3-4 | 1.017 | 0.108 | 2021.82 | 0 | 0 | *** |
| Methionine | other__auxos__bin5-6 | 1.863 | 0.12 | 2021.82 | 0 | 0 | *** |
| Methionine | other__auxos__bin7-8 | 2.638 | 0.131 | 2021.82 | 0 | 0 | *** |
| Methionine | other__auxos__bin9-10 | 3.738 | 0.132 | 2021.82 | 0 | 0 | *** |
| Methionine | other__auxos__bin11-12 | 4.98 | 0.217 | 2021.82 | 0 | 0 | *** |
| Methionine | other__auxos__bin13+ | 5.179 | 0.316 | 2021.82 | 0 | 0 | *** |
| Tryptophan | Intercept (bin 0) | -1.778 | 0.062 | 1885.62 | 0 | 0 | *** |
| Tryptophan | other__auxos__bin1-2 | 0.741 | 0.082 | 1885.62 | 0 | 0 | *** |
| Tryptophan | other__auxos__bin3-4 | 1.646 | 0.096 | 1885.62 | 0 | 0 | *** |
| Tryptophan | other__auxos__bin5-6 | 1.81 | 0.121 | 1885.62 | 0 | 0 | *** |
| Tryptophan | other__auxos__bin7-8 | 2.467 | 0.124 | 1885.62 | 0 | 0 | *** |
| Tryptophan | other__auxos__bin9-10 | 4.021 | 0.153 | 1885.62 | 0 | 0 | *** |
| Tryptophan | other__auxos__bin11-12 | 6.135 | 0.416 | 1885.62 | 0 | 0 | *** |
| Phenylalanine | Intercept (bin 0) | -3.052 | 0.111 | 3128.26 | 0 | 0 | *** |
| Phenylalanine | other__auxos__bin1-2 | 0.855 | 0.135 | 3128.26 | 0 | 0 | *** |
| Phenylalanine | other__auxos__bin3-4 | 1.537 | 0.143 | 3128.26 | 0 | 0 | *** |
| Phenylalanine | other__auxos__bin5-6 | 2.561 | 0.149 | 3128.26 | 0 | 0 | *** |

*Continued on next page*

*... continued*

| Amino acid | Term | Estimate (log-odds) | Std. error | LRT $\chi^2$ | LRT P | LRT P adj | Sig |
|---|---|---|---|---|---|---|---|
| Phenylalanine | other_auxos_bin7-8 | 3.742 | 0.155 | 3128.26 | 0 | 0 | *** |
| Phenylalanine | other_auxos_bin9-10 | 5.06 | 0.169 | 3128.26 | 0 | 0 | *** |
| Phenylalanine | other_auxos_bin11-12 | 6.353 | 0.271 | 3128.26 | 0 | 0 | *** |
| Phenylalanine | other_auxos_bin13+ | 7.848 | 0.719 | 3128.26 | 0 | 0 | *** |
| Tyrosine | Intercept (bin 0) | -3.688 | 0.151 | 3034.3 | 0 | 0 | *** |
| Tyrosine | other_auxos_bin1-2 | 1.329 | 0.172 | 3034.3 | 0 | 0 | *** |
| Tyrosine | other_auxos_bin3-4 | 2.082 | 0.177 | 3034.3 | 0 | 0 | *** |
| Tyrosine | other_auxos_bin5-6 | 2.866 | 0.184 | 3034.3 | 0 | 0 | *** |
| Tyrosine | other_auxos_bin7-8 | 3.874 | 0.183 | 3034.3 | 0 | 0 | *** |
| Tyrosine | other_auxos_bin9-10 | 5.172 | 0.184 | 3034.3 | 0 | 0 | *** |
| Tyrosine | other_auxos_bin11-12 | 6.143 | 0.225 | 3034.3 | 0 | 0 | *** |
| Tyrosine | other_auxos_bin13+ | 7.791 | 0.526 | 3034.3 | 0 | 0 | *** |
| Cysteine | Intercept (bin 0) | -4.404 | 0.215 | 1475.95 | 8.660e-300 | 1.070e-298 | *** |
| Cysteine | other_auxos_bin1-2 | 0.789 | 0.259 | 1475.95 | 8.660e-300 | 1.070e-298 | *** |
| Cysteine | other_auxos_bin3-4 | 1.972 | 0.248 | 1475.95 | 8.660e-300 | 1.070e-298 | *** |
| Cysteine | other_auxos_bin5-6 | 2.461 | 0.261 | 1475.95 | 8.660e-300 | 1.070e-298 | *** |
| Cysteine | other_auxos_bin7-8 | 2.573 | 0.267 | 1475.95 | 8.660e-300 | 1.070e-298 | *** |
| Cysteine | other_auxos_bin9-10 | 3.207 | 0.244 | 1475.95 | 8.660e-300 | 1.070e-298 | *** |
| Cysteine | other_auxos_bin11-12 | 3.953 | 0.228 | 1475.95 | 8.660e-300 | 1.070e-298 | *** |
| Cysteine | other_auxos_bin13+ | 5.476 | 0.252 | 1475.95 | 8.660e-300 | 1.070e-298 | *** |
| Leucine | Intercept (bin 0) | -4.551 | 0.231 | 3900.43 | 0 | 0 | *** |
| Leucine | other_auxos_bin1-2 | 1.491 | 0.256 | 3900.43 | 0 | 0 | *** |
| Leucine | other_auxos_bin3-4 | 3.288 | 0.244 | 3900.43 | 0 | 0 | *** |
| Leucine | other_auxos_bin5-6 | 4.614 | 0.249 | 3900.43 | 0 | 0 | *** |
| Leucine | other_auxos_bin7-8 | 5.977 | 0.265 | 3900.43 | 0 | 0 | *** |
| Leucine | other_auxos_bin9-10 | 7.11 | 0.283 | 3900.43 | 0 | 0 | *** |
| Leucine | other_auxos_bin11-12 | 7.044 | 0.287 | 3900.43 | 0 | 0 | *** |
| Leucine | other_auxos_bin13+ | 8.941 | 0.625 | 3900.43 | 0 | 0 | *** |
| Histidine | Intercept (bin 0) | -2.768 | 0.097 | 2904.37 | 0 | 0 | *** |
| Histidine | other_auxos_bin1-2 | 1.014 | 0.117 | 2904.37 | 0 | 0 | *** |

*Continued on next page*

*. . . continued*

| Amino acid | Term | Estimate (log-odds) | Std. error | LRT $\chi^2$ | LRT P | LRT P adj | Sig |
|---|---|---|---|---|---|---|---|
| Histidine | other_auxos_bin3-4 | 1.635 | 0.126 | 2904.37 | 0 | 0 | *** |
| Histidine | other_auxos_bin5-6 | 2.46 | 0.14 | 2904.37 | 0 | 0 | *** |
| Histidine | other_auxos_bin7-8 | 3.513 | 0.144 | 2904.37 | 0 | 0 | *** |
| Histidine | other_auxos_bin9-10 | 5.487 | 0.2 | 2904.37 | 0 | 0 | *** |
| Histidine | other_auxos_bin11-12 | 6.429 | 0.308 | 2904.37 | 0 | 0 | *** |
| Histidine | other_auxos_bin13+ | 8.257 | 1.007 | 2904.37 | 0 | 0 | *** |
| Proline | Intercept (bin 0) | -2.659 | 0.092 | 2141.95 | 0 | 0 | *** |
| Proline | other_auxos_bin1-2 | 1.128 | 0.11 | 2141.95 | 0 | 0 | *** |
| Proline | other_auxos_bin3-4 | 1.946 | 0.118 | 2141.95 | 0 | 0 | *** |
| Proline | other_auxos_bin5-6 | 2.608 | 0.136 | 2141.95 | 0 | 0 | *** |
| Proline | other_auxos_bin7-8 | 3.328 | 0.143 | 2141.95 | 0 | 0 | *** |
| Proline | other_auxos_bin9-10 | 4.176 | 0.143 | 2141.95 | 0 | 0 | *** |
| Proline | other_auxos_bin11-12 | 4.99 | 0.183 | 2141.95 | 0 | 0 | *** |
| Proline | other_auxos_bin13+ | 5.545 | 0.3 | 2141.95 | 0 | 0 | *** |
| Asparagine | Intercept (bin 0) | -4.605 | 0.237 | 1307.58 | 2.480e-279 | 2.830e-279 | *** |
| Asparagine | other_auxos_bin1-2 | 0.267 | 0.314 | 1307.58 | 2.480e-279 | 2.830e-279 | *** |
| Asparagine | other_auxos_bin3-4 | 0.868 | 0.328 | 1307.58 | 2.480e-279 | 2.830e-279 | *** |
| Asparagine | other_auxos_bin5-6 | 1.436 | 0.343 | 1307.58 | 2.480e-279 | 2.830e-279 | *** |
| Asparagine | other_auxos_bin7-8 | 1.981 | 0.321 | 1307.58 | 2.480e-279 | 2.830e-279 | *** |
| Asparagine | other_auxos_bin9-10 | 2.707 | 0.278 | 1307.58 | 2.480e-279 | 2.830e-279 | *** |
| Asparagine | other_auxos_bin11-12 | 3.831 | 0.251 | 1307.58 | 2.480e-279 | 2.830e-279 | *** |
| Asparagine | other_auxos_bin13+ | 4.973 | 0.262 | 1307.58 | 2.480e-279 | 2.830e-279 | *** |
| Valine | Intercept (bin 0) | -5.703 | 0.409 | 3126.63 | 0 | 0 | *** |
| Valine | other_auxos_bin1-2 | 3.567 | 0.415 | 3126.63 | 0 | 0 | *** |
| Valine | other_auxos_bin3-4 | 5.269 | 0.415 | 3126.63 | 0 | 0 | *** |
| Valine | other_auxos_bin5-6 | 6.396 | 0.423 | 3126.63 | 0 | 0 | *** |
| Valine | other_auxos_bin7-8 | 7.768 | 0.441 | 3126.63 | 0 | 0 | *** |
| Valine | other_auxos_bin9-10 | 8.833 | 0.461 | 3126.63 | 0 | 0 | *** |
| Valine | other_auxos_bin11-12 | 7.652 | 0.434 | 3126.63 | 0 | 0 | *** |
| Valine | other_auxos_bin13+ | 5.544 | 0.426 | 3126.63 | 0 | 0 | *** |

*Continued on next page*

*... continued*

| Amino acid | Term | Estimate (log-odds) | Std. error | LRT $\chi^2$ | LRT P | LRT P adj | Sig |
|---|---|---|---|---|---|---|---|
| Threonine | Intercept (bin 0) | -2.822 | 0.1 | 2653.94 | 0 | 0 | *** |
| Threonine | other__auxos__bin1-2 | 1.046 | 0.119 | 2653.94 | 0 | 0 | *** |
| Threonine | other__auxos__bin3-4 | 1.57 | 0.13 | 2653.94 | 0 | 0 | *** |
| Threonine | other__auxos__bin5-6 | 2.581 | 0.138 | 2653.94 | 0 | 0 | *** |
| Threonine | other__auxos__bin7-8 | 3.785 | 0.154 | 2653.94 | 0 | 0 | *** |
| Threonine | other__auxos__bin9-10 | 4.864 | 0.164 | 2653.94 | 0 | 0 | *** |
| Threonine | other__auxos__bin11-12 | 5.521 | 0.213 | 2653.94 | 0 | 0 | *** |
| Threonine | other__auxos__bin13+ | 5.591 | 0.284 | 2653.94 | 0 | 0 | *** |

Significance: *** $p < 0.001$, ** $p < 0.01$, * $p < 0.05$, · $p < 0.1$. P adj = BH-corrected $p$-value.

**Table S8a.** PGLMM enrichment profiles: crossover summary. $\Delta$AIC: full bin model vs intercept-only PGLMM. Enrichment fold-range: ratio of maximum to minimum predicted loss probability. Crossover bin: first bin where predicted probability exceeds global auxotrophy prevalence.

| Amino acid | delta_AIC | Enrichment fold-range | Crossover bin | Crossover mid. | Burden pattern |
|---|---|---|---|---|---|
| L-Isoleucine | 4895.9 | 575.27 | 3-4 | 3.5 | moderate-burden |
| L-Serine | 185.6 | 7.69 | 3-4 | 3.5 | moderate-burden |
| L-Tryptophan | 2407.1 | 6.83 | 3-4 | 3.5 | moderate-burden |
| L-Valine | 4207.7 | 288.23 | 3-4 | 3.5 | moderate-burden |
| L-Arginine | 4023.4 | 67.94 | 5-6 | 5.5 | mid-burden |
| L-Methionine | 2809.7 | 8.71 | 5-6 | 5.5 | mid-burden |
| L-Phenylalanine | 3920.9 | 21.98 | 5-6 | 5.5 | mid-burden |
| L-Tyrosine | 3702.9 | 40.31 | 5-6 | 5.5 | mid-burden |
| L-Leucine | 4820.7 | 94.51 | 5-6 | 5.5 | mid-burden |
| L-Histidine | 3793.3 | 16.85 | 5-6 | 5.5 | mid-burden |
| L-Proline | 2948.7 | 14.47 | 5-6 | 5.5 | mid-burden |
| L-Threonine | 3479.6 | 16.76 | 5-6 | 5.5 | mid-burden |
| L-Cysteine | 1537.2 | 61.67 | 7-8 | 7.5 | mid-burden |
| L-Lysine | 1748.5 | 595.84 | 9-10 | 9.5 | high-burden |
| L-Asparagine | 1304.5 | 59.66 | 9-10 | 9.5 | high-burden |
| L-Glutamine | 1124.4 | 125.26 | 11-12 | 11.5 | extreme-burden |

**Table S8b.** PGLMM enrichment profiles: full $\log_2$ enrichment table.

| Amino acid | Bin | Pred. prob. | Global prev. | Enrichment ratio | Log2 enrichment |
|---|---|---|---|---|---|
| Lysine | 1 | 0.0017 | 0.1324 | 0.013 | -6.269 |
| Lysine | 2-3 | 0.0101 | 0.1324 | 0.0764 | -3.7098 |
| Lysine | 4-5 | 0.036 | 0.1324 | 0.272 | -1.8785 |
| Lysine | 6-7 | 0.0405 | 0.1324 | 0.3056 | -1.7102 |
| Lysine | 8-9 | 0.0676 | 0.1324 | 0.5106 | -0.9696 |
| Lysine | 10-11 | 0.1724 | 0.1324 | 1.3018 | 0.3805 |
| Lysine | 12-15 | 0.6498 | 0.1324 | 4.9081 | 2.2952 |
| Isoleucine | 1 | 0.0026 | 0.4714 | 0.0055 | -7.5163 |
| Isoleucine | 2-3 | 0.1619 | 0.4714 | 0.3434 | -1.5421 |
| Isoleucine | 4-5 | 0.5139 | 0.4714 | 1.0901 | 0.1244 |
| Isoleucine | 6-7 | 0.7399 | 0.4714 | 1.5694 | 0.6502 |
| Isoleucine | 8-9 | 0.9099 | 0.4714 | 1.9299 | 0.9486 |
| Isoleucine | 10-11 | 0.9587 | 0.4714 | 2.0336 | 1.024 |
| Isoleucine | 12-15 | 0.9705 | 0.4714 | 2.0585 | 1.0416 |
| Arginine | 1 | 0.0223 | 0.3619 | 0.0617 | -4.0192 |
| Arginine | 2-3 | 0.0877 | 0.3619 | 0.2423 | -2.0449 |
| Arginine | 4-5 | 0.1997 | 0.3619 | 0.5518 | -0.8578 |
| Arginine | 6-7 | 0.3757 | 0.3619 | 1.0383 | 0.0542 |
| Arginine | 8-9 | 0.5944 | 0.3619 | 1.6425 | 0.7159 |
| Arginine | 10-11 | 0.8689 | 0.3619 | 2.4013 | 1.2638 |
| Arginine | 12-15 | 0.9733 | 0.3619 | 2.6897 | 1.4274 |
| Glutamine | 1 | 0.006 | 0.0805 | 0.0746 | -3.7443 |
| Glutamine | 2-3 | 0.0084 | 0.0805 | 0.1047 | -3.2555 |
| Glutamine | 4-5 | 0.0147 | 0.0805 | 0.1829 | -2.4507 |
| Glutamine | 6-7 | 0.0116 | 0.0805 | 0.1436 | -2.8002 |
| Glutamine | 8-9 | 0.0085 | 0.0805 | 0.1049 | -3.2523 |
| Glutamine | 10-11 | 0.0341 | 0.0805 | 0.4236 | -1.2392 |
| Glutamine | 12-15 | 0.4543 | 0.0805 | 5.6416 | 2.4961 |
| Serine | 1 | 0.0515 | 0.106 | 0.4861 | -1.0407 |
| Serine | 2-3 | 0.1012 | 0.106 | 0.955 | -0.0665 |
| Serine | 4-5 | 0.1309 | 0.106 | 1.2358 | 0.3054 |
| Serine | 6-7 | 0.078 | 0.106 | 0.7365 | -0.4412 |
| Serine | 8-9 | 0.0535 | 0.106 | 0.5051 | -0.9853 |
| Serine | 10-11 | 0.0575 | 0.106 | 0.5422 | -0.883 |
| Serine | 12-15 | 0.2293 | 0.106 | 2.1637 | 1.1135 |
| Methionine | 1 | 0.1906 | 0.4827 | 0.3947 | -1.341 |
| Methionine | 2-3 | 0.2799 | 0.4827 | 0.5799 | -0.7862 |
| Methionine | 4-5 | 0.3355 | 0.4827 | 0.695 | -0.5249 |
| Methionine | 6-7 | 0.5491 | 0.4827 | 1.1375 | 0.1859 |
| Methionine | 8-9 | 0.6366 | 0.4827 | 1.3188 | 0.3992 |
| Methionine | 10-11 | 0.9031 | 0.4827 | 1.8707 | 0.9035 |

*Continued on next page*

... continued

| Amino acid | Bin | Pred. prob. | Global prev. | Enrichment ratio | Log2 enrichment |
|---|---|---|---|---|---|
| Methionine | 12-15 | 0.9705 | 0.4827 | 2.0103 | 1.0074 |
| Tryptophan | 1 | 0.2609 | 0.5742 | 0.4545 | -1.1377 |
| Tryptophan | 2-3 | 0.3997 | 0.5742 | 0.6961 | -0.5227 |
| Tryptophan | 4-5 | 0.5696 | 0.5742 | 0.992 | -0.0116 |
| Tryptophan | 6-7 | 0.5462 | 0.5742 | 0.9514 | -0.0719 |
| Tryptophan | 8-9 | 0.7296 | 0.5742 | 1.2707 | 0.3456 |
| Tryptophan | 10-11 | 0.9641 | 0.5742 | 1.6791 | 0.7477 |
| Tryptophan | 12-15 | 0.9986 | 0.5742 | 1.7392 | 0.7984 |
| Phenylalanine | 1 | 0.073 | 0.4224 | 0.1727 | -2.5334 |
| Phenylalanine | 2-3 | 0.1577 | 0.4224 | 0.3733 | -1.4217 |
| Phenylalanine | 4-5 | 0.2488 | 0.4224 | 0.5889 | -0.7638 |
| Phenylalanine | 6-7 | 0.474 | 0.4224 | 1.1221 | 0.1662 |
| Phenylalanine | 8-9 | 0.7239 | 0.4224 | 1.7139 | 0.7773 |
| Phenylalanine | 10-11 | 0.9354 | 0.4224 | 2.2144 | 1.1469 |
| Phenylalanine | 12-15 | 0.9887 | 0.4224 | 2.3408 | 1.227 |
| Tyrosine | 1 | 0.0386 | 0.382 | 0.1011 | -3.3061 |
| Tyrosine | 2-3 | 0.1383 | 0.382 | 0.362 | -1.4661 |
| Tyrosine | 4-5 | 0.2308 | 0.382 | 0.604 | -0.7273 |
| Tyrosine | 6-7 | 0.3786 | 0.382 | 0.991 | -0.013 |
| Tyrosine | 8-9 | 0.5831 | 0.382 | 1.5263 | 0.61 |
| Tyrosine | 10-11 | 0.8707 | 0.382 | 2.2792 | 1.1885 |
| Tyrosine | 12-15 | 0.9789 | 0.382 | 2.5623 | 1.3574 |
| Cysteine | 1 | 0.0189 | 0.1734 | 0.1089 | -3.1985 |
| Cysteine | 2-3 | 0.0413 | 0.1734 | 0.2383 | -2.069 |
| Cysteine | 4-5 | 0.1146 | 0.1734 | 0.6609 | -0.5976 |
| Cysteine | 6-7 | 0.1503 | 0.1734 | 0.8669 | -0.206 |
| Cysteine | 8-9 | 0.1296 | 0.1734 | 0.7474 | -0.42 |
| Cysteine | 10-11 | 0.1741 | 0.1734 | 1.0045 | 0.0065 |
| Cysteine | 12-15 | 0.7018 | 0.1734 | 4.0484 | 2.0173 |
| Leucine | 1 | 0.0163 | 0.4174 | 0.0391 | -4.6775 |
| Leucine | 2-3 | 0.0717 | 0.4174 | 0.1717 | -2.5418 |
| Leucine | 4-5 | 0.3273 | 0.4174 | 0.7843 | -0.3505 |
| Leucine | 6-7 | 0.6618 | 0.4174 | 1.5858 | 0.6652 |
| Leucine | 8-9 | 0.8563 | 0.4174 | 2.0518 | 1.0369 |
| Leucine | 10-11 | 0.9282 | 0.4174 | 2.224 | 1.1531 |
| Leucine | 12-15 | 0.9705 | 0.4174 | 2.3253 | 1.2174 |
| Histidine | 1 | 0.097 | 0.4654 | 0.2084 | -2.2624 |
| Histidine | 2-3 | 0.2344 | 0.4654 | 0.5037 | -0.9895 |
| Histidine | 4-5 | 0.3322 | 0.4654 | 0.7139 | -0.4862 |
| Histidine | 6-7 | 0.4971 | 0.4654 | 1.0682 | 0.0951 |
| Histidine | 8-9 | 0.7775 | 0.4654 | 1.6706 | 0.7403 |
| Histidine | 10-11 | 0.9533 | 0.4654 | 2.0484 | 1.0345 |

*Continued on next page*

*. . . continued*

| **Amino acid** | **Bin** | **Pred. prob.** | **Global prev.** | **Enrichment ratio** | **Log2 enrichment** |
|---|---|---|---|---|---|
| Histidine | 12-15 | 0.9972 | 0.4654 | 2.1427 | 1.0994 |
| Proline | 1 | 0.1082 | 0.4747 | 0.2279 | -2.1338 |
| Proline | 2-3 | 0.2841 | 0.4747 | 0.5986 | -0.7403 |
| Proline | 4-5 | 0.4452 | 0.4747 | 0.9379 | -0.0926 |
| Proline | 6-7 | 0.5607 | 0.4747 | 1.1812 | 0.2403 |
| Proline | 8-9 | 0.6986 | 0.4747 | 1.4717 | 0.5575 |
| Proline | 10-11 | 0.8348 | 0.4747 | 1.7588 | 0.8146 |
| Proline | 12-15 | 0.9648 | 0.4747 | 2.0326 | 1.0234 |
| Asparagine | 1 | 0.0155 | 0.1199 | 0.1289 | -2.9558 |
| Asparagine | 2-3 | 0.0202 | 0.1199 | 0.1688 | -2.5666 |
| Asparagine | 4-5 | 0.0327 | 0.1199 | 0.2731 | -1.8728 |
| Asparagine | 6-7 | 0.0491 | 0.1199 | 0.4099 | -1.2868 |
| Asparagine | 8-9 | 0.0648 | 0.1199 | 0.5404 | -0.8878 |
| Asparagine | 10-11 | 0.0969 | 0.1199 | 0.8087 | -0.3063 |
| Asparagine | 12-15 | 0.5823 | 0.1199 | 4.8572 | 2.2801 |
| Valine | 1 | 0.0052 | 0.4484 | 0.0115 | -6.4441 |
| Valine | 2-3 | 0.1796 | 0.4484 | 0.4005 | -1.3202 |
| Valine | 4-5 | 0.5516 | 0.4484 | 1.23 | 0.2986 |
| Valine | 6-7 | 0.7688 | 0.4484 | 1.7144 | 0.7777 |
| Valine | 8-9 | 0.9324 | 0.4484 | 2.0792 | 1.056 |
| Valine | 10-11 | 0.9443 | 0.4484 | 2.1059 | 1.0744 |
| Valine | 12-15 | 0.73 | 0.4484 | 1.6278 | 0.7029 |
| Threonine | 1 | 0.0918 | 0.4533 | 0.2026 | -2.3031 |
| Threonine | 2-3 | 0.2302 | 0.4533 | 0.5078 | -0.9776 |
| Threonine | 4-5 | 0.2962 | 0.4533 | 0.6535 | -0.6137 |
| Threonine | 6-7 | 0.5723 | 0.4533 | 1.2625 | 0.3363 |
| Threonine | 8-9 | 0.738 | 0.4533 | 1.6282 | 0.7033 |
| Threonine | 10-11 | 0.9264 | 0.4533 | 2.0438 | 1.0312 |
| Threonine | 12-15 | 0.9634 | 0.4533 | 2.1255 | 1.0878 |

### Supplementary Discussion

#### Supplementary Discussion 1. Low-auxotrophy amino acids: transamination and biosynthetic cost

Alanine, aspartate, and glutamate — the three amino acids with the lowest predicted auxotrophy rates — are synthesized via a single transamination step from the central metabolic intermediates pyruvate, oxaloacetate, and $\alpha$-ketoglutarate, respectively (Price et al. 2020). These reactions are catalyzed by aminotransferases with broad substrate ranges that are deeply embedded in nitrogen assimilation and redistribution — functions that must be maintained regardless of amino acid availability in the environment. Because the enzymes responsible are not dedicated biosynthetic machinery but pleiotropic nitrogen-handling catalysts, the cell cannot lose the biosynthetic capacity for these amino acids without simultaneously compromising core nitrogen metabolism. This functional constraint makes auxotrophy for alanine, aspartate, and glutamate selectively costly in a way that is qualitatively different from auxotrophies affecting dedicated biosynthetic pathways.

Compounding this, the ATP investment required to synthesize these three amino acids is among the lowest of any proteinogenic amino acid (Akashi and Gojobori 2002). The selective benefit of avoiding their synthesis — and thus the fitness advantage that could offset the cost of maintaining auxotrophy — is therefore minimal compared with energetically expensive amino acids such as tryptophan or histidine.

---

#### Supplementary Discussion 2. Per-amino-acid auxotrophy rates: biosynthetic cost and pathway entanglement

Auxotrophy rates varied substantially across amino acids (Figure 1b). Amino acids with consistently low auxotrophy rates — glutamine, asparagine, serine, cysteine, and lysine — share either low biosynthetic costs or deep entanglement with essential cellular processes.

**Glutamine and asparagine** are synthesized in single ATP-dependent steps from glutamate and aspartate respectively, representing among the lowest biosynthetic costs of any amino acid (Akashi and Gojobori 2002; Kaleta et al. 2013). Beyond their proteomic role, glutamine synthetase acts as a critical regulatory node in nitrogen metabolism, integrating signals from multiple sensing pathways and directly regulating downstream gene expression (Reitzer 2003, 2004; Yuan et al. 2009). The dual role of this enzyme — biosynthetic and regulatory — means that auxotrophy for glutamine may impose costs beyond the loss of the amino acid itself.

**Serine** is energetically inexpensive to synthesize and serves as the obligate precursor for both glycine and cysteine biosynthesis (Akashi and Gojobori 2002). Loss of serine biosynthesis would therefore necessitate simultaneous exogenous acquisition of multiple downstream metabolites, potentially raising the effective cost of auxotrophy.

**Cysteine** biosynthesis is coupled to sulfur assimilation: the pathway incorporates inorganic sulfate into organic form, meaning that cysteine auxotrophy requires either an exogenous organic sulfur source or specialized adaptations in sulfur metabolism (Fernandez et al. 2025). This dependency may constrain the environments in which cysteine auxotrophy is viable.

**Lysine** biosynthesis, though requiring substantial energy investment and multiple enzymatic steps, proceeds through the diaminopimelate (DAP) pathway, which is also required for peptidoglycan synthesis in most bacteria (Hutton et al. 2007; Kaleta et al. 2013; Akashi and Gojobori 2002). Loss of this pathway therefore simultaneously eliminates lysine biosynthesis and cell wall synthesis capacity, imposing a severe pleiotropic cost that likely constrains the evolution of lysine auxotrophy to lineages that have undergone radical cell wall remodelling.

In contrast, the 11 amino acids with higher auxotrophy rates are synthesized via longer biosynthetic routes — requiring more enzymatic steps and greater phosphate bond investment — and with fewer overlaps with essential pathways. This holds even for amino acids with critical downstream metabolic roles: **methionine**, for instance, is the precursor to S-adenosylmethionine (SAM), the universal methyl donor, yet shows high predicted auxotrophy rates (Fontecave et al. 2004). Unlike glutamine synthetase or the DAP pathway, the methionine biosynthetic pathway is not known to perform additional cellular functions — a cell can synthesize SAM from exogenously acquired methionine, meaning auxotrophy carries no inherent penalty provided methionine is available.

---

#### Supplementary Discussion 3. MAG quality controls for the auxotrophy rate difference

To ensure that the higher auxotrophy rates observed in MAGs compared to non-MAG genomes (Figure 1c) reflect a biological difference rather than a technical artefact of lower assembly quality, we performed two quality control analyses.

First, we compared auxotrophy rates of near-complete genomes from each group (98%–100% completeness; Supplementary Figure 6). The higher auxotrophy rates in MAGs persisted even at this stringent quality threshold, ruling out systematic incompleteness as the primary driver of the observed gap.

Second, we examined whether auxotrophy counts increase with genome fragmentation in MAGs (Supplementary Figure 5). Auxotrophy counts did not increase with greater fragmentation; in fact, higher auxotrophy rates were associated with higher-quality, low-contig assemblies. This indicates that our metabolic reconstructions are not prone to false-positive auxotrophy predictions caused by assembly gaps, as increased fragmentation did not result in the artificial 'loss' of amino acid biosynthetic pathways.

Third, we validated gapseq's ability to recover auxotrophy predictions from incomplete genomes directly, using simulated MAGs from five synthetic communities constructed with CAMISIM. Metagenome-assembled genomes were reconstructed from simulated reads, and auxotrophy predictions were generated using the same gapseq pipeline applied to the main dataset. Predictions for high-quality MAGs ($\geq$85% completeness; n = 14 bins) were compared against experimentally confirmed auxotrophies for the corresponding reference strains (Supplementary Table S1b). For these bins, gapseq achieved sensitivity 0.895 and specificity 0.928, comparable to the isolate benchmark (sensitivity 0.78, specificity 0.98; Supplementary Script S01). The modest reduction in specificity relative to the isolate benchmark is consistent with contaminating sequences in co-assembled bins providing spurious pathway genes. Together, these results confirm that gapseq accurately recovers auxotrophy from MAGs meeting standard quality thresholds.

Together, these controls indicate that the elevated auxotrophy rates in MAGs are not an artefact of assembly completeness or fragmentation, but instead likely reflect the biological underrepresentation of auxotrophic bacteria in culture collections.

---

#### Supplementary Discussion 4. Phylogenetic correction: illustrative cases

Phylogenetic correction altered the estimated effect of environment on auxotrophy in ways that were neither uniform across amino acids nor consistent across environments, reflecting a complex relationship between habitat-specific selection and phylogenetic inertia. Two cases illustrate the range of effects.

**Attenuation of an apparent enrichment.** For animal-associated asparagine auxotrophy, the uncorrected GLM suggested a 2.9-fold increase in odds relative to other environments. After phylogenetic correction, this reduced to a non-significant 1.2-fold (adjusted $p = 0.326$). The shift reflects two simultaneous effects: down-weighting of phylogenetically clustered auxotrophs within the animal niche reduced the estimated probability of auxotrophy from 14.9% to 10.9%, while correction for redundant prototrophic lineages in other environments raised the grand mean probability from 7.1% to 10.2%. The apparent animal enrichment was therefore largely an artefact of phylogenetic overrepresentation of auxotrophic lineages in that niche rather than a recurring independent signal.

**Unmasking of a convergent signal.** For aquatic lysine auxotrophy, the uncorrected model suggested a non-significant 1.56-fold increase in odds. Phylogenetic correction more than doubled this estimate to 3.16-fold (adjusted $p = 2.37 \times 10^{-6}$), driven by an increase in estimated auxotrophy probability from 7.9% to 31%, far exceeding the mean shift from 7.2% to 18.6%. This pattern suggests that lysine auxotrophy has arisen independently across multiple distantly related aquatic lineages — a signal of convergent evolution that was obscured in the uncorrected analysis by the phylogenetic structure of the dataset.

Together these examples illustrate that phylogenetic correction can both increase and decrease estimated effect sizes and alter their statistical significance, depending on whether phylogenetic structure inflates or masks the true ecological signal (Supplementary Figure 2).

---

#### Supplementary Discussion 5. Mechanistic scenarios for sequential auxotrophy accumulation

The consistent priority ordering of auxotrophy losses — rare auxotrophies restricted to genomes already carrying many other losses — is consistent with sequential accumulation. Two non-mutually-exclusive mechanisms could produce this pattern.

**Genomic cascade.** The loss of one biosynthetic pathway may alter nutrient requirements, metabolic flux, network redundancy, or regulatory architecture in ways that render additional pathways dispensable — making each successive loss more likely given the losses that preceded it. Ordered gene loss and progressive genomic degradation have been extensively documented in obligate endosymbionts, where biosynthetic pathways are lost in a structured sequence over evolutionary time (Moran et al. 2008; Wernegreen 2002). The auxotrophies observed here may represent an early or intermediate stage of this same process, in which environmental bacteria have begun the trajectory of genome reduction without yet committing to the obligate dependence characteristic of endosymbionts. The conditions permitting rare auxotrophies are therefore potentially only met once a sufficient number of prior losses have already reshaped the metabolic landscape.

**Ecological feedback loop.** An initial auxotrophy does not merely change the genome — it changes where the organism can live. A bacterium that loses tryptophan synthesis may become more competitive in nutrient-rich environments where tryptophan is reliably supplied, and those same environments tend to supply other amino acids as well. Deeper ecological integration into these environments then progressively relaxes selection on additional biosynthetic pathways, driving further auxotrophy. In this model, auxotrophy leads to niche change, which leads to more auxotrophy.

Both scenarios share the prediction that genomes with low and moderate auxotrophy burdens represent organisms mid-trajectory, still accumulating losses on the pathway of genome reduction.

A third possibility — rapid, simultaneous loss of multiple pathways following a single environmental shift — would predict that intermediate-burden genomes are rare and that rare auxotrophies are not intrinsically late losses. The observation that rare auxotrophies are consistently underrepresented at low and intermediate burden counts argues against this scenario and in favour of sequential accumulation, but cannot definitively exclude it.

Distinguishing between the genomic cascade and ecological feedback scenarios would require tracing the order of auxotrophy acquisition across phylogenetic lineages — specifically, whether late-crossover amino acids are lost after lifestyle transitions or whether lifestyle transitions follow their loss.

---

#### Supplementary Discussion 6. Why habitat labels explain limited auxotrophy variance

Two factors likely decouple present-day habitat classification from the selective pressures that actually shape auxotrophy.

First, environmental classes are imprecise proxies for metabolic reality. Chemical variation within a single environment type can be substantial — dissolved free amino acid concentrations in marine systems span two orders of magnitude (Amaral et al. 2024; Niu et al. 2023) — potentially diluting environment-specific signals. Additionally, very different nominal environments may exhibit similar metabolite landscapes: a streambed downstream of a wastewater outflow may chemically resemble the gut far more than a glacial stream, yet both would be classified as aquatic.

Second, environmental metadata identifies where a genome was sampled, not the selective pressures that drove trait fixation. A lineage recently dispersed from a nutrient-rich host environment into an aquatic system may still harbour auxotrophies fixed under ancestral conditions. Because trait fixation operates on evolutionary timescales while dispersal and habitat transition can be rapid, present-day habitat labels and present-day selective pressures can be substantially misaligned. Consequently, even highly refined environment labels cannot recover signals that are fundamentally encoded in evolutionary history rather than present-day ecology.

---

#### Supplementary Discussion 7. Genome size sensitivity analysis

Genome size is a downstream consequence of streamlining — it co-varies with auxotrophy burden and COG composition rather than causing them independently. Adding genome size as a covariate therefore risks over-adjustment by blocking part of the causal pathway from streamlining to the outcomes being modelled. Nevertheless, we re-fitted both the COG-burden models (script 06) and the auxotrophy identity models (script 08) with log(Genome_Size) as an additional fixed-effect covariate to assess whether the reported signals are carried entirely by genome size variation.

For the COG-burden models, COG fractions are proportions (percentage of annotated genes in each category), which inherently normalise for total gene count and genome size. Adding log genome size nonetheless modified the highest-burden-bin coefficients (Pearson $r = 0.596$ between original and size-adjusted estimates; Supplementary Figure 17), as expected given that genome size and COG composition co-vary under streamlining. Critically, the genome size coefficient varied in sign across COG categories (range: $-0.891$ to $+0.719$), inconsistent with a simple dilution effect in which smaller genomes would have uniformly lower fractions of all categories.

For the auxotrophy identity models, the $\log_2$ enrichment profiles were highly stable when genome size was added as a covariate (Pearson $r = 0.955$; mean $|\Delta\log_2 \text{ enrichment}| = 0.54$; Supplementary Figure 18). The priority ordering and crossover structure were preserved across all amino acids, confirming that these patterns reflect structured, ordered pathway loss rather than undifferentiated gene loss associated with overall genome reduction.
